# Evaluating the impact and detectability of mass extinctions on total-evidence dating

**DOI:** 10.1101/2025.09.28.679059

**Authors:** Minghao Du, Wenhui Wang, Jingqiang Tan, Joëlle Barido-Sottani

## Abstract

Fossils are crucial for accurately dating phylogenetic trees because their ages provide vital constraints on the timing of macroevolutionary events, and their morphological characters offer key information on evolutionary rates and phylogenetic positions. The fossilized birth-death (FBD) process is a diversification model that incorporates both extant and extinct species, serving as a tree prior that seamlessly integrates fossils into phylogenetic inference. While the FBD model can account for mass extinctions, which caused rapid, widespread organismal loss, few studies have utilized FBD models incorporating these events in phylogenetic inference. This is likely because the detectability of mass extinctions and their impact on phylogenetic inference remain unclear. Through simulations, we assessed the influence of mass extinctions on divergence time and topology inference and evaluated the detectability of mass extinction signals in total-evidence dating. We examined three FBD tree priors: without mass extinction, with known mass extinction time and survival probability, and with known mass extinction time but unknown survival probability. Our results show that the FBD model with known mass extinction time and unknown survival probability was able to reliably detect mass extinctions when they occurred, and correctly refrained from detecting mass extinctions when they were absent. Moreover different FBD models generate similar divergence time and tree topology errors. Even when the FBD model used for tree inference did not explicitly account for mass extinction events, signals of mass extinction were still detectable on the resulting MCC trees. The accuracy of the detection was similar to the one obtained from MCC trees inferred using an FBD model that includes mass extinction parameters. We also reduced the fossilization rate and the number of morphological characters, obtaining results consistent with the aforementioned findings. However, reducing the fossilization rate decreased the accuracy of detecting mass extinctions when they occurred, and reducing the number of morphological characters decreased the accuracy of divergence time inference. Furthermore, we adjusted the priors for the existence of mass extinction and the survival probability of mass extinction. We found that the prior for the existence of mass extinction had no effect on inference, whereas the prior for the survival probability of mass extinction significantly influenced both the detection of mass extinctions and the estimation of survival probabilities. Finally, we applied these models to empirical datasets of tetraodontiform fishes and crinoids and found that, consistent with our simulation results, the inclusion of a mass extinction event in the tree prior had a negligible impact on the inferred topologies and divergence times.

## 1 Introduction

Fossil records indicate that several mass extinction events have occurred throughout geological history, leading to the rapid disappearance of numerous species within relatively short timeframes (Raup and Sepkoski, 1982; Jablonski, 2005; Bambach, 2006; Fan et al., 2020; Marshall, 2023). However, there is considerable debate regarding the definition of a mass extinction, including the proportion of organisms that must perish and the timescale over which the event occurs, as different mass extinctions exhibit distinct patterns (Bambach, 2006; Marshall, 2023). For example, the end-Permian mass extinction, known for its rapidity, is estimated to have eliminated approximately 80% of species in around 60,000 years (Burgess et al., 2014; Fan et al., 2020), while the Cretaceous-Paleogene event caused the extinction of all non-avian dinosaurs in approximately 32,000 years (Renne et al., 2013). In contrast, the Frasnian–Famennian (Late Devonian) extinction was a more protracted event, potentially lasting more than 20 million years (Fan et al., 2020). The duration of a mass extinction is often closely linked to its ultimate cause; for instance, the rapid Cretaceous-Paleogene and end-Permian events are considered catastrophic, triggered by a bolide impact (Renne et al., 2013) and Large Igneous Province eruptions (Burgess et al., 2014; Algeo and Shen, 2024), respectively, which led to massive carbon release. In contrast, the more protracted Frasnian–Famennian extinction is considered a bioevolution-triggered carbon-burial event, where the expansion of land plants led to increased nutrient runoff, oceanic anoxia, and carbon burial, resulting in long-term climatic cooling (Algeo and Shen, 2024). The mass extinctions modeled in this study refer to the former type—rapid, catastrophic events with a duration of less than one million years. These events significantly impact the patterns of biodiversity evolution (Sepkoski, 1984) and the shape of phylogenetic trees when the diversification rates vary among different lineages or when the mass extinction is selective (Heard and Mooers, 2002).

The birth-death process—a continuous-time Markov model that describes speciation and extinction through specific birth and death rates—is fundamental in phylogenetic inference for reconstructing evolutionary relationships among species (Stadler, 2013; Morlon, 2014; Morlon et al., 2024). This framework can be extended to incorporate mass extinctions, modeling them as instantaneous events where each contemporaneous lineage has a fixed probability of survival, analogous to the sampling of extant lineages at the present (Stadler, 2011a; Höhna, 2015; MacPherson et al., 2022). Just as each lineage surviving to the present is included in the reconstructed phylogeny with probability *ρ*, so too can a survival probability be assigned to all lineages in the complete phylogeny at a given time point, where the collective loss of these lineages constitutes a mass extinction event (Figure S1). This catastrophic loss of lineages is modeled as a Dirac delta function, i.e. an instantaneous spike in the survival probability, acting in addition to the background extinction rate. Simulations have shown that this method effectively detects mass extinctions from phylogenies of extant taxa (May et al., 2016; Culshaw et al., 2019) and those including fossils (Magee and Höhna, 2021; Didier and Laurin, 2024), provided the phylogenetic tree is sufficiently large, the survival probability is low, and the extinction time is neither too close to the origin nor the present. This approach has been applied to various taxonomic groups, including Tetraodontiform fishes (Arcila and Tyler, 2017), geckos (Brennan and Oliver, 2017), crocodilians (Magee and Höhna, 2021), and angiosperms (Thompson and Ramírez-Barahona, 2023). However, these simulation and empirical studies typically rely on a known, fixed phylogeny to detect mass extinctions. In simulations, mass extinctions are explicitly introduced, ensuring their presence in the phylogeny. In contrast, empirical studies often do not account for mass extinctions during phylogenetic inference. For instance, Arcila and Tyler (2017) used a constant-rate fossilized birth-death (FBD) process model without incorporating mass extinction for phylogenetic reconstruction, then detected mass extinctions based on the inferred tree of extant taxa. This approach raises questions about the accuracy of estimated topologies and divergence times from phylogenies derived under the assumption of no mass extinction, and whether signals of mass extinction remain detectable in such trees.

Bayesian phylogenetic inference simultaneously estimates phylogenetic relationships, substitution rates, clock models, and diversification rates using an integrated tripartite (substitution, clock, and tree) model (Warnock and Wright, 2020). Consequently, neglecting to account for mass extinction events within the tree model could result in inaccurate phylogenetic reconstructions. To our knowledge, the impact of mass extinctions on phylogenetic inference has not been thoroughly investigated. In this manuscript, we focus on total-evidence phylogenetic inference, which uses morphological information from fossils to determine their positions within phylogenetic trees (Zhang et al., 2016; Gavryushkina et al., 2017). This method offers significant advantages over node dating and topological constraint approaches by more effectively accounting for uncertainties in fossil placement within phylogenies, leading to more accurate evolutionary inferences (Barido-Sottani et al., 2023). Additionally, total-evidence dating eliminates the need for subjective prior calibration times for fossils, thereby avoiding biases associated with prior selection (Warnock et al., 2012; Budd and Mann, 2023). Furthermore, by directly incorporating extinct lineages into the tree structure, this approach provides a more robust framework for estimating extinction rates—an inference that previous studies have shown to be unreliable from phylogenies of extant species (Mitchell et al., 2019)—and detecting mass extinctions than methods that only use fossils as node calibrations. The FBD model (Stadler, 2010; Wright et al., 2022) incorporates fossil sampling rates into the traditional birth-death process (Kendall, 1948), providing tree priors that seamlessly integrate fossils into phylogenetic inference. An FBD model that includes mass extinction can serve as a natural prior, embedding mass extinction events within the framework of phylogenetic inference.

Our study aimed to address three key questions: (1) Can mass extinctions be reliably detected in total-evidence dating when the mass extinction time is known? (2) Does using an FBD model without mass extinction reduce the accuracy of estimating divergence times and topologies compared to a model that incorporates mass extinction? (3) In the presence of mass extinction, can these events still be detected in inferred trees when using an FBD model without mass extinction for phylogenetic inference? To answer these questions, we conducted simulations using FBD models both with and without mass extinction, generating data for fossils and extant species. We simulated morphological characters for fossils and both morphological and molecular characters for extant species. Using these data, we performed phylogenetic inference through total-evidence dating, employing three different FBD models as tree priors: (1) a constant FBD model without mass extinction, (2) a constant FBD model including mass extinction with fixed (true) extinction time and survival probability, and (3) a constant FBD model with mass extinction, fixed (true) extinction time, but unknown survival probability. We compared the accuracy of phylogenetic inferences derived from these tree priors and used the FBD model with mass extinction to detect such events in the inferred phylogenetic trees.

## 2 Methods

### 2.1 Simulation

To reduce computational demands and enhance the accuracy of model inference (excluding mass extinction), we opted for simple models (i.e., those with fewer parameters). This strategy allows for a clearer comparison of the impact of mass extinction. The parameters for these simulations were chosen to approximate common macroevolutionary scenarios while maintaining computational tractability.

Our simulation study was designed to assess the performance of total-evidence dating under a variety of conditions. We generated datasets across a range of scenarios by varying four key factors: (1) the presence or absence of a mass extinction event, (2) the fossil sampling rate (high vs. low), (3) the number of morphological characters (high vs. low), and (4) the models of character evolution, contrasting simple versus complex scenarios (involving both the clock and substitution models) for molecular and morphological data. The following sections detail the specific parameters used for each component of the simulation.

#### 2.1.1 Tree Simulation

Phylogenetic trees were simulated using the R package `TreeSim` (Stadler, 2011b). Each simulation included a fixed number of 50 extant species with a sampling probability of 1. We simulated trees to have a fixed number of extant species, rather than the same origin time, to ensure that the amount of information from the molecular characters is comparable across the different simulation scenarios. This also aligns with the methodology of many previous simulation studies (e.g., Barido-Sottani et al. 2023). We applied constant speciation and extinction rates of 0.15 and 0.1, respectively. The simulation ends at the present (time = 0Ma). For scenarios involving mass extinction, we introduced a mass extinction event at 30 million years before the end of the simulation (30Ma), with a survival probability of 0.2—meaning each species had an 80% chance of extinction. This timing was selected to ensure the mass extinction event was neither too close to the origin nor the present, facilitating effective detection as suggested by (May et al., 2016). The low survival probability was chosen to amplify the extinction signal. We excluded trees with origin times less than 45 Ma or greater than 160 Ma to mitigate the influence of outliers. We discuss this filtering criteria in more details in Section S3. Additionally, we simulated trees without mass extinction, excluding those with origin times less than 34 Ma or greater than 125 Ma. Since these simulations were conditioned on a fixed number of extant species, trees without mass extinction events had shorter origin times compared to those with mass extinctions.

#### 2.1.2 Fossil Simulation

Fossils were simulated using the R package `FossilSim` (Barido-Sottani et al., 2019) on the complete phylogeny. We implemented two fossil sampling scenarios: a high sampling rate (*ψ* = 0.03) and a low sampling rate (*ψ* = 0.015). For subsequent inferences in `RevBayes`, fossil age ranges were set to [true fossil age *×* 0.999, true fossil age *×* 1.001]. To reduce the influence of outliers, we excluded trees with fewer than 21 or more than 100 fossils for high sampling, and those with fewer than 9 or more than 51 fossils for low sampling, in scenarios involving mass extinction. In the absence of mass extinction, trees with fewer than 14 or more than 52 fossils were excluded under high sampling, and those with fewer than 6 or more than 28 fossils were excluded under low sampling. We discuss this filtering criteria in more details in Section S3. The number of fossils was lower in non-extinction scenarios due to shorter origin times. A total of 400 trees were generated, including 100 trees for each combination of high and low sampling rates under both mass extinction and non-mass extinction scenarios.

#### 2.1.3 Molecular Character Simulation

We simulated 1,000-site molecular sequence alignments using the R package `phyclust` (Chen et al., 2023). Data were generated under two distinct evolutionary models, each pairing a clock model with a specific nucleotide substitution model.

First, we simulated data under a strict molecular clock, where the substitution rate was fixed at 0.01 substitutions per site per million years across all branches. These simulations used the homogeneous JC69 model of nucleotide substitution. This first scenario represents a very simplified situation with little uncertainty in the alignment.

Second, to model rate heterogeneity among lineages and between sites, we used an uncorrelated lognormal (UCLN) relaxed clock combined with the HKY+Γ substitution model (*α* = 0.25, five discrete gamma rate categories). The rate for each branch was drawn through a hierarchical process. Initially, a mean evolutionary rate for the tree was drawn from a gamma distribution, Γ(1, 0.005). Subsequently, this mean was used to parameterize a lognormal distribution, Lognormal(mean rate, 0.2^2^), from which a unique rate was drawn independently for each branch. This second scenario is designed to represent a more realistic dataset.

Given the variation in total tree length across our simulations, we supplement the substitution rate with additional summary statistics of the simulated molecular characters. For datasets generated under both strict and relaxed clock models, we report the mean and median of the expected number of substitutions per site, as well as the number of variable sites (Section S1).

#### 2.1.4 Morphological Character Simulation

To examine the effect of character quantity on our analyses, we simulated two sets of binary morphological characters: one with 25 characters (*c* = 25) and another with 250 (*c* = 250). Data were generated under two distinct evolutionary models, each pairing a clock model with a specific character state substitution model.

First, we simulated data under a strict morphological clock, where the substitution rate was fixed at 0.01 substitutions per site per million years across all branches. These simulations used a homogeneous 2-state Mk model (Lewis, 2001) and were performed using the `sim.char` function from the R package `geiger` (Pennell et al., 2014).

Second, to model rate heterogeneity across both lineages and characters, we used a UCLN relaxed clock combined with an Mk+Γ model. These binary data were generated by first simulating four-state data for each character under the JC69 model using the R package `phyclust` (Chen et al., 2023). The resulting nucleotide data were then recoded to a binary format by merging A and G into state 0, and C and T into state 1. The branch-specific rates for the UCLN clock were determined through a hierarchical process: a mean clock rate for the tree was drawn from a gamma distribution, 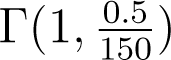, and this mean was then used to parameterize a lognormal distribution, Lognormal(mean rate, 0.2^2^), from which a unique rate was drawn independently for each branch. Similar to the molecular alignments, these two scenarios are designed to represent respectively an ideal situation with little uncertainty, and a more realistic dataset with larger variations.

### 2.2 Inference on the Simulated Data

Phylogenetic inference was performed using the total-evidence dating method implemented in `RevBayes` (Höhna et al., 2016). To ensure model accuracy (excluding mass extinction parameters) and optimize computational efficiency, we selected models and priors that mirrored our simulation conditions (details are provided in the `RevBayes` code). To ensure computational feasibility across our extensive simulations, a single MCMC chain was run for each dataset. We discarded 20% of the chain from the strict clock analyses and 10% from the UCLN clock analyses as burn-in and used the R package `coda` (Plummer et al., 2006) to calculate effective sample sizes (ESS). Since the number of iterations required to meet this threshold varied for each simulated tree, this resulted in different MCMC chain lengths for each simulated tree. Inferences with ESS below 200 (Table S1), even after extended runs, were excluded from further analysis. Non-convergence issues varied across datasets, impacting between 1 and 19 out of 100 replicates.

#### 2.2.1 Tree Inference

We employed constant-rate FBD models as tree priors, each with exponential priors: speciation and extinction rates had an exponential prior with a rate of 10, and the fossilization rate had an exponential prior with a rate of 30. The age of each fossil was assigned a uniform prior distribution, spanning the narrow uncertainty range established during our simulation (see Section 2.1.2). Given the narrowness of these priors, the fossil ages were effectively treated as true values with negligible uncertainty.

For the inference of mass extinction events, we opted not to use an FBD model that simultaneously infers both the timing and survival probability. This decision was informed by a preliminary analysis of 20 simulated trees containing mass extinction events (see Section S4), which revealed that such an approach frequently led to the erroneous detection of multiple extinction events. Instead, we assumed a known mass extinction time to estimate the corresponding survival probability. This assumption of a known mass extinction time is reasonable, as we can rely on paleobiological and paleoenvironmental knowledge to identify periods likely associated with mass extinctions, such as the recognized timings of the famous big five mass extinction events in the Phanerozoic. For comparison, we also used two alternative models: an FBD model with no mass extinction, and one in which both the time and survival probability of mass extinction were fixed to the true simulated values. Therefore, the three FBD models used in our analysis were:

1. FBD model without mass extinction.
2. FBD model with a fixed mass extinction time of 30 Ma and a fixed survival probability of 0.2.
3. FBD model with a fixed mass extinction time of 30 Ma, while treating the survival probability as an unknown parameter. To test for the presence of a mass extinction, we used a reversible-jump mixture model framework. This approach allows the MCMC simulation to jump between two competing models: one that includes a mass extinction and one that does not. The prior was assigned an equal weight of 0.5 to each of the two models (prior=0.5): (i) a state representing no mass extinction, where the survival probability of mass extinction is fixed at exactly 1 (a point mass), and (ii) a state representing the presence of a mass extinction, where the survival probability is an unknown parameter drawn from a prior distribution. For the latter, we assigned a uniform prior distribution from 0.0 to 0.5 for the survival probability. Additionally, we examined the effect of varying the prior weights for the absence of mass extinction by adjusting the weights to 0.1 (favored the presence of a mass extinction, prior=0.1) and 0.9 (favored the absence of a mass extinction, prior=0.9). To assess the influence of different prior assumptions for the survival probability of mass extinction, we also replaced the flat prior Uniform(0.0, 0.5) with a stronger prior Beta(*α* = 10*, β* = 40), which has a mean of 0.2 and a 95% confidence interval ranging from approximately 0.102 to 0.320.

#### 2.2.2 Substitution Inference

For molecular characters, we used a substitution model which matched the model used for the dataset, i.e. the JC69 and HKY+Γ substitution model. Similarly, for morphological characters, we used the two-state Mk and Mk+Γ model matching the simulation model.

#### 2.2.3 Clock Inference

Consistent with our simulations, we inferred evolutionary rates using two distinct frameworks: a strict clock and a UCLN relaxed clock. In both approaches, separate clock models were applied to the molecular and morphological datasets to allow them to have different underlying evolutionary dynamics.

Under the strict clock framework, we assigned the exponential prior distribution with a rate of 100 for both the molecular and morphological clock rates.

Under the UCLN relaxed clock framework, for the morphological data, the hyperprior for the mean of the rate distribution was an exponential with a rate of 100, and for the standard deviation, an exponential with a rate of 5. For the molecular data, the hyperprior for the mean rate was an exponential with a rate of 500, while the hyperprior for the standard deviation was an exponential with a rate of 5.

#### 2.2.4 Inference in MCC Tree

We sought to test whether the signal of mass extinction could still be detected on trees that were inferred using different FBD models as tree priors. To do this, we conducted a two-step analysis where we first inferred maximum clade credibility (MCC) trees and then estimated mass extinction and diversification rates on these fixed trees using `RevBayes` (Figure S2B). The use of MCC trees was a necessary and appropriate summary of the posterior distribution due to the computational expense of analyzing all posterior trees. The MCC trees used in this analysis were themselves inferred from a representative simulation scenario characterized by a high sampling rate of 0.03, 250 morphological characters, a strict clock, and the presence of a mass extinction event. These trees were generated using two distinct FBD priors: one assuming no mass extinctions and another estimating the survival probability. Our choice of this single scenario is justified by the high degree of similarity in topology and divergence times observed across all simulation scenarios between trees inferred under these two FBD priors. With the trees fixed, we were able to efficiently infer both the timing and survival probability of mass extinctions simultaneously.

Speciation, extinction, and fossilization rates were treated as constant throughout the tree, following the same prior distributions as in the total-evidence dating analysis: speciation and extinction rates had an exponential prior with a rate of 10, and the fossilization rate had an exponential prior with a rate of 30. We established a set of discrete, potential mass extinction times. These potential mass extinction times were defined at 1-Myr intervals from the present (time 0) to the rounded root age (e.g., at 1 Ma, 2 Ma, 3 Ma, etc.). We assigned a prior probability of a mass extinction event to each potential time point, calculated as *p* = 0.5*/N*, where *N* is the total number of potential mass extinction times. The prior distribution for survival probability in each mass extinction was specified as Uniform(0.0, 0.5).

### 2.3 Evaluation Metrics

#### 2.3.1 Detectability of Mass Extinction in Total-Evidence Dating

Following the approach of May et al. (2016), we employed Bayes factors to test the hypothesis of a mass extinction at a specific time *t* (30 Ma in our simulations). We compared two competing models: one that included a mass extinction event at time *t* (BF_ME_) and one that did not (BF_WME_). The Bayes factor quantifies the relative evidence for these models provided by the data, and is defined as the ratio of their marginal likelihoods:

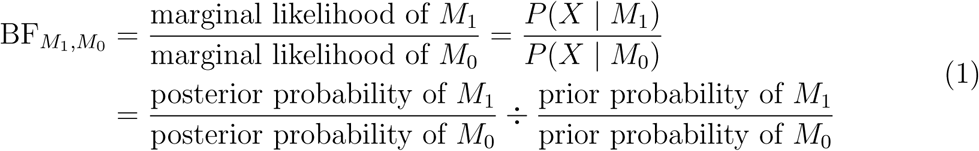

where *X* represents data. The posterior odds of a mass extinction are determined by the proportion of posterior samples that support the event (i.e., those where the survival probability is less than 1.0). The BF_ME_,_WME_, however, is calculated from the ratio of the posterior odds to the prior odds, and is therefore influenced by the prior probability for the existence of mass extinction. For instance, if the prior probability for the existence of mass extinction is set to 0.5, the prior odds are 1 (i.e., 0.5*/*(1 *−* 0.5)). In this scenario, a 2 log BF_ME_,_WME_ = 0 is obtained when the posterior probability of a mass extinction is also 0.5, which corresponds to 50% of posterior samples supporting the mass extinction, yielding posterior odds of 1. Conversely, if a strong prior probability of 0.9 for the existence of mass extinction is used, the prior odds are 9 (i.e., 0.9*/*(1 *−* 0.9)). Consequently, a 2 log BF_ME_,_WME_ = 0 is obtained only when the posterior probability reaches 0.9, which corresponds to 90% of posterior samples supporting the mass extinction, at which point the posterior odds also equal 9, matching the prior odds. A detailed derivation of how the Bayes factor was calculated from the MCMC output is provided in the Section S2. We employed the reversible jump method rather than alternative approaches like the stepping-stone algorithm (Xie et al., 2011) to compute marginal likelihoods. For some analyses, all posterior samples supported the mass extinction model, resulting in an infinite Bayes factor. To facilitate visualization, these cases were assigned a maximum 2 log BF_ME_,_WME_ = 0 value of 20.

#### 2.3.2 Divergence Time

Similar to Barido-Sottani et al. (2023), we assessed the accuracy of divergence time estimation by calculating the absolute relative error of divergence times (the absolute difference between the mean estimate and the true value, divided by the true value, averaged across all nodes in the extant phylogeny), and the 95% highest posterior density (HPD) coverage for the MCC extant tree compared to the true extant tree. To compute the relative error, we excluded nodes where the true divergence time was less than 1 Ma, which accounted for 13.6–14.9% of the data, as these cases exhibited excessively high relative errors that could skew results (for example, a true divergence time of 0.01 Ma with inferred divergence time of 1 Ma results in a relative error of 99).

We categorized the true divergence times into 1 million-year intervals (e.g., 29.5–30.5 Ma) and calculated the average relative error of divergence time estimates for each interval across all simulated phylogenetic trees. This “by time” approach was specifically chosen to assess whether the magnitude of divergence time error was elevated in periods close to the simulated mass extinction event (30 Ma).

#### 2.3.3 Tree Topology

Similar to Barido-Sottani et al. (2023), we measured the accuracy of tree topology by calculating the average normalized Robinson-Foulds (RF) distance between all posterior trees and the true extant tree using the R package `phangorn` Schliep et al. (2024). We focused primarily on the topology and divergence times of the extant trees for two main reasons. First, the phylogenetic relationships among extant species are the central focus of many evolutionary studies, while the phylogenetic placement of fossil species and sampled ancestors is subject to uncertainty. Second, previous simulation studies have shown that inferences of the full tree topology are generally less accurate than for the extant-only subtree (e.g., Barido-Sottani et al. 2023). However, we have also provided the comparisons of both topology and divergence times for the reconstructed MCC tree including fossils (Figure S3).

#### 2.3.4 Detectability of Mass Extinction in MCC Tree

To assess whether mass extinction events were detectable in the MCC trees, we used Bayes factors as a criterion for determining the presence of mass extinction.

When using the criterion 2 log BF_ME_,_WME_ *>* 6 for mass extinction, we define the time with the highest Bayes Factor as the inferred extinction time, following the approach of May et al. (2016). The potential mass extinction interval is then defined as the contiguous block of time points surrounding and including the inferred mass extinction time, where all points within this block have a 2 log BF_ME_,_WME_ *>* 0. The mass extinction survival probability is then calculated as the average of the survival probabilities for these intervals. If, within an MCMC iteration, two or more extinction intervals had survival probabilities less than 1—indicating simultaneous mass extinction events over a short period—we calculated the overall mass extinction survival probability as the product of the survival probabilities from each individual interval. This approach treats these pulses as a sequence of events and correctly calculates the cumulative probability of a lineage surviving the entire series. Additionally, the number of such iterations was usually small, minimizing the impact on overall results.

#### 2.3.5 Diversification Rates

Following the approach used for divergence time estimation (see Section 2.3.2), we evaluated the accuracy of speciation, extinction, and sampling rates inferred through total-evidence dating and on the MCC tree by assessing coverage.

### 2.4 Empirical Data Analysis

We evaluated the impact of tree priors on phylogenetic inference using two empirical datasets associated with historical mass extinction events: one comprising tetraodontiform fishes (Arcila and Tyler, 2017) and another of crinoids (Cole et al., 2025). To investigate the effect of a mass extinction, we performed three parallel sets of analyses for each dataset, each employing a different tree prior:

1. An FBD model without mass extinction.
2. Model 2: An FBD model with a fixed mass extinction time. The prior probability for the absence of the mass extinction was set to 0.5 (corresponding to the third model in Section 2.2.1, prior=0.5), and the survival probability was inferred.
3. An FBD model with a fixed mass extinction time. The prior probability for the absence of the mass extinction was set to 0, i.e., the event was assumed to be a certainty (corresponding to the third model in Section 2.2.1, prior=0; or the second model in Section 2.2.1, but with an unknown survival probability), and only the survival probability was inferred.

For the specific models and priors, please see Sections S5 and S6.

## 3 Results

### 3.1 Inference of Mass Extinctions in Total-Evidence Dating

#### 3.1.1 Bayes Factor

Using an FBD model with fixed mass extinction time and estimated survival probabilities as a tree prior in total-evidence dating allows for accurate detection of mass extinction events when they occurred. Moreover, in the absence of mass extinction, the model did not falsely identify such events. Under a strict clock, and with a threshold of 2 log BF_ME_,_WME_ *>* 0, a prior probability of 0.5 for the existence of mass extinction, and flat prior for the survival probability of mass extinction, the accuracy of detecting mass extinction when present was 95.6%, while the accuracy of correctly identifying its absence was 89.2% (Table 1, Figure 1). However, when a stricter threshold of 2 log BF_ME_,_WME_ *>* 6 was applied, the detection accuracy dropped to 73.2% (Table 1). Similarly, with a stricter threshold of 2 log BF_ME_,_WME_ *< −*6, the accuracy of confirming the absence of mass extinction decreased to 18.6% (Table 1). Total-evidence dating showed higher accuracy in detecting mass extinctions when they occurred than in confirming their absence when they did not (Table 1, Figure 1). Specifically, when a mass extinction occurred, the magnitude of 2 log BF_ME_,_WME_ *>* 0 was significantly greater than the magnitude of 2 log BF_ME_,_WME_ *<* 0 in cases where mass extinction was absent (Figure 1). These findings were robust to the choice of clock model. We observed similar patterns of detection accuracy and asymmetry in analyses conducted under the relaxed clock model (Table 1, Figure 1).

**Table 1:** Accuracy rate of detecting mass extinction (ME) and absence of mass extinction (WME) events using various 2 log BF_ME,WME_ thresholds. For the definitions of “ME”, <u>“WME”, “</u>*<u>c</u>*<u>”, “</u>*<u>ψ</u>*<u>”, “prior”, “strong prior”, and “relaxed clock”, please refer to</u> Figure 1.

| Model | $2 \log \text{BF}$<br>$< -6$ | $2 \log \text{BF}$<br>$< -2$ | $2 \log \text{BF}$<br>$< 0$ | $2 \log \text{BF}$<br>$> 0$ | $2 \log \text{BF}$<br>$> 2$ | $2 \log \text{BF}$<br>$> 6$ |
| --- | --- | --- | --- | --- | --- | --- |
| ME, $c=250$, $\psi=0.03$,<br>prior=0.1 | 0 | 0.011 | 0.056 | 0.944 | 0.900 | 0.722 |
| ME, $c=250$, $\psi=0.03$,<br>prior=0.9 | 0 | 0.011 | 0.054 | 0.946 | 0.924 | 0.707 |
| ME, $c=250$, $\psi=0.03$,<br>prior=0.5 | 0 | 0.011 | 0.044 | 0.956 | 0.933 | 0.722 |
| ME, $c=250$, $\psi=0.015$,<br>prior=0.5 | 0 | 0.032 | 0.064 | 0.936 | 0.819 | 0.638 |
| ME, $c=25$, $\psi=0.03$,<br>prior=0.5 | 1 | 0 | 0.022 | 0.077 | 0.923 | 0.681 |
| ME, $c=250$, $\psi=0.03$,<br>prior=0.5, strong prior | 0 | 0.033 | 0.066 | 0.934 | 0.912 | 0.736 |
| ME, $c=250$, $\psi=0.03$,<br>prior=0.5, relaxed clock | 0 | 0.010 | 0.042 | 0.958 | 0.927 | 0.792 |
| WME, $c=250$, $\psi=0.03$,<br>prior=0.1 | 0.148 | 0.659 | 0.864 | 0.136 | 0.045 | 0.011 |
| WME, $c=250$, $\psi=0.03$,<br>prior=0.9 | 0.169 | 0.629 | 0.854 | 0.146 | 0.056 | 0.022 |
| WME, $c=250$, $\psi=0.03$,<br>prior=0.5 | 0.181 | 0.711 | 0.892 | 0.108 | 0.036 | 0.012 |
| WME, $c=250$, $\psi=0.015$,<br>prior=0.5 | 0.129 | 0.624 | 0.882 | 0.118 | 0.035 | 0 |
| WME, $c=25$, $\psi=0.03$,<br>prior=0.5 | 0.098 | 0.646 | 0.890 | 0.110 | 0.098 | 0 |
| WME, $c=250$, $\psi=0.03$,<br>prior=0.5, strong prior | 0.418 | 0.725 | 0.890 | 0.110 | 0.044 | 0.011 |
| WME, $c=250$, $\psi=0.03$,<br>prior=0.5, relaxed clock | 0.144 | 0.649 | 0.876 | 0.124 | 0.052 | 0.010 |

**Figure 1:**
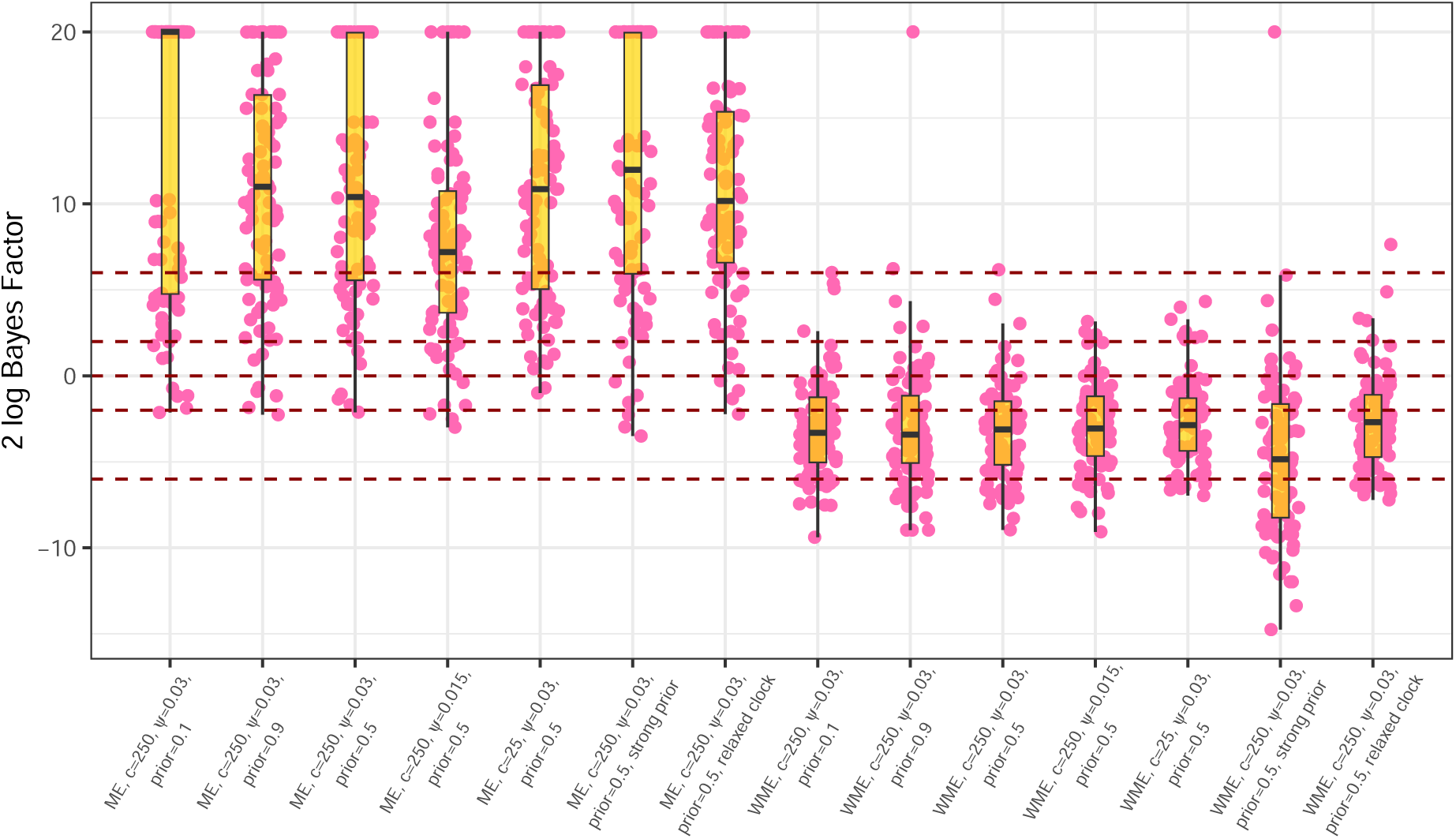
Box plot showing the 2 log BF_ME__WME_ for detecting mass extinction events using the FBD model with known mass extinction times and unknown mass extinction survival probabilities as tree priors in total-evidence dating. The yellow box plot indicates the median (center line) and the interquartile range (the box). Pink dots represent individual results from each simulated tree, while the red horizontal lines correspond to 2 log BF_ME_,_WME_ at *−*6, *−*2, 0, 2, and 6. Abbreviations are defined as follows: “ME”, simulations with mass extinction; “WME”, simulations without mass extinction; “*c*”, the number of morphological characters; “*ψ*”, the fossil sampling rate; “prior”, the prior for the existence of mass extinction; “strong prior”, the prior for the mass extinction survival probability specified as Beta(10, 40); “relaxed clock”, the UCLN relaxed clock model and HKY+Γ substitution model was used for molecular characters, and UCLN relaxed clock model and Mkv+Γ substitution model was used for morphological characters.

Reducing the fossil sampling rate decreased the BF_ME_,_WME_ in scenarios with mass extinction (Table 1, Figure 1), but it did not significantly affect the ability to identify the absence of mass extinction when none occurred (Table 1, Figure 1). Additionally, decreasing the number of morphological characters had little impact on the BF_ME_,_WME_ (Table 1, Figure 1).

Varying the prior for the existence of mass extinction did not significantly influence the BF_ME_,_WME_, regardless of whether a mass extinction event occurred (Table 1, Figure 1). However, employing a stronger prior for the survival probability of mass extinction reduced the BF_ME_,_WME_ in both scenarios with and without mass extinction (Table 1, Figure 1).

The observed patterns across all simulated trees (Table 1, Figure 1) do not necessarily apply to each individual tree (Figure S4 and S5). Within individual trees, fossil sampling rate, number of morphological characters, and priors did not exhibit a consistent pattern (Figure S4 and S5). Instead, the influence of these factors on the Bayes factor varied significantly from tree to tree (Figure S4 and S5).

In scenarios with mass extinction, the BF_ME_,_WME_ exhibited a general increasing trend with the number of species before the extinction event for both complete and rec trees (Figure S6). The trend for the reconstructed MCC tree was more complex, however, showing an initial increase to a peak, followed by a brief decrease before rising again (Figure S6B). Conversely, in the absence of mass extinction, the BF_ME_,_WME_ decreased as the number of species before the event (30Ma, despite no actual mass extinction occurring) increased for both complete and reconstructed MCC trees (Figure S7), though this decrease was less pronounced than the increase observed when mass extinction was present.

Using the FBD model that simultaneously infer the time and survival probability of mass extinctions, we found that the criterion 2 log BF_ME_,_WME_ *>* 0 successfully detected mass extinctions at the true event times (Figure S8). However, this approach frequently identified multiple mass extinction events, even when only one occurred (Figure S8), and there was considerable uncertainty in the inferred time of mass extinction (Figure S8).

#### 3.1.2 Survival Probability of Mass Extinction

In scenarios with mass extinction and a flat prior for the survival probability of mass extinction, trees with 2 log BF_ME_,_WME_ *>* 6 provided more reliable estimates of survival probabilities. In contrast, lower Bayes factor values often led to overestimation of survival probabilities (Figures 2, and S9). Estimating survival probabilities was highly uncertain (Figures 3), with significant variability across different trees (Figure S10). This pattern was consistent under both the strict and relaxed clock models (Figure 2).

**Figure 2:**
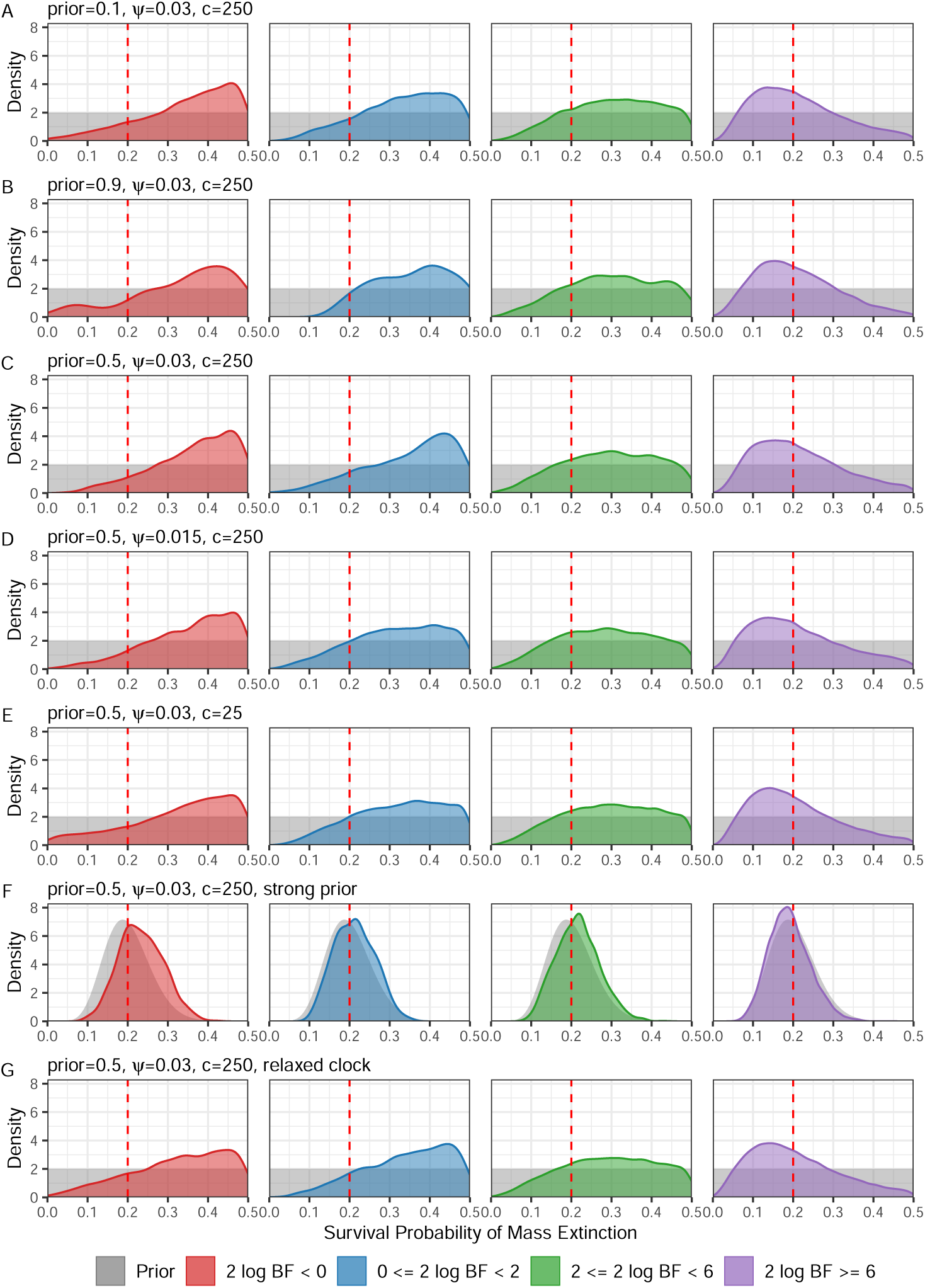
Posterior probability density plots of mass extinction survival probabilities across all simulated trees, estimated using the FBD model with known mass extinction times and unknown mass extinction survival probabilities as tree priors in total-evidence dating. To ensure equal contributions from each tree when plotting the probability density curves for specific groups (e.g., 2 log BF *<* 0 in panel A), MCMC sample sizes were normalized between replicates. Specifically, all post-burn-in MCMC chains within a group were randomly subsampled to the length of the shortest chain. The red vertical line indicates the true mass extinction survival probability (0.2). For the definitions of “ME”, “*c*”, “*ψ*”, “prior”, and “strong prior”, please refer to Figure 1.

**Figure 3:**
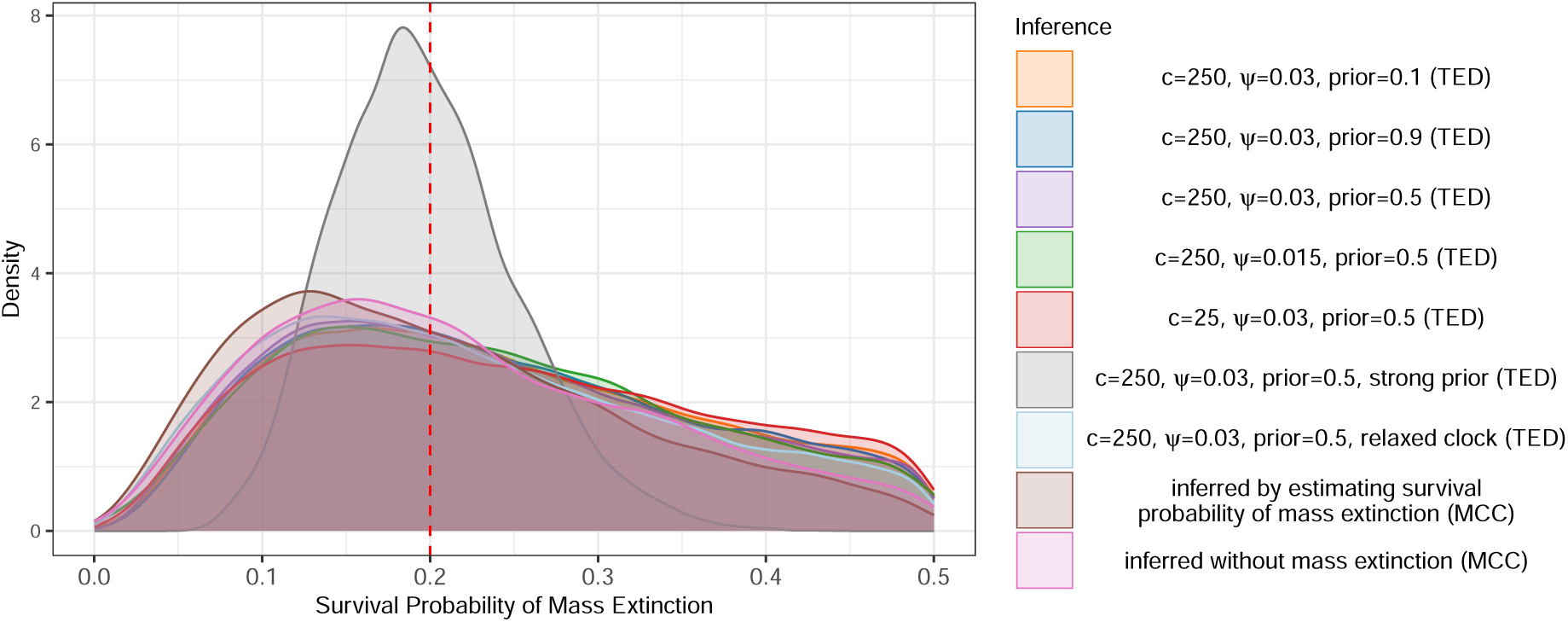
Posterior probability density plots of mass extinction survival probabilities for all simulated trees, obtained using different methods. For details on how the probability density curves were generated, please refer to Figure 2. The red vertical line indicates the true mass extinction survival probability (0.2). Abbreviations are defined as follows: “TED” refers to inferences conducted in the total-evidence dating framework; “MCC” refers to inferences based on the MCC tree. For the definitions of “*c*”, “*ψ*”, “prior”, and “strong prior”, please refer to Figure 1.

Reducing the fossil sampling rate and the number of morphological characters did not significantly impact the estimated mass extinction survival probabilities (Figures 3 and S10 to S12). However, lower fossil sampling rates caused inconsistencies in survival probability estimates for some trees compared to other methods (Figure S10).

Different prior for the existence of mass extinction did not affect the inferred survival probabilities (Figure 3, S10), also resulted in higher estimates when Bayes factors were low (Figure 2 and S9).

Using a strong prior for the survival probability of mass extinction significantly alters its posterior probability (Figure 3, S11, S10, and S12). Moreover, after applying a strong prior, the survival probability of mass extinction no longer shows significant variation with changes in the Bayes factor (Figure 2 and S9).

With a flat prior for the survival probability of mass extinction, the accuracy of the estimated survival probability improved as the number of species before the mass extinction event increased for the complete tree (Figure S13A). However, for the reconstructed MCC tree, the accuracy first decreased and then slightly increased (Figure S13B). In contrast, when a strong prior is applied, the survival probability of mass extinction remains relatively stable regardless of the number of species before the event for both complete and reconstructed MCC trees, consistently close to the true value (0.2) (Figure S13).

### 3.2 Divergence Time and Topology Inference

#### 3.2.1 Topology Inference

In scenarios with mass extinction, trees inferred using three different FBD models exhibit similar Robinson-Foulds (RF) distances to the true tree (Figure 4A). Both strict and relaxed clock models produced similar overall patterns; however, analyses under the relaxed clock consistently yielded higher RF distances (Figure 4A). A similar trend is observed when reducing the fossil sampling rate and morphological characters, with no significant decrease in RF distance compared to trees inferred with a high fossil sampling rate and more morphological characters (Figure 4A). The prior for the existence of mass extinction events and survival probability of mass extinction did not impact the inference of tree topology (Figure 4A).

**Figure 4:**
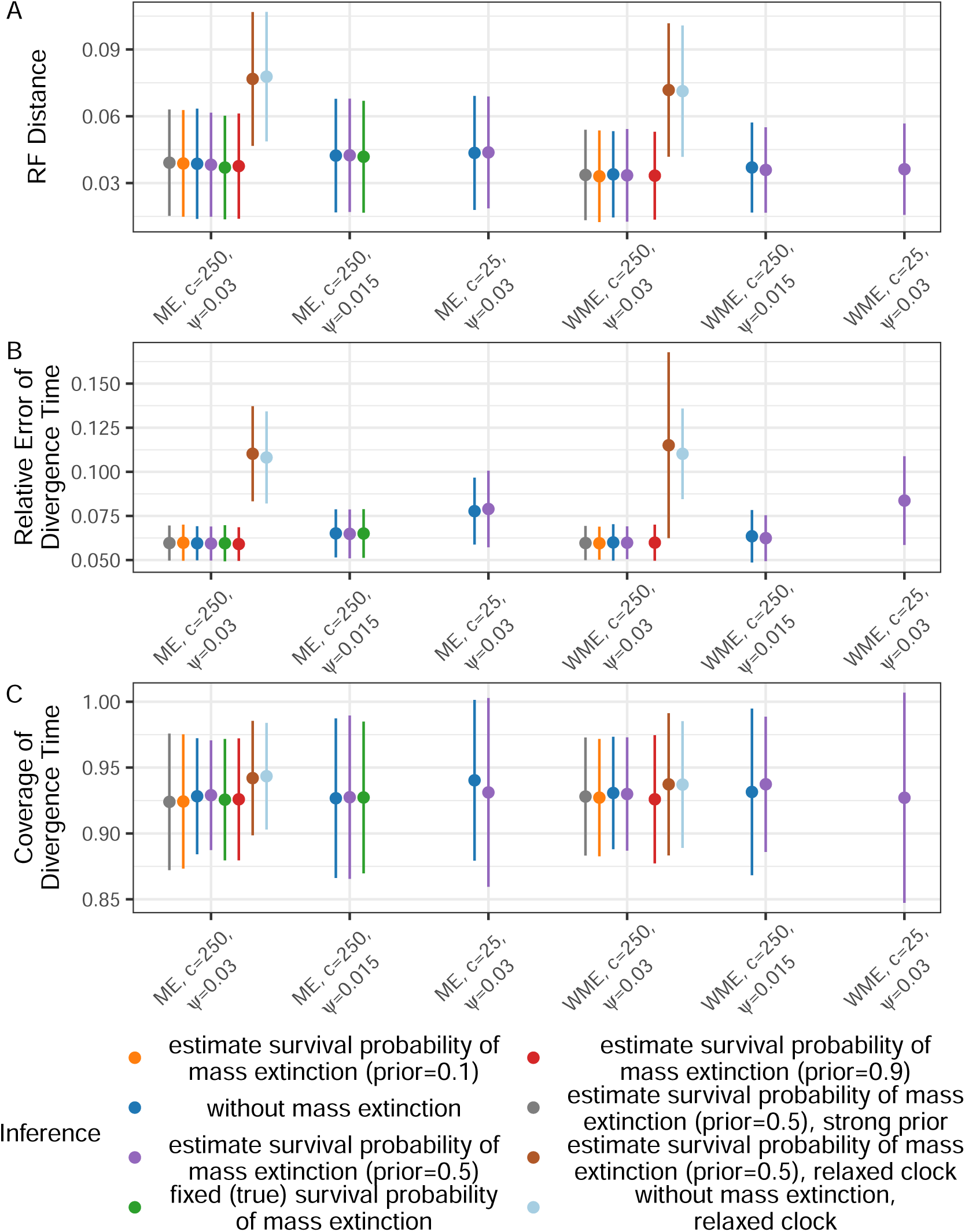
RF distance, relative error of divergence time, and coverage of divergence time between the inferred and true extant species tree when using different FBD models as tree priors. The points and bars represent the mean and standard deviation across all simulated trees. The FBD models are defined as follows: “fixed (true) survival probability of mass extinction” refers to FBD model with known (true) time and survival probability of mass extinction; “estimate survival probability of mass extinction”, refers to FBD model with known time but unknown survival probability of mass extinction; “without mass extinction” refers to FBD model without mass extinction. For the definitions of “ME”, “*c*”, “*ψ*”, “prior”, and “strong prior”, please refer to Figure 1.

In the absence of mass extinction, similar patterns were observed. Trees inferred using both the FBD model without mass extinction and the model with unknown survival probabilities showed consistent RF distances to the true tree (Figure 4A). Both strict and relaxed clocks revealed similar patterns, but the relaxed clock resulted in higher RF distances (Figure 4A), as expected since there is more uncertainty under a relaxed clock. Reducing the fossil sampling rate and the number of morphological characters yielded comparable results, with no significant decrease in RF distance compared to trees inferred with a higher fossil sampling rate and more morphological characters (Figure 4A). The prior for the existence of mass extinction events and survival probability of mass extinction did not influence the inference of tree topology (Figure 4A).

#### 3.2.2 Divergence Time Inference

In the presence of mass extinction, trees inferred using three different FBD models exhibited relative errors of divergence time (Figure 4B) and coverage of divergence time (Figure 4C) with the true extant trees that are largely consistent. Both strict and relaxed clocks revealed similar patterns, but the relaxed clock resulted in higher relative errors of divergence time (Figure 4B) A similar trend was observed when the fossil sampling rate and the number of morphological characters are reduced. However, the relative error of divergence times increased under these conditions, particularly with fewer morphological characters (Figure 4B). The prior probability assigned to the existence of mass extinction events does not impact the inference of divergence time (Figure 4B, and C).

In the absence of mass extinction, the trees inferred from both the FBD model without mass extinction and the FBD model with unknown survival probabilities also showed consistent relative errors (Figure 4B) and coverage of divergence time (Figure 4C) with the true extant tree. Both strict and relaxed clocks revealed similar patterns, but the relaxed clock resulted in higher relative errors of divergence time (Figure 4B). Similar to scenarios with mass extinction, reducing fossil sampling rates and the number of morphological characters led to decreased inference accuracy, especially with fewer morphological characters (Figures 4B and 4C). The prior probability assigned to the existence of mass extinction events does not impact the inference of divergence time (Figure 4B, and C).

The relative error of divergence time estimation for trees inferred using three different FBD models shows a consistent pattern across various divergence times regardless of the presence or absence of mass extinction events (Figure 5). Specifically, relative errors increased as divergence times approached the present, but did not show significant increases near mass extinction events (Figure 5). A similar pattern was observed when different clock models were used (Figures 5A), and fossil sampling rates (Figures 5B and E) and the number of morphological characters were reduced (Figures 5C and F).

**Figure 5:**
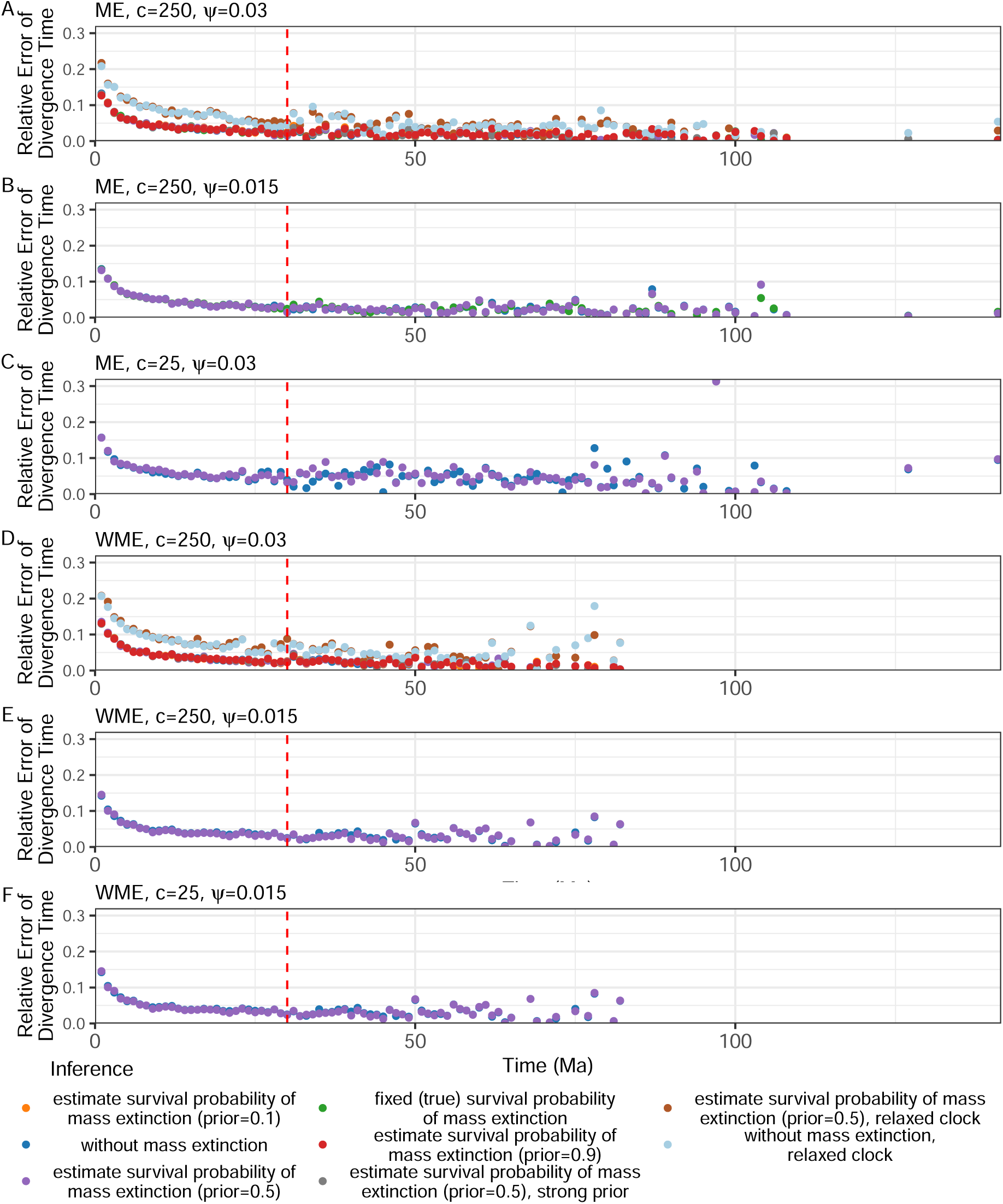
Relative error of divergence times between the inferred and true extant species tree when using different FBD models as tree priors. Each point represents a 1 million year time bin, and the error is the mean relative error of divergence times across all simulated trees within that bin. For the definitions of “fixed (true) survival probability of mass extinction”, “estimate survival probability of mass extinction”, “without mass extinction”, “ME”, “WME”, “*c*”, “*ψ*”, “prior”, and “strong prior”, please refer to Figure 1.

### 3.3 Inference of Mass Extinctions in Inferred MCC Trees

In the presence of mass extinction, the majority of MCC trees inferred using both the FBD model without extinction and the FBD model with unknown mass extinction survival probabilities successfully detected mass extinction events. When using the criterion of 2 log BF_ME_,_WME_ *>* 0, all MCC trees inferred with unknown survival probabilities and without mass extinction identified mass extinction events (Figure 6). However, this criterion often resulted in the detection of multiple mass extinction events within a single tree (Figure S14). When applying a stricter criterion of 2 log BF_ME_,_WME_ *>* 6 and excluding mass extinctions that occur too close to the root time (greater than 0.85 times the root time) and the present (less than 0.15 times the root time), to minimize the detection of false-positive mass extinction events, among the 90 converged MCC trees inferred using the FBD model with unknown survival probabilities (Table S2), 64 trees detected a single mass extinction, 7 trees detected two mass extinctions, and 19 trees did not detect any, yielding a detection rate of 71.1%. In contrast, out of the 89 converged MCC trees inferred using the FBD model without mass extinction, 64 trees detected one mass extinction event, 7 trees detected two mass extinctions, and 18 trees did not detect any, yielding a detection rate of 71.9%.

**Figure 6:**
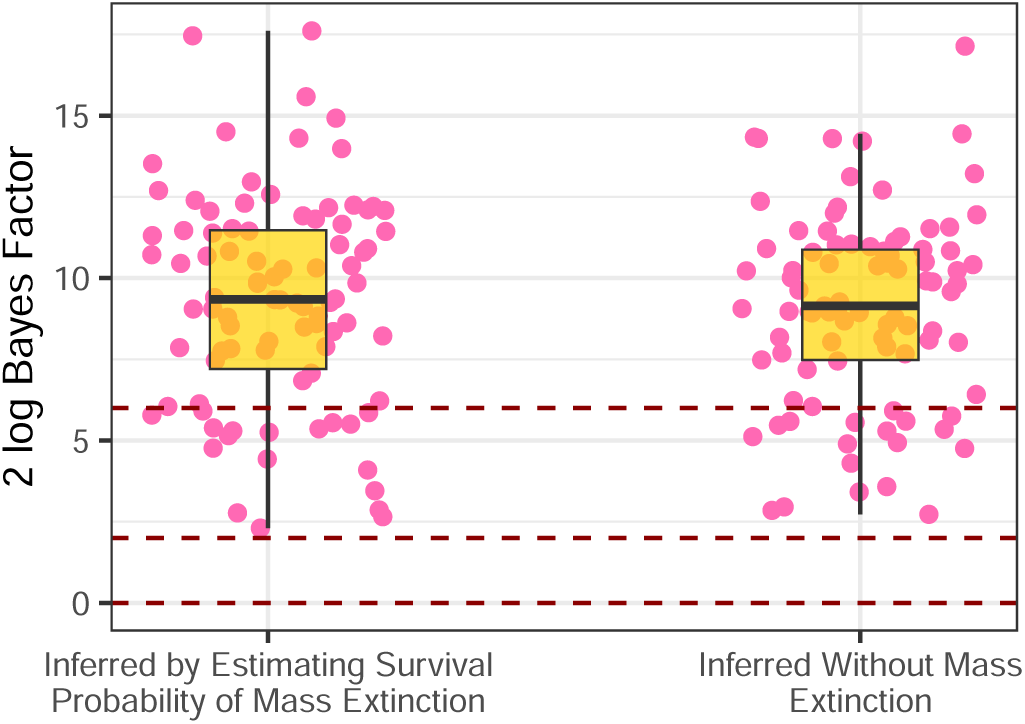
Box plot showing the maximum 2 log BF_ME_,_WME_ for all possible mass extinction periods inferred from the MCC tree. The yellow box plot indicates the median (center line) and the interquartile range (the box). Pink dots represent individual results from each MCC tree, while the red horizontal lines correspond to 2 log BF_ME_,_WME_ at 0, 2, and 6. The MCC trees were inferred using different FBD model in total-evidence dating: “inferred by estimating survival probability of mass extinction” indicates that the MCC trees were inferred using an FBD model with known mass extinction times and unknown survival extinction probabilities in total-evidence dating; “inferred without mass extinction” indicates that the MCC trees were inferred using an FBD model without mass extinction in total-evidence dating.

Using the criterion of 2 log BF_ME_,_WME_ *>* 6, the MCC trees that detect a single mass extinction provide accurate estimates of extinction times (Figure 7). The average error in estimating mass extinction time from the MCC trees inferred using the FBD model with unknown mass extinction survival probabilities is 1.02, while the average error from trees using the FBD model without mass extinction is 1.16, indicating that the estimates of extinction time were unaffected by the choice of tree model used for inference. The average length of mass extinction intervals inferred from the FBD model with unknown survival probabilities is 5.53 (Figure S15), with 52 trees (81.3%) containing the true mass extinction time within their intervals (Figure 7). In contrast, the average length of mass extinction intervals from trees using the FBD model without mass extinction is 5.56 (Figure S15), with 49 trees (76.7%) including the true mass extinction time (Figure 7).

**Figure 7:**
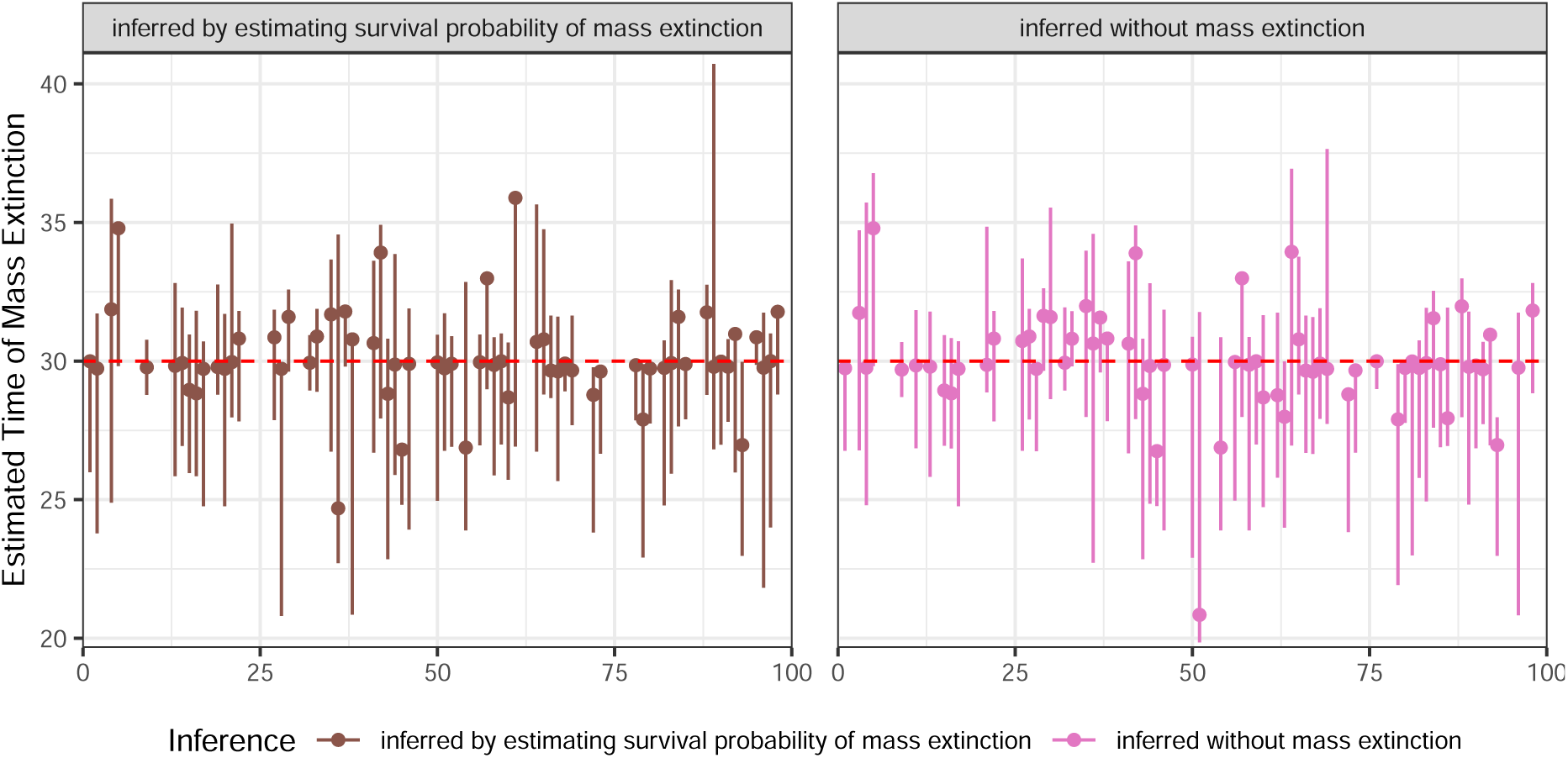
The inferred timing of a single mass extinction event detected in the MCC tree. Points represent the time with the highest Bayes factor, while the lines indicate the time intervals where 2 log BF *>* 0. The red horizontal line indicates the true time of mass extinction (30Ma). For the definitions of “inferred by estimating survival probability of mass extinction” and “inferred without mass extinction”, please refer to Figure 6.

Accurate estimates of mass extinction survival probabilities can also be obtained from MCC trees that detect a single mass extinction using the criterion of 2 log BF_ME_,_WME_ *>* 6 (Figure 3). The average estimated mass extinction survival probability from MCC trees inferred using the FBD model with unknown extinction probabilities is 0.185, while the average from trees using the FBD model without mass extinction is 0.202, indicating that mass extinction survival probability estimates were also unaffected by the choice of tree model.

Whether considering the entire set of trees (Figure S10 and S12) or individual trees (Figure 3 and S11), the posterior distribution of mass extinction survival probabilities inferred from MCC trees closely matched those obtained through total-evidence dating using the same priors, despite significant uncertainty in both approaches.

### 3.4 Diversification Rate

When using a strict clock in simulations for total-evidence dating, the prior for the existence of mass extinction events and survival probability of mass extinction, fossil sampling rates, and the number of morphological characters did not significantly influence the estimation of diversification rates (Figure S16–S33). However, when mass extinction events were present, applying an FBD model without mass extinction led to overestimation of both speciation (Figure S18–S21) and extinction rates (Figure S22–S25), with extinction rates being particularly affected. This overestimation of extinction rate consequently resulted in an underestimation of the net diversification rate (Figure S26–S29).

When using a relaxed clock, we observed a systematic underestimation of speciation (Figure S16A) and extinction rates (Figure S16B) across almost all scenarios. The sole exception to this pattern was the extinction rate, which was accurately inferred when an FBD model without mass extinction event was applied to data simulated with one (Figure S16B). This apparent accuracy may, however, be artefactual, potentially resulting from two counteracting biases: the underestimation of extinction rates inherent to the relaxed clock model and a model misspecification effect where the FBD model without mass extinction overestimates background extinction rates. The net diversification rate estimates support this interpretation. While the net diversification rate was overestimated in all other scenarios, it was underestimated when the misspecified FBD model was used on data with a mass extinction (Figure S16C). The exceptionally low coverage for the net diversification rate in this scenario further confirms its poor estimation (Figure S17C). Finally, the fossil sampling rate was consistently overestimated across all analyses, regardless of the model or the presence of a mass extinction (Figure S16D).

Applying an FBD model with mass extinction could accurately estimate diversification rates on MCC tree inferred using the FBD model without extinction and the FBD model with unknown survival probabilities (Figure S16–S33). Furthermore, the estimated diversification rates were consistent with those obtained using an FBD model with mass extinction in total-evidence dating (Figure S16–S33).

### 3.5 Empirical Data

Our application of Model 2 (see Section 2.4) to these datasets did not detect a strong mass extinction signal in either the tetraodontiform (see Section S5.2.7) or crinoid (see Section S6.2.6) empirical datasets. However, our goal was to quantify how explicitly assuming or omitting the mass extinction event within the FBD tree prior affects phylogenetic inference. Therefore, we focused primarily on Model 1 (see Section 2.4) and Model 3 (see Section 2.4), comparing the differences in the topologies and divergence times of the MCC trees inferred using these two models.

For both the tetraodontiform and crinoid empirical datasets, the choice of FBD model as tree prior—-either with a fixed mass extinction or without-—had a negligible impact on the resulting phylogenetic inferences. The MCC trees inferred under the two priors were highly congruent, showing minimal differences in both topology (RF distance) (Figures S40, S41, S49 and S50) and divergence time estimates (Figures 8, S38, S39, S47 and S48). Notably, the small discrepancies observed between the different models were comparable in magnitude to the variance found among replicate MCMC chains of the same model. This was especially apparent in the phylogeny of extant tetraodontiformes (fossils excluded) (Figures S39 and S41) and that of crinoid fossil species (sampled ancestors excluded) (Figures S48 and S50).

**Figure 8:**
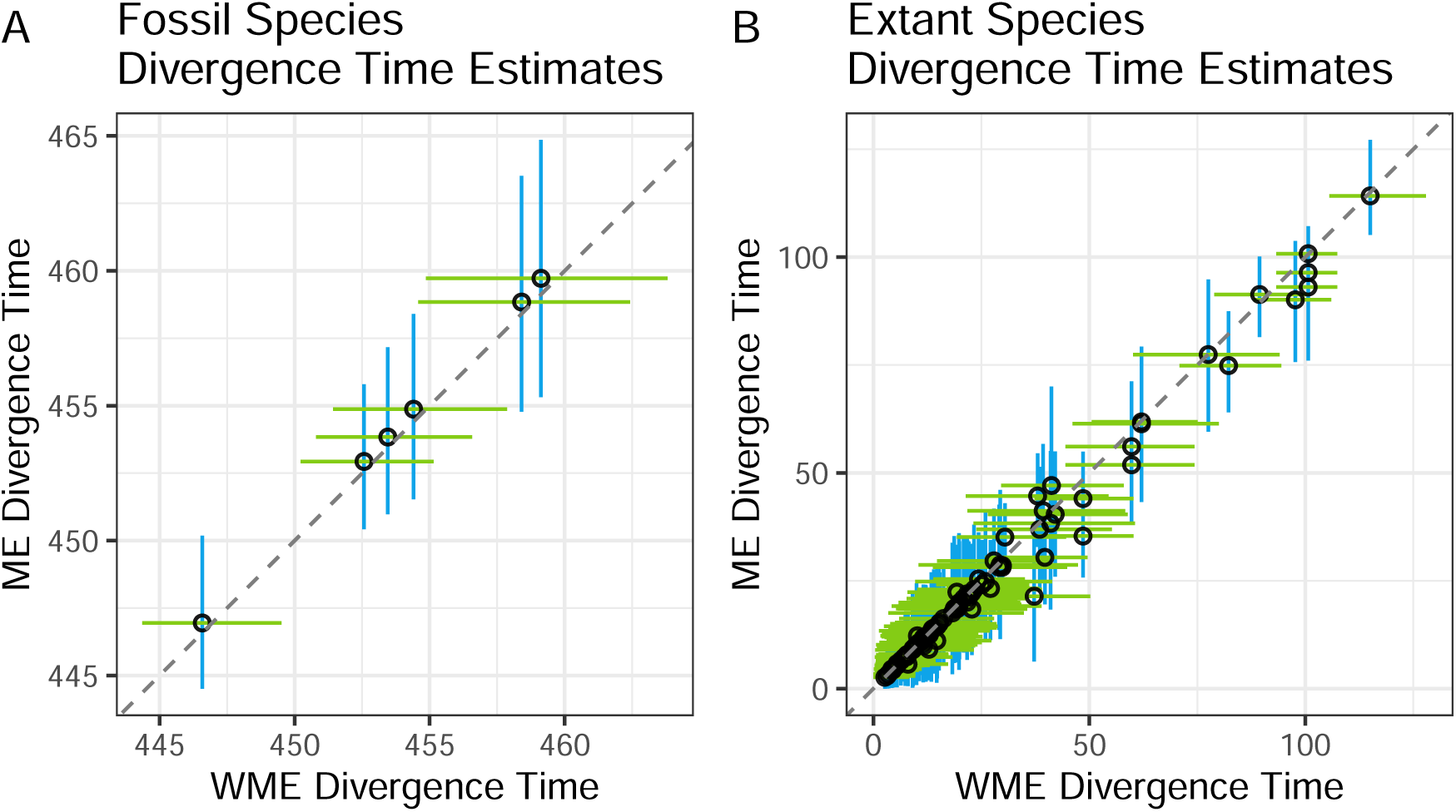
Scatter plot of (A) tetraodontiform fishes extant species and (B) crinoid fossil species divergence time estimates from RevBayes, comparing results from a FBD model without a mass extinction (WME) against a model with a mass extinction (ME). The estimates were derived from the MCC trees of the combined MCMC chains for each analysis. The green and blue lines represent the 95% HPD intervals for the divergence times. Divergence times from the analysis with mass extinction (ME) are plotted relative to the corresponding divergence time estimated from the analysis without a mass extinction event (WME).

The high degree of congruence was not confined to the overall timescale; individual divergence time estimates were also remarkably similar (Figures 8, S35 to S37 and S42 to S44). Although some divergence time estimates in the tetraodontiform dataset showed apparent differences (Figures S35-S37), these were linked to nodes with topologically inconsistent positions between the MCC trees. This consistency further extended to fossil-specific parameters, such as their inferred ages (Figure S45) and posterior probabilities of being sampled ancestors (Figure S46) in crinoids, which remained nearly identical across the two FBD model analyses. In sum, these findings strongly indicate that the incorporation of a fixed mass extinction event into the tree prior did not substantially alter the phylogenetic inference.

## 4 Discussion

### 4.1 Inference of Mass Extinctions in Total-evidence Dating

#### 4.1.1 Bayes Factor

In total-evidence dating, using the FBD model with known mass extinction times but unknown mass extinction survival probabilities proves effective for detecting mass extinctions while simultaneously performing phylogenetic inference (Table 1, Figure 1). Furthermore, when no mass extinction occurs, this model does not falsely detect one (Table 1, Figure 1). Contrasting with previous studies where mass extinction detection on a fixed tree showed lower accuracy in identifying mass extinctions compared to their absence (May et al., 2016), our results in total-evidence dating demonstrate consistently high accuracy in identifying both the presence and absence of mass extinctions (Table 1, Figure 1). This improved performance can be attributed to three key factors: (1) In our total-evidence dating approach, the mass extinction time was fixed rather than inferred, reducing the uncertainty associated with time estimation; (2) Total-evidence dating incorporates both extant species and fossil data, whereas the study by (May et al., 2016) relied solely on phylogenetic trees of extant species; (3) Total-evidence dating integrates mass extinction detection across the entire posterior distribution of trees informed by character data, whereas analyses on a single, reconstructed summary tree (e.g., an MCC tree) can lose information related to topological and temporal uncertainty. The advantages of the first and third factors are particularly stark when comparing the results of our full total-evidence dating analysis with those from a post-hoc analysis on the resulting MCC trees. Across the 90 replicate analyses that reached convergence, a mass extinction was detected in 86 replicates (accuracy: 95.6%) using a threshold of 2 log BF_ME_,_WME_ =*>* 0 (Table 1 and Figure 1). In contrast, for the corresponding 90 MCC trees, a mass extinction was detected in only 71 cases (including those with two inferred mass extinctions) using a threshold of 2 log BF_ME_,_WME_ =*>* 6 (accuracy: 78.9%) (Figure 6).

Different priors for the existence of mass extinctions yielded similar Bayes factors (Table 1, Figure 1). This result underscores the robustness of reversible jump methods in minimizing the impact of prior assumptions—whether mass extinctions are deemed likely, uncertain, or unlikely—on their detection. This robustness arises because the prior for the existence of mass extinctions influence both posterior ratio and prior ratio in Equation S1. Ultimately, these effects cancel each other out when calculating the Bayes factor, ensuring that the Bayes factor remains unaffected by the choice of prior for the existence of mass extinctions.

However, the choice of prior for the survival probability of mass extinction does affect detection outcomes. A flat prior, being more permissive, increases the likelihood of identifying mass extinction events, but it also raises the risk of misidentifying false extinctions with probabilities close to 0.5. The fundamental reason for this potential misidentification is that the data may lack sufficient signal to distinguish between a mass extinction with a survival probability near 0.5 and a scenario with no mass extinction, causing the inference to be influenced by the flat prior. In contrast, a stronger prior is more conservative, reducing the likelihood of false detections by making it harder to infer mass extinctions unless the evidence is compelling. The reason the prior for the survival probability of mass extinction influences the Bayes factor is that, although the prior for the survival probability of mass extinction does not affect prior ratio *P* (ME)*/P* (WME) in Equation (S1) of Section S2, it impacts the posterior ratio.

Reducing the fossil sampling rate decreases the accuracy of detecting both the presence and absence of mass extinctions (Table 1, Figure 1). This finding aligns with previous research suggesting that increasing fossil data can enhance the accuracy of extinction-rate estimates (Mitchell et al., 2019). Fossils offer direct information about extinct organisms, thereby improving the inference of extinction rates and the detection accuracy of mass extinction events. However, the improvement in accuracy from increasing the sampling rate appears limited in our study. This limitation could be due to two primary reasons. Firstly, the models employed—including tree models, clock models, and substitution models—are relatively simple. As a result, even with low sampling rates and a smaller number of fossils, the models can still produce accurate inferences. Secondly, the so-called “high” sampling rates used in our simulations may not be substantially high in a practical sense, leading to insufficient contrast between different sampling rates to observe a significant effect. In our simulations with the highest number of fossils—those that included a mass extinction event and high fossil sampling rate—the average number of fossils across 100 simulated trees was 29.81, with a median of 28. In comparison, we had 50 extant species in the same scenario. In empirical studies, some clades have more fossils. For example, Gavryushkina et al. (2017) documented 36 fossils and only 19 extant species in their study, and Slater et al. (2017) reported 63 fossils to 13 extant species, but the number of fossils depends on the specific clade.

Reducing the number of morphological characters does not significantly impact the detection of mass extinctions either. This outcome may also stem from the simplicity of the other models used in our analyses, which enables accurate inferences even with fewer morphological characters.

Our results show that mass extinction events can be reliably detected even when complex clock and substitution models are employed. This demonstrates that accurate inference of mass extinction is achievable, provided that an appropriate combination of tree prior, clock model, and substitution model is selected to reflect the underlying evolutionary processes.

The number of species present before the mass extinction event is a critical signal for detecting the occurrence of mass extinction in both complete and reconstructed MCC trees. In scenarios with mass extinction, the BF_ME_,_WME_ increases with the number of species present before the extinction (Figure S6). Conversely, in the absence of mass extinction, the BF_ME_,_WME_ shows an opposite trend (Figure S7). Using a threshold of 2 log BF_ME_,_WME_ = 0 to determine whether a mass extinction event has occurred, approximately 25 species in the complete tree are required before the extinction event to detect it (Figure S6A). If the fossil sampling rate decreases, this threshold rises to over 40 species (Figure S6A). Interestingly, for the reconstructed MCC tree, detection remains feasible even with very few species before mass extinction, suggesting that a strong mass extinction signal can be identified even with sparse data. In contrast, the absence of mass extinction can be confirmed with just a few species (Figure S7). Although our simulations used the same diversification rate and the same number of extant species, the number of species bofore mass extinction (30 Ma) was significantly lower in scenarios without mass extinction compared to those with mass extinction. This difference likely explains why, in the absence of mass extinction, the change in BF_ME_,_WME_ with the number of species before the extinction is less pronounced than in scenarios with mass extinction. While the number of species before a mass extinction event is a key signal, this information is entirely unknown in empirical studies. Methods such as data augmentation (Maliet and Morlon, 2022) could be employed to infer this signal, potentially improving the validation of mass extinction detection. Encouragingly, the generally positive relationship between the BF_ME_,_WME_ and species count before the mass extinction in the reconstructed MCC tree mirrors the overall trend in the complete tree (Figure S6). This suggests that the number of species before mass extinction in the reconstructed MCC tree could help assess the feasibility of detection. However, the distinct peak in the BF_ME_,_WME_ at four to five species in the reconstructed MCC tree (Figure S6B) requires further investigation to determine if it is a genuine pattern or an artifact of simulations with few species.

Using an FBD model that simultaneously infers mass extinction times and survival probabilities often leads to the identification of multiple mass extinction events (Figure S8). Although applying a stronger prior for mass extinction survival probability can help mitigate this issue, false mass extinction events may still be detected during periods with low species diversity. Addressing the influence of such periods with reduced species numbers remains a critical challenge for improving the accuracy of mass extinction detection.

#### 4.1.2 Survival Probability of Mass Extinction

With flat prior for the survival probability of mass extinction, the model performs well in estimating mass extinction survival probabilities when Bayes factors are very high (Figure 2, S9). However, for lower Bayes factors, mass extinction survival probabilities are frequently overestimated (Figure 2 and S9). Therefore, while a threshold of 2 log BF_ME_,_WME_ *>* 0 is effective for identifying the occurrence of mass extinctions, when the 2 log BF_ME_,_WME_ *<* 6, survival probabilities tend to be overestimated (Figure 2 and S9). This finding has important implications for empirical applications: the severity of a mass extinction should be interpreted with caution unless the model including the event is substantially better supported than a model without it. Theoretically, we would expect a compensatory effect where an overestimation of the mass extinction survival probability leads to a corresponding overestimation of the background extinction and net diversification rates. However, we did not observe significant compensatory changes in these background rates (Figures S52 and S53). This may be due to the high uncertainty already present in their posterior distributions and the limited impact of overestimating survival probability at a single time point on the overall background rate estimated from the origin to the present. We speculate that if a skyline model were used, this compensatory error in the background extinction rate would be more pronounced within a short time interval spanning the mass extinction event. The significant uncertainty observed in the posterior probabilities of survival probability (Figure 3, S11, S10, and S12) highlights the considerable difficulty in reliably inferring mass extinction survival probabilities.

Contrary to the detection of mass extinctions, reducing morphological character data impacts the inferred survival probability of mass extinction in certain phylogenetic trees, with some probabilities increasing and others decreasing (Figure 3, S11, S10, and S12). This influence likely arises because morphological character data are crucial for accurately quantifying the intensity of mass extinction events. However, overall, neither fossil sampling rates nor morphological character data substantially enhance the accuracy or reduce the uncertainty in estimating mass extinction survival probabilities (Figure 3, S11).

As expected, the priors for the existence of a mass extinction event occurring do not influence the estimation of the magnitude (survival probability) of these events (Figure 3, S11, S10, and S12).

The choice of priors for the survival probability of mass extinction plays a crucial role in its inference (Figure 3, S11, S10, and S12). Therefore, using a stronger prior—particularly one informed by robust paleontological or paleoenvironmental evidence—can not only enhance the accuracy of detecting the presence or absence of mass extinction events (Table 1, Figure 1) but also significantly improve the precision of estimating the survival probability of mass extinction when such events occur (Figure 3, S11, S10, and S12). Given that fossil sampling rates and the number of morphological characters have minimal influence on the inference of the survival probability of mass extinction, selecting an appropriate prior for the probability of mass extinction becomes even more critical in ensuring accurate results.

Similar to the Bayes factor, the accuracy of inferring the survival probability of mass extinction increases with the number of species present before the extinction event in the complete tree (Figure S13A). However, achieving a relatively accurate estimate of the survival probability of mass extinction requires approximately 100 species before mass extinction (Figure S13A). This finding highlights the necessity of a strong mass extinction signal to reliably infer the survival probability of mass extinction. Likewise, the number of species before mass extinction in the reconstructed MCC tree provides information (Figure S13B), but the exceptionally high accuracy observed at four and five species requires further investigation to determine whether it is a genuine pattern or a statistical artifact.

### 4.2 Accuracy of Different FBD Models as Tree Priors

Using different FBD models as tree priors does not significantly affect the estimation of phylogenetic tree topology and divergence times. This consistency may be attributed to the simplicity of the models employed in our simulations. According to equation (8) in May and Rothfels (2023), the posterior probability consists of four components: (1) Parameter priors: This includes priors on the FBD, morphological and molecular substitution, and clock model parameters. (2) Likelihood of fossil ages: This is based on the FBD model parameters. (3) Phylogenetic tree prior: This is based on the FBD model parameters and fossil ages. (4) Likelihood of morphological and molecular characters: This is based on the phylogenetic tree, the substitution and clock model parameters. As a result, if an incorrect FBD model is used (e.g., an FBD model without a mass extinction for a scenario where a mass extinction did occur), the true phylogenetic tree’s prior probability (component 3) will be low. However, because the true phylogenetic tree accurately reflects evolutionary relationships and divergence times, the likelihood of morphological and molecular characters (component 4) will remain high. Conversely, a phylogenetic tree constructed using an incorrect FBD model may have a high tree prior (component 3), but its evolutionary relationships and divergence times will be inaccurate, resulting in a low likelihood for morphological and molecular characters (component 4). The posterior probability is ultimately a balance between components (3) and (4). If the influence of the tree prior (3) outweighs that of the likelihood (4), an incorrect phylogenetic tree may be inferred. On the other hand, if the likelihood of morphological and molecular characters (4) dominates, it is still possible to infer the correct phylogenetic tree, even with an incorrect FBD model.

This robustness suggests that total-evidence dating is resilient to the choice of tree prior, aligning with findings from previous studies (Barido-Sottani and Morlon, 2025). Considering that prior total-evidence dating studies often did not incorporate FBD models with mass extinctions, the similarity in inferred tree topologies and divergence times across different FBD models is encouraging. It indicates that even if mass extinctions have occurred, they may not substantially impact inference results. However, further validation using real datasets is necessary to confirm this observation.

The robustness of the inferred topology to the choice of FBD tree prior, even under complex model conditions, highlights the strong phylogenetic signal within the character data. However, the complexity of the clock and substitution models themselves proved to be a critical factor. Even when the inference model was correctly specified to match the simulation model, our analyses under complex conditions resulted in lower topological accuracy than analyses under simple conditions. This suggests that the increased parameter space of more realistic models can introduce estimation error that degrades inference accuracy, underscoring the need for larger character datasets to reliably support such complex models.

The fossil sampling rate had minimal impact on the accuracy of inference results (Figure 4), which may again be due to the simplicity of the other models employed, leading to accurate inferences regardless of sampling intensity.

Consistent with previous findings (Barido-Sottani et al., 2020), our study demonstrates that having very few morphological characters (25 in this case) significantly increases the relative error of divergence time (Figure 4B). This reduction in accuracy likely arises because morphological characters play a critical role in total-evidence dating. These characters are present in both fossils and extant species, providing essential information that links fossils to living taxa. All data about already extinct species are preserved in fossils, and morphological characteristics serve as key indicators for placing these species within the phylogenetic framework. Additionally, morphological clocks offer valuable insights into evolutionary timelines and phylogenetic relationships. However, the coverage of divergence time (Figure 4C) and the accuracy of topology (Figure 4A) estimates did not decrease, indicating that compared to the relative error of divergence time, these two metrics are easier to infer accurately.

Although different FBD models can produce similar phylogenetic trees, our results reveal a critical consequence of model misspecification on the inference of diversification rate. Specifically, when a mass extinction event has occurred but is omitted from the inference model, the background extinction rate is significantly overestimated, leading to a corresponding underestimation of the net diversification rate. This finding highlights the importance of carefully selecting the appropriate FBD model when analyzing diversification rates. The mechanism underlying this bias is a classic example of parameter compensation. The inference framework attempts to explain the observed data—including the sudden loss of many lineages at the time of the mass extinction—using only the parameters available in the specified model. Lacking a parameter for a pulsed extinction event, the model is forced to attribute the signal of catastrophic lineage loss to the only available mechanism for extinction: the continuous background extinction rate. To account for the sharp decline in lineages, the model inflates the estimate of extinction rate across the entire tree, making the observed pattern more likely under the misspecified model. This result serves as a crucial cautionary note for empirical studies: an inferred period of exceptionally high background extinction could, in some cases, be an analytical artifact caused by an unmodeled, pulsed extinction event.

### 4.3 Inference of Mass Extinctions in Inferred MCC Trees

Unlike previous studies that analyzed mass extinctions using trees composed solely of extant species (May et al., 2016), our MCC trees include fossils. Fossils provide direct information about extinct organisms, and prior research has demonstrated that incorporating fossils enhances the accuracy of extinction rate estimations (Mitchell et al., 2019). Similarly, including fossils may improve the precision in detecting mass extinction events. By integrating fossil data, we avoided the need to impose strong priors on survival probabilities (often assumed to be very low), which in earlier studies may lead to unreliable survival probability estimates (May et al., 2016). Instead, we employed flatter prior distributions for both extinction timing and survival probability. Our results indicate that we can simultaneously estimate mass extinction times (Figure 7) and survival probabilities (Figure 3); however, the posterior distributions for survival probabilities still exhibit considerable uncertainty.

Detecting mass extinctions on MCC trees requires a higher Bayes Factor threshold for significance compared to the total-evidence dating analysis, where a threshold of 2 log BF_ME_,_WME_ *>* 0 (i.e., any positive support) is sufficient. For the MCC tree analyses, and consistent with previous studies on extant species trees (May et al., 2016), we recommend using a threshold of 2 log BF *>* 6 as the criterion for significant mass extinction events. However, even with this stricter threshold, our analysis of MCC trees occasionally inferred two mass extinctions, suggesting a tendency to identify false positives. This issue might be mitigated by applying stronger priors on the survival probability.

Our analysis of MCC trees also highlights the potential for inferring spurious mass extinctions near the root and the present. Previous studies have shown that extinctions occurring during these periods are challenging to detect accurately (May et al., 2016). Therefore, any mass extinctions detected at these times may not be reliable. In our analysis, we excluded mass extinctions inferred near these critical periods. Future research should consider incorporating constraints directly into the prior distributions, setting the survival probabilities to one during these intervals, rather than manually removing them post hoc. When lineage counts are low, such as near the root of a phylogeny, background extinctions can be statistically mistaken for mass extinctions. This is because the loss of even a few species can represent a high proportion of the total diversity (e.g., 3 out of 5). This small-sample-size artifact makes low survival probability an insufficient sole criterion for identifying true mass extinctions, rendering inferences from these periods unreliable. To reduce the false detection of such events, the survival probability criterion should be supplemented with a minimum threshold for the absolute number of extinct species (e.g., at least 10). Developing methodologies to accurately infer extinctions during these challenging periods or demonstrating their inherent undetectability remains an area for further investigation. This is particularly important as the detection of mass extinctions near the present is highly relevant due to the ongoing debate about a potential sixth mass extinction (Barnosky et al., 2011), and some recent studies using the FBD model have indeed observed a sharp increase in extinction rates towards the present (Allen et al., 2025).

### 4.4 Consistency Between Total-Evidence Dating and Inferences from MCC Trees

Our study demonstrates that survival probabilities inferred through total-evidence dating are largely consistent with those derived from the MCC tree with same prior (Figure 3, S11, S10, and S12). Notably, even when using an FBD model that does not include mass extinctions in the total-evidence dating analysis, the mass extinction survival probabilities obtained from the inferred MCC tree remain consistent with those derived using an FBD model that incorporates mass extinctions.

Similarly, diversification rates estimated through total-evidence dating align with those derived from the MCC tree when using the same FBD model (Figure S16–S33). Although employing an FBD model without mass extinctions in total-evidence dating may lead to an underestimation of extinction (Figure S22–S25) and net rates (Figure S26–S29), applying an FBD model that includes mass extinctions on the MCC tree can still produce extinction and net rate estimates consistent with those from total-evidence dating that includes mass extinctions. For large datasets, it may be feasible to initially use a simpler FBD model for total-evidence dating and subsequently apply a more complex FBD model to the MCC tree for diversity rate analysis. Additionally, conducting model selection with different tree models on the MCC tree could help identify and address any model mismatches present in the total-evidence dating analysis. However, it is important to note that all these approaches rely on the accuracy of the substitution and clock models, which are crucial for ensuring the reliability of the phylogenetic tree.

### 4.5 Empirical Data

Although our analyses under the Model 2 (see Section 2.4) did not recover a significant signal for a mass extinction event in either dataset, this finding should not be interpreted as conclusive evidence that no such extinction occurred. Our use of a constant-rate model, which assumes uniform speciation and extinction rates through time, represents a significant simplification of complex evolutionary histories. Furthermore, our random sampling of extant species may also contribute to the non-detection of an extinction signature. This contrasts with the approach of Arcila and Tyler (2017), who successfully identified a mass extinction in Tetraodontiformes. Their success was likely attributable to a combination of methods better suited for this question: they employed a skyline model, which explicitly allows for diversification rates to vary through time, and utilized a diversified sampling strategy designed to maximize the phylogenetic diversity of extant taxa included in the analysis.

Therefore, the absence of a detectable signal in our study may be a consequence of these methodological constraints rather than a reflection of the true evolutionary history of the group. We conclude that while our analysis does not support a mass extinction hypothesis under a constant-rate framework, this question remains open. Future investigations employing alternative, variable-rate models (e.g., skyline approaches) and more strategic sampling protocols will be essential to conclusively test for the signature of a mass extinction.

Nevertheless, there was no significant difference between the phylogenies inferred under the FBD model that enforced a mass extinction (see Section 2.4) and the model that did not (see Section 2.4). This finding is congruent with the simulation results, providing further evidence that the selection of an FBD model with or without a mass extinction exerts minimal influence on phylogenetic inference within a total-evidence dating framework.

### 4.6 Future Research Questions

Following on the findings of our study, several key issues warrant further investigation.

Firstly, our current inferences are predicated on the assumption that the timing of mass extinction events is known and accurate. Integrating an FBD model that simultaneously infers both extinction timing and survival probability within the total-evidence dating framework remains unresolved. While theoretically feasible, employing such a model poses significant computational challenges and may lead to overly flat posterior distributions for mass extinction. This could result in the inference of extinctions across multiple time periods, diluting the signal of any single extinction event. Deep learning techniques might offer a promising solution to this problem. For instance, deep learning could be utilized initially to estimate mass extinction time from molecular and morphological characters, which could then inform an FBD model with known extinction times but unknown survival probabilities in subsequent total-evidence dating analyses.

Secondly, the assumption of known and precise extinction times may not be fulfilled by all empirical datasets, and the implications of incorrect extinction time settings are not yet fully understood. For example, if we set the extinction time to 30Ma while the actual extinction occurred at 40Ma, several questions arise: Would we still detect a mass extinction at 30Ma? Would the inferred MCC tree be able to detect the mass extinction at all? If a mass extinction is detected on the MCC tree, would it be reported at 30Ma or at the true time of 40Ma? Exploring these scenarios is essential to understand the robustness of total-evidence dating under conditions of temporal uncertainty.

Thirdly, our conclusions are predicated on highly idealized conditions, including known models of trees, clocks, and substitution, along with accurate priors and precise fossil ages. While consistent results were obtained under both simple and complex simulated scenarios, the application to empirical data presents greater complexity. In empirical studies, the appropriate models of trees, clocks, and substitution, as well as their corresponding priors, are often unknown. Moreover, factors such as incomplete character matrices and the inherent uncertainty of fossil ages can further complicate the inference of mass extinctions. The absence of a detected mass extinction in our empirical analysis underscores these challenges. This result does not necessarily negate the existence of a mass extinction but may instead reflect the selection of inappropriate models or an insufficient extant species sampling strategy. Additional investigation is required to fully understand the influence of these uncertainties on the inference of mass extinction events.

Fourthly, our study employed a very low survival probability (0.2) to facilitate the detection of the mass extinction event. It is recognized that some mass extinctions in the geological record may have been characterized by even lower or, conversely, higher survival probabilities (Bambach, 2006; Fan et al., 2020). However, the detectability of such an event is not solely a function of its severity. Rather, statistical power is determined by a complex interplay of factors—including the size of the phylogeny, the timing of the extinction event (May et al., 2016), and the background extinction rate—the individual and synergistic effects of which remain incompletely understood. For instance, a mass extinction with a comparatively high survival probability could nonetheless produce a detectable signal in a large phylogeny. Conversely, even a catastrophic extinction might be virtually undetectable if inferred from a small tree. Likewise, the detection of mass extinctions is impeded by high background extinction rates but facilitated by low ones. Our study utilized a single background extinction rate of 0.15; therefore, the interplay between a broader range of background extinction rates, survival probabilities, and statistical power for detecting mass extinctions merits further investigation.

Fifthly, distinguishing between different patterns of mass extinction is challenging, which involves a choice between two distinct modeling philosophies. Our study models mass extinctions as instantaneous “pulse” events, where each lineage faces a probability of survival. This approach is conceptually suited for testing hypotheses about rapid, catastrophic triggers like a bolide impact. Its primary advantage is that it provides a direct statistical test for an abrupt, geologically instantaneous event, isolating it from background rate dynamics. An alternative and valuable approach is the skyline FBD model, which can model a protracted “press” of elevated extinction risk lasting millions of years, as might be caused by long-term climate change. For instance, one could use this framework to specify a three-interval model: a background extinction rate, a mass extinction interval with a much higher extinction rate (comprising both background and mass extinction components, for which a prior favoring higher values could be specified), followed by a return to the background level—an approach whose effectiveness for detecting rate shifts has been demonstrated in previous studies (Culshaw et al., 2019). This framework is more appropriate for testing hypotheses about extended extinction crises. The choice between models, therefore, depends on the underlying hypothesis. The instantaneous model directly tests for a “pulse”, while the skyline model tests for a “press”. However, this distinction presents a significant analytical problem. An instantaneous-event model may incorrectly infer a single pulse from a prolonged period of high extinction, while a skyline model could “smear” the signal of a truly abrupt event across a time interval, underestimating its severity. This challenge is compounded when background extinction rates themselves fluctuate over time, as such variations can mask or be mistaken for a true mass extinction event. The conditions under which these distinct extinction patterns can be statistically distinguished remain unresolved. A further consideration is the potential for sampling heterogeneity driven by research effort. Mass extinction intervals are subjects of intense paleontological focus, leading to an extreme spike in fossil collection that may be difficult for standard skyline model to fully capture. The estimated sampling rate may therefore still represent an underestimation of the true sampling rate during that specific period. Given the close relationship between parameters in the FBD model, to reconcile the observed high density of fossils with this underestimated sampling probability, the model would be forced to infer a higher extinction rate for that mass extinction interval. Paradoxically, while this would bias the parameter estimates, such a bias would likely amplify the signal of the mass extinction, making it more, rather than less, detectable.

Furthermore, the distinction between pulse and press models is itself a simplification, as many historical crises were more complex. The Late Ordovician mass extinction, for example, is now understood to be a multi-pulse event comprising at least two distinct extinction episodes separated by approximately one million years (Harper et al., 2014; Rasmussen et al., 2023). The current instantaneous-event model would likely conflate such closely spaced events into a single, larger extinction. On the other hand, it may incorrectly identify a single event as a multi-pulse one. For instance, in our simulations with a single known mass extinction, the inference on the MCC tree sometimes split this event into two adjacent intervals with high survival probabilities; in such cases, we consolidated the signal by multiplying these survival probabilities. However, with empirical data, it is impossible to distinguish between such a model artifact and a true multi-pulse extinction event. Therefore, a key goal for future methodological development is to create a framework capable of discerning between a single catastrophic event, a series of rapid pulses, and an extended period of elevated extinction pressure.

Finally, an important consideration is identifiability. Previous studies have established that time-dependent FBD models without mass extinctions are identifiable, if the removal-after-sampling probability is fixed to a value less than 1 (Truman et al., 2025). This probability refers to the probability that a lineage is removed from the diversification process after it has been sampled (for a full explanation, see Truman et al. 2025). In the macroevolutionary context of our study, this probability is fixed to zero. However, it remains unclear whether identifiability is preserved in a time-dependent FBD framework where mass extinction events are included and the removal-after-sampling probability is fixed to zero.

Overall, addressing these questions will enhance our understanding of the impact of mass extinctions on phylogenetic inference and improve the applicability of total-evidence dating in more complex and realistic scenarios.

## 5 Conclusion

By simulating phylogenetic trees with and without mass extinction events and employing different FBD models as tree priors in total-evidence dating, we have arrived at the following three key findings:

1. When the timing of a mass extinction is known, using an FBD model that includes a mass extinction with a fixed (and true) extinction time but an unknown survival probability as the tree prior allows us to detect the occurrence of the mass extinction and estimate the survival probability during phylogenetic inference. Importantly, this approach does not lead to false positives when no mass extinction has occurred.
2. In scenarios where a mass extinction has occurred, using different FBD models—as tree priors in total-evidence dating—yields phylogenetic trees with similar topologies and divergence times. This holds true whether the FBD model lacks mass extinction, includes mass extinction with fixed (and true) extinction time and fixed (and true) survival probability, or includes mass extinction with fixed (and true) extinction time but unknown survival probability. However, using a model with no mass extinction leads to biased estimates of the extinction and net diversification rates.
3. Even when using an FBD model without mass extinction or one with mass extinction and fixed (and true) extinction time but unknown survival probability as tree priors in total-evidence dating, we can still detect the occurrence of mass extinction events in the inferred phylogenetic trees when a mass extinction has indeed occurred.

It is important to note that these conclusions are contingent upon the correct selection of FBD models, clock models, substitution models, and prior distributions in the total-evidence dating analysis.

## Acknowledgements

We would like to express our sincere gratitude to the Associate Editor April Wright, Bethany Allen, and two anonymous reviewers for their insightful comments and suggestions on a previous version of this manuscript. We thank the Morlon team at IBENS for their very helpful suggestions and feedback on an earlier draft of this manuscript. JBS received support from the European Union’s Horizon 2020 Research and Innovation Programme under the Marie Sk-lodowska-Curie grant agreement No. 101022928. MD received support from the China Scholarship Council (grant number 202306370127) and the Society of Systematic Biologists Early Systematists Award. JT received support from the National Natural Science Foundation of China (grant number U23B20155) and the Science and Technology Innovation Program of Hunan Province (grant number 2023RC1021). WW received support from the National Natural Science Foundation of China (grant numbers 42472027, 42342042) and the Natural Science Foundation of Hunan Province (grant number 2023JJ20063). We used OpenAI’s ChatGPT to refine the language of the manuscript.

## Data Availability Statement

The R scripts used for simulation, analysis and plotting, and the RevBayes, BEAST2, MrBayes scripts used for inference can be found in the Dryad repository.

