## Supplementary information for "Evaluating the impact and detectability of mass extinctions on total-evidence dating"

#### S1 Expected Substitutions Per Site

For each simulated tree, we calculated the expected number of substitutions per site for the molecular characters under scenarios with and without mass extinction, using both strict and UCNL relaxed clocks (Figure S34). Furthermore, across the 100 simulated trees, we calculated the mean and median for both the expected number of substitutions per site and the number of invariant characters.

#### S2 Bayes Factor

To test the hypothesis that a mass extinction occurred at a specific time $t$ (in this case, 30 Ma), we define the Bayes factor comparing the presence ("ME") versus the absence ("WME") of a mass extinction at $t$:

$$\mathrm{BF}_{\mathrm{ME,WME}} = \frac{P(\mathrm{ME} \mid X)}{P(\mathrm{WME} \mid X)} \div \frac{P(\mathrm{ME})}{P(\mathrm{WME})} = \frac{P(\mathrm{ME} \mid X)}{1 - P(\mathrm{ME} \mid X)} \div \frac{P(\mathrm{ME})}{1 - P(\mathrm{ME})} \tag{S1}$$

where "ME" represents the scenario where a mass extinction occurs at time $t$, and "WME" represents the scenario without a mass extinction at that time. Since "ME" and "WME" are mutually exclusive and exhaustive events, their probabilities satisfy $P(\mathrm{ME}) + P(\mathrm{WME}) = 1$.

When we employed $1 -$ `dnReversibleJumpMixture(0.0, dnUniform(0.5, 1.0), 0.5)` as the prior distribution for the survival probability at time $t$, this prior reflects the assumption that a mass extinction at time $t$ is equally as likely as no extinction, assigning:

$$P(\mathrm{ME}) = \frac{1}{2} \quad \text{and} \quad P(\mathrm{WME}) = 1 - P(\mathrm{ME}) = \frac{1}{2} \tag{S2}$$

Given these equal prior probabilities, the Bayes factor simplifies to the ratio of posterior probabilities:

$$\mathrm{BF}_{\mathrm{ME,WME}} = \frac{P(\mathrm{ME} \mid X)}{1 - P(\mathrm{ME} \mid X)} \tag{S3}$$

The posterior probability $P(\mathrm{ME} \mid X)$ can be approximated directly from the MCMC samples by calculating the proportion of samples where the extinction probability at time $t$ is greater than or equal to 0.5:

$$P(\mathrm{ME} \mid X) = \frac{1}{N} \sum_{i=1}^{N} \begin{cases} 1 & \text{if probability of Mass Extinction at } t \geq 0.5 \\ 0 & \text{if probability of Mass Extinction at } t = 0 \end{cases} \tag{S4}$$

where $N$ is the number of MCMC samples. This approach allows us to quantify the evidence supporting a mass extinction event at time $t$ based on our data $X$.

### S3 Justification for Simulation Filtering Criteria

To justify and quantify the impact of our tree and fossil filtering criteria, we performed a larger simulation for each of the four primary scenarios (with/without mass extinction, and high/low fossil sampling rate), generating 1,000 birth-death trees for each. We first filtered these trees based on their origin times, excluding those falling outside our predefined ranges as specified in the main text (e.g., <34 Ma or >125 Ma for the scenario without mass extinction). For the remaining set of trees in each scenario, we simulated the fossil sampling process and applied a second filter, excluding simulations that resulted in fossil counts outside our specified minimum and maximum bounds.

This two-step filtering process ensured that our final dataset did not contain extreme outliers that could disproportionately influence the results. For example, in the "without mass extinction, $\psi$=0.015" scenario, 22 trees were excluded for having an origin time less than 34 Ma and 31 trees were excluded for having an origin time greater than 125 Ma. From the remaining 947 trees, subsequent fossil-based filtering removed 25 trees for having fewer than 6 fossils and 28 trees for having more than 28 fossils (Table S3).

This analysis demonstrates that our filtering criteria were not overly restrictive, as only a small fraction of the total simulations were excluded. This ensures the removal of extreme outliers without introducing significant bias or fundamentally altering the core dataset. It's worth noting that in our simulations in the manuscript, we did not pre-specify a fixed number of trees and then discard those that did not meet the criteria. Instead, we continued simulating until we obtained 100 FBD trees that satisfied the conditions.

### S4 FBD model with unknown time and survival probability of mass extinction

We employed the FBD model to simultaneously infer the timing and survival probability of mass extinction events. Although the exact timing of extinctions is unknown, it is not entirely arbitrary (Figure S8). The model allows mass extinctions to occur at specific time points: $1, 2, \ldots, 60$, resulting in 59 permissible time points. To ensure that the total prior probability for the existence of mass extinction across all these time points sums to 0.5, the prior probability for the existence of a mass extinction at each individual time point is set to 0.5/59. At each permissible time point, the prior for the survival probability follows Uniform$(0.0, 0.5)$, maintaining consistency with the model's assumptions. At the actual time of the mass extinction event (time point 30), the only difference from the previous model with a fixed extinction time is that the prior probability for the existence of a mass extinction has been adjusted from 0.5 to 0.5/59.

### S5 Tetraodontiform Fishes Analysis

We re-analyzed the total-evidence dataset for tetraodontiform fishes from Arcila and Tyler (2017). The original study identified a mass extinction event at the Paleocene–Eocene Thermal Maximum ($\sim$55 Ma). This inference was not derived directly from their total-evidence dating analysis, which used a constant-rate FBD model as the tree prior. Instead, the mass extinction signal was detected in a subsequent, separate analysis that applied a skyline birth-death model to the majority-rule consensus tree of extant species.

#### S5.1 Data

We used the dataset from Arcila and Tyler (2017) with a modification to the outgroup sampling. From the original matrix, we removed the two fossil outgroups, *Lophiodes monodi* and *Cyttus novaezealandiae*, because they only possess morphological characters and lack molecular characters, retaining *Antigonia capros* as the sole outgroup taxon for all analyses. We adopted the same data partitioning scheme as (Arcila and Tyler, 2017). The dataset comprised a total of 95 extant species (including the outgroup) and 52 fossil taxa, each with a corresponding temporal age range. The character matrix included 14,648 molecular and 210 morphological characters.

#### S5.2 Models

Phylogenetic inference was performed using the total-evidence dating method implemented in `RevBayes` (Höhna et al., 2016). It should be noted that we did not have access to the MrBayes configuration files from Arcila and Tyler (2017), only their result files; therefore, we opted for weakly informative priors wherever possible. Furthermore, in the supplementary material of Arcila and Tyler (2017), all posterior trees from all eight chains, including the burn-in phase, had identical topologies.

##### S5.2.1 Molecular Substitution Model

We modeled nucleotide substitution for the molecular data partition using the General Time Reversible (GTR) model. This model was combined with a parameter for a proportion of invariable sites (+I) and a gamma distribution to account for rate heterogeneity across variable sites (+Γ). The continuous gamma distribution was approximated using five discrete rate categories.

##### S5.2.2 Morphological Substitution Model

We modeled the evolution of the morphological characters using the Mk model, where the number of states (k) was set to the maximum number of states observed across all characters in the dataset. To correct for the ascertainment bias that can arise from analyzing datasets that exclude constant characters, we applied the Mkv model variant.

To account for heterogeneity in evolutionary rates among the morphological characters, we applied a gamma distribution ($+\Gamma$). A single gamma distribution, common to all morphological characters and their state transitions, was used to model this rate variation and was approximated using ten discrete rate categories.

##### S5.2.3 Molecular Clock Model

We modeled the evolution of the molecular data using the UCLN relaxed clock model. The prior for the mean evolutionary rate was an exponential distribution with a rate of 500. The prior for the standard deviation of the lognormal distribution was an exponential distribution with a rate of 5.

##### S5.2.4 Morphological Clock Model

We modeled the evolution of the morphological data using the UCLN relaxed clock model. The prior for the mean evolutionary rate was an exponential distribution with a rate of 100. The prior for the standard deviation of the lognormal distribution was an exponential distribution with a rate of 5.

##### S5.2.5 Tree Model

We employed three constant-rate FBD models as tree priors, each with exponential priors: speciation and extinction rates had an exponential prior with a rate of 10, and the fossilization rate had an exponential prior with a rate of 30. Fossil ages followed a uniform distribution. The three FBD models were:

1. FBD model without mass extinction.

2. FBD model with a fixed mass extinction time of 55 Ma, while treating the survival probability as an unknown parameter. Following the third model in Section 2.2.1, the prior for the survival probability was defined using a reversible-jump mixture model framework. An equal prior probability of 0.5 was assigned to the models with and without a mass extinction event. In the model that includes a mass extinction, the survival probability is treated as an unknown parameter drawn from a $\text{Uniform}(0.0, 0.5)$ distribution.

3. FBD model with a fixed mass extinction time of 55 Ma, while treating the survival probability as an unknown parameter. The prior for the survival probability was $\text{Uniform}(0.0, 0.5)$.

###### S5.2.6 Convergence of the Inference

We ran eight independent MCMC chains for 40,000 iterations each for Models 1 and 3 (requiring 40–50 days of computation time per chain), and two chains for 20,000 iterations for Model 2. We did not extend the runs for Model 2 for a different reason: the Bayes Factor results decisively rejected the inclusion of a mass extinction, making its posterior estimates redundant with those of Model 1. Therefore, further sampling was scientifically unnecessary. After a 25% burn-in, we assessed convergence for all runs. While the independent chains consistently converged on the same posterior distribution and the ESS for the tree topology exceeded 100, some scalar parameters remained below the conventional threshold of 200. Given the consistency across chains and the substantial computational investment required for marginal improvements to the ESS of all parameters, we concluded these runs were sufficient for our primary comparisons.

###### S5.2.7 Detection of Mass Extinction

We found no evidence for a mass extinction event when explicitly testing for it under Model 2. Two independent chains yielded decisive 2 log Bayes Factors of -9.4 and -11.4 against the presence of a mass extinction event.

##### S5.2.8 Comparison of Models

Because the posterior estimates from Model 2 were consequently congruent with those from the no-extinction model (Model 1), we focus our subsequent discussion on the comparison between Model 1 and Model 3. This comparison is informative as Model 3 assumes a mass extinction event is a fixed component of the model, which allows for a more direct assessment of the impact of such an event on phylogenetic inference.

The topologies (Figures S40 and S41) and divergence times (Figures 8 and S35 to S37) of the MCC extant and full trees inferred from total-evidence dating were highly similar, regardless of whether the FBD model without mass extinction (Model 1) or with a mass extinction event (Model 3) was employed as the tree prior. The differences between the estimates from the two models were comparable to the variance among different MCMC chains of the same model (Figures S38-S41), suggesting that the choice of FBD tree prior has a negligible impact on the phylogenetic inference.

### S6 Crinoids Analysis

We re-analyzed the total-evidence dataset for crinoids from Cole et al. (2025). This dataset is particularly suitable for our study as its fossil sampling spans the Ordovician and Silurian periods. This geological timeframe encompasses the Late Ordovician Mass Extinction, the first of the five major extinction events of the Phanerozoic Eon. The occurrence of this major biotic crisis within the sampling window suggests that crinoids may have been significantly impacted.

#### S6.1 Data

We used the dataset from Cole et al. (2025). The dataset consisted of 42 fossil taxa from the Ordovician and Silurian periods, each with a corresponding temporal age range. The character matrix was composed of 25 morphological characters.

#### S6.2 Models

Phylogenetic inference was performed using the total-evidence dating method implemented in `RevBayes` (Höhna et al., 2016), with additional comparative analyses conducted in BEAST2 and MrBayes. For the MrBayes analyses, we used the configuration files from Cole et al. (2025) but conducted new runs, as they provided these files rather than the final result files.

##### S6.2.1 Morphological Substitution Model

We modeled the evolution of the morphological characters using the Mk model, where the number of states (k) was set to the maximum number of states observed across all characters in the dataset. To correct for the ascertainment bias that can arise from analyzing datasets that exclude constant characters, we applied the Mkv model variant in all three software packages.

##### S6.2.2 Morphological Clock Model

In RevBayes and BEAST2, we modeled the evolution of the morphological data using the UCLN relaxed clock model. The prior for the mean evolutionary rate was an exponential distribution with a rate of 10. The prior for the standard deviation of the lognormal distribution was an exponential distribution with a rate of 10/3. MrBayes used the analogous uncorrelated gamma relaxed clock model

##### S6.2.3 Tree Model

We used three constant-rate FBD models as tree priors across our analyses. In our primary analyses (conducted in RevBayes), we placed direct exponential priors on the speciation, extinction, and fossilization rates, each with a rate of 10. For the comparative analyses, MrBayes and BEAST2 used the alternative parameterization required by those programs, setting priors on net diversification, turnover, and sampling proportion. We opted for the direct parameterization with speciation, extinction, and fossilization rates for two main reasons: first, to maintain consistency with our simulations, and second, because the configuration in MrBayes requires the net diversification rate to be positive, which forces the speciation rate to exceed the extinction rate. These comparative analyses were limited to Model 1, as the Model 2 and Model 3 frameworks are not currently supported in MrBayes or BEAST2. The sampling probability of extant species ($\rho$) in MrBayes and BEAST2 was fixed to 1.0, whereas in RevBayes it could only be set to 0 for a clade that does not include extant species. The three FBD models were:

1. FBD model without mass extinction.

2. FBD model with a fixed mass extinction time of 443.8 Ma, while treating the survival probability as an unknown parameter. Following the third model in Section 2.2.1, the prior for the survival probability was defined using a reversible-jump mixture model framework. An equal prior probability of 0.5 was assigned to the models with and without a mass extinction event. In the model that includes a mass extinction, the survival probability is treated as an unknown parameter drawn from a Uniform$(0.3, 0.6)$ distribution.

3. FBD model with a fixed mass extinction time of 443.8 Ma, while treating the survival probability as an unknown parameter. The prior for the survival probability was Uniform$(0.0, 0.6)$.

##### S6.2.4 Convergence of the Inference

We ran two independent MCMC chains for 1,000,000 iterations each for both Models. While the independent chains consistently converged on the same posterior distribution and the ESS for the tree topology exceeded 200, some scalar parameters remained below the conventional threshold of 200. Given the consistency across chains and the substantial computational investment required for marginal improvements to the ESS of all parameters, we concluded these runs were sufficient for our primary comparisons.

##### S6.2.5 Fossil Species

The inference of sampled ancestors varied substantially across the three softwares. The posterior mean number of sampled ancestors was approximately 28 in RevBayes, 20 in BEAST2, and 17 in MrBayes. We defined "fossil species" as those fossils rejected as sampled ancestors across all chains and all software analyses, yielding a set of 9 such species. For context, under the no-mass-extinction model (Model 1) in RevBayes, one chain yielded only 10 fossil species, indicating our set of 9 is representative of the RevBayes results.

##### S6.2.6 Detection of Mass Extinction

We found no evidence for a mass extinction event when explicitly testing for it under Model 2. Two independent chains yielded decisive 2 log Bayes Factors of -1.8 and -1.8 against the presence of a mass extinction event.

##### S6.2.7 Comparison of Models

Because the posterior estimates from Model 2 were consequently congruent with those from the no-extinction model (Model 1), we focus our subsequent discussion on the comparison between Model 1 and Model 3. This comparison is informative as Model 3 assumes a mass extinction event is a fixed component of the model, which allows for a more direct assessment of the impact of such an event on phylogenetic inference.

The topologies (Figures S49 and S50) and divergence times (Figures 8 and S42 to S44) of the MCC fossil species and full trees inferred from total-evidence dating were highly similar, regardless of whether the FBD model without mass extinction (Model 1) or with a mass extinction event (Model 3) was employed as the tree prior. The estimated fossil ages (Figure S45) and the probabilities of fossils as sampled ancestors (Figure S46) were also comparable between the two FBD model configurations. The differences between the estimates from the two models were comparable to the variance among different MCMC chains of the same model (Figures S47-S50), suggesting that the choice of FBD tree prior has a negligible impact on the phylogenetic inference.

When no mass extinction was modeled, the phylogenetic trees and fossil age estimates produced by the three software packages were likewise highly congruent.

Table S1: Maximum number of MCMC iterations run for different scenarios in total-evidence dating and the number of simulated trees (out of 100) with an ESS greater than 200. For the definitions of “ME”, “WME”, “$c$”, “$\psi$”, “prior”, and “strong prior”, please refer to Figure 1.

| Clock | Simulation | Inference | $\psi$ | Survival Probability of Mass Extinction | Prior | $c$ | Maximum Number of Iteration | Number of Trees with ESS > 200 |
|---|---|---|---|---|---|---|---|---|
| Strict | ME | ME | 0.03 | 0.2 | - | 250 | 200000 | 87 |
| Strict | ME | ME | 0.03 | unknown | 0.1 | 250 | 200000 | 90 |
| Strict | ME | ME | 0.03 | unknown | 0.5 | 250 | 200000 | 90 |
| Strict | ME | ME | 0.03 | unknown | 0.5 | 25 | 200000 | 91 |
| Strict | ME | ME | 0.03 | unknown | 0.9 | 250 | 200000 | 92 |
| Strict | ME | ME | 0.015 | 0.2 | - | 250 | 400000 | 96 |
| Strict | ME | ME | 0.015 | unknown | 0.5 | 250 | 400000 | 94 |
| Strict | ME | WME | 0.03 | 1 | - | 250 | 200000 | 89 |
| Strict | ME | WME | 0.03 | 1 | - | 25 | 200000 | 91 |
| Strict | ME | WME | 0.015 | 1 | - | 250 | 400000 | 98 |
| Strict | ME | ME | 0.03 | unknown | 0.5, strong prior | 250 | 200000 | 91 |
| UCLN | ME | ME | 0.03 | unknown | 0.5 | 250 | 1000000 | 96 |
| UCLN | ME | WME | 0.03 | 1 | - | 250 | 1000000 | 89 |
| Strict | WME | ME | 0.03 | unknown | 0.1 | 250 | 200000 | 88 |
| Strict | WME | ME | 0.03 | unknown | 0.5 | 250 | 200000 | 83 |
| Strict | WME | ME | 0.03 | unknown | 0.5 | 25 | 200000 | 82 |
| Strict | WME | ME | 0.03 | unknown | 0.9 | 250 | 200000 | 89 |
| Strict | WME | ME | 0.015 | unknown | 0.5 | 250 | 400000 | 85 |
| Strict | WME | WME | 0.03 | 1 | - | 250 | 200000 | 86 |
| Strict | WME | WME | 0.03 | 1 | - | 25 | 200000 | 87 |
| Strict | WME | ME | 0.03 | unknown | 0.5, strong prior | 250 | 200000 | 91 |
| UCLN | WME | ME | 0.03 | unknown | 0.5 | 250 | 1000000 | 97 |
| UCLN | WME | WME | 0.03 | 1 | - | 250 | 1000000 | 95 |

Table S2: Maximum number of MCMC iterations run for different scenarios in MCC tree and the number of simulated trees (out of 100) with an ESS greater than 200. For the definitions of "inferred by estimating survival probability of mass extinction" and "inferred without mass extinction", please refer to Figure 6.

| Inference | Maximum Number of Iteration | Number of Trees with ESS $>$ 200 |
|---|---|---|
| Inferred by estimating survival probability of mass extinction | 450000 | 90 |
| Inferred without mass extinction | 450000 | 89 |

Table S3: Quantification of Simulated Trees Excluded by Filtering Criteria. For each of the four main simulation scenarios, 1,000 trees were initially simulated. The table shows the number of trees removed by the origin time filter, followed by the number of trees removed by the fossil count filter from the remaining pool.

| Mass Extinction | $\psi$ | Origin Time $<$ Min | Origin Time $>$ Max | Fossils $<$ Min | Fossils $>$ Max |
|---|---|---|---|---|---|
| Without | 0.015 | 22 | 31 | 25 | 28 |
| With | 0.015 | 21 | 36 | 22 | 56 |
| Without | 0.03 | 33 | 20 | 17 | 26 |
| With | 0.03 | 21 | 36 | 37 | 56 |

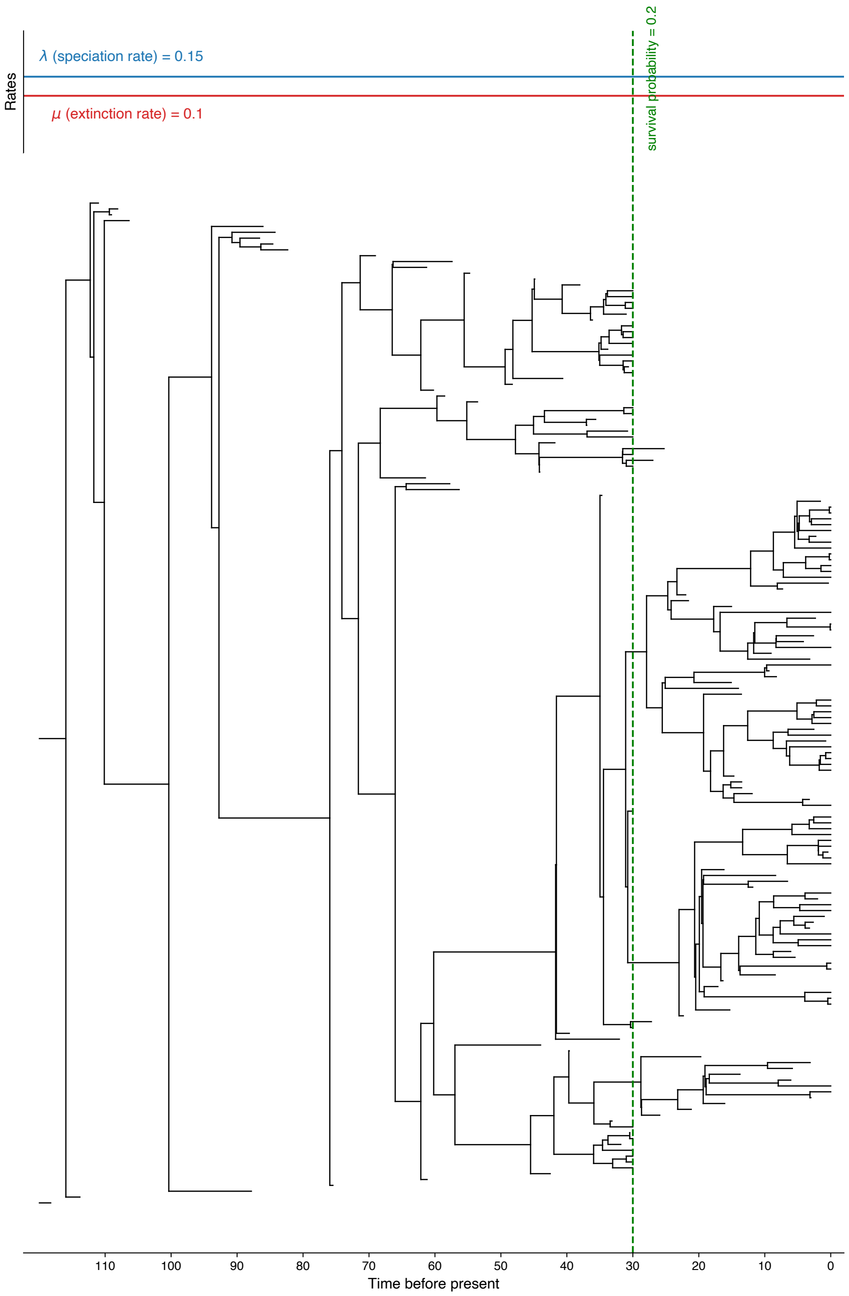


Figure S1: A schematic illustration of a simulated complete phylogenetic tree subjected to a mass extinction event, adapted from Culshaw et al. (2019). The phylogeny evolves under constant background rates of speciation ($\lambda = 0.15$) and extinction ($\mu = 0.1$). The green vertical line at 30 time units before the present marks the mass extinction event, which is modeled using a Dirac delta function, where each extant lineage has a survival probability of 0.2.

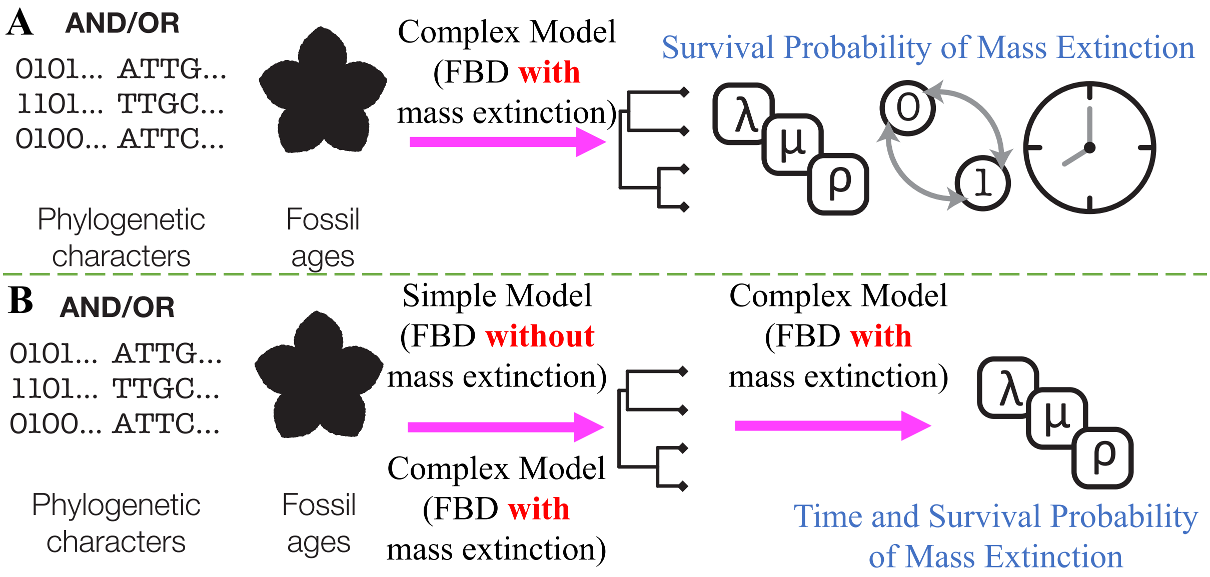


Figure S2: The two strategies used to infer mass extinction. (A) The one-step approach. The FBD model with mass extinction is used as the tree prior to jointly infer the phylogeny and the survival probability of the mass extinction. (B) The two-step approach. First, an MCC tree is inferred using either the FBD model with or without mass extinction as the tree prior. Second, the time and survival probability of the mass extinction are inferred on the fixed MCC tree.

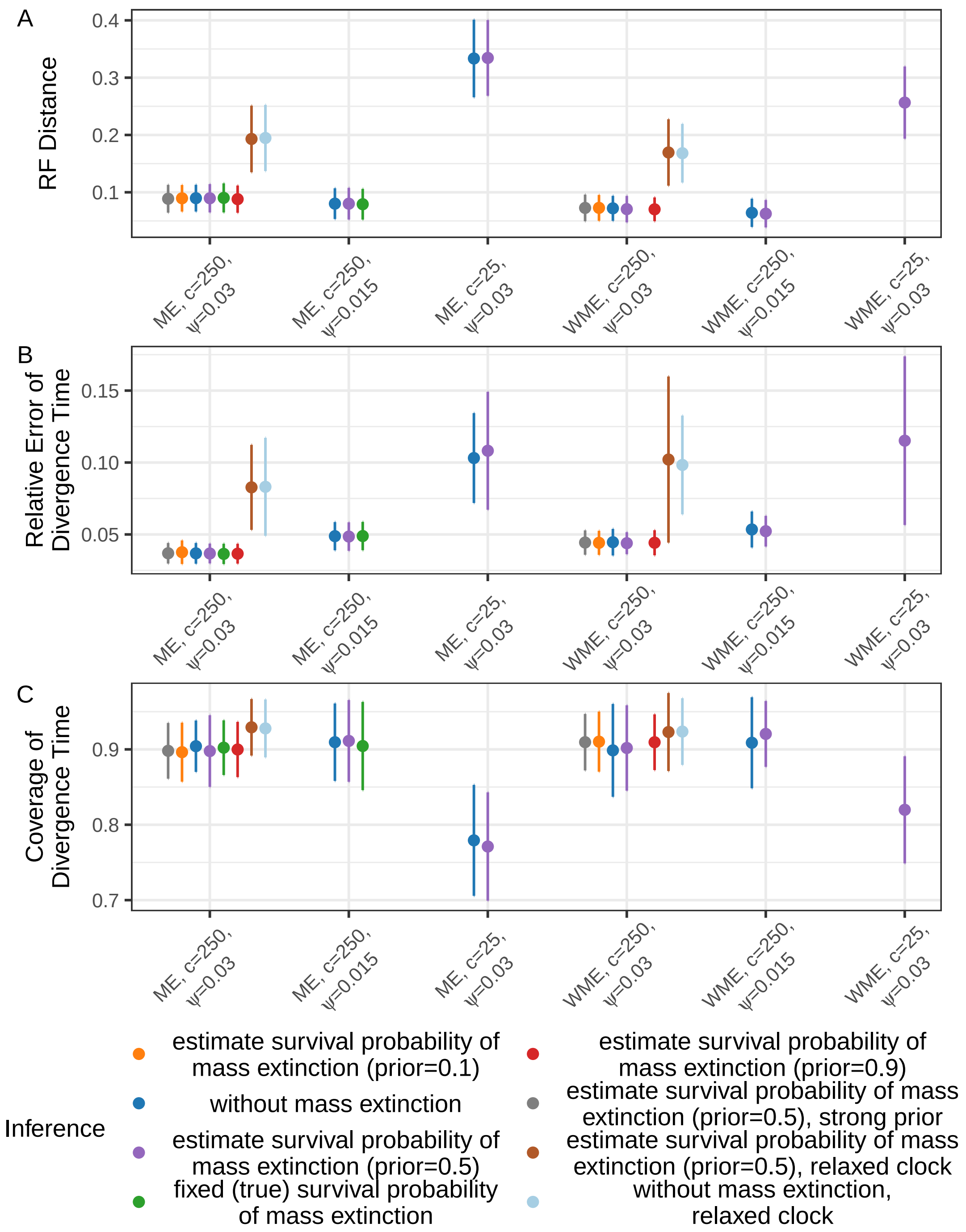


Figure S3: RF distance, relative error of divergence time, and coverage of divergence time between the inferred and true sampled tree with fossils when using different FBD models as tree priors. The points and bars represent the mean and standard deviation across all simulated trees. “fixed (true) survival probability of mass extinction” refers to FBD model with known (true) time and survival probability of mass extinction, “estimate survival probability of mass extinction”, refers to FBD model with known time but unknown survival probability of mass extinction, “without mass extinction” refers to FBD model without mass extinction. For the definitions of “ME”, “$c$”, “$\psi$”, “prior”, and “strong prior”, please refer to Figure 1.

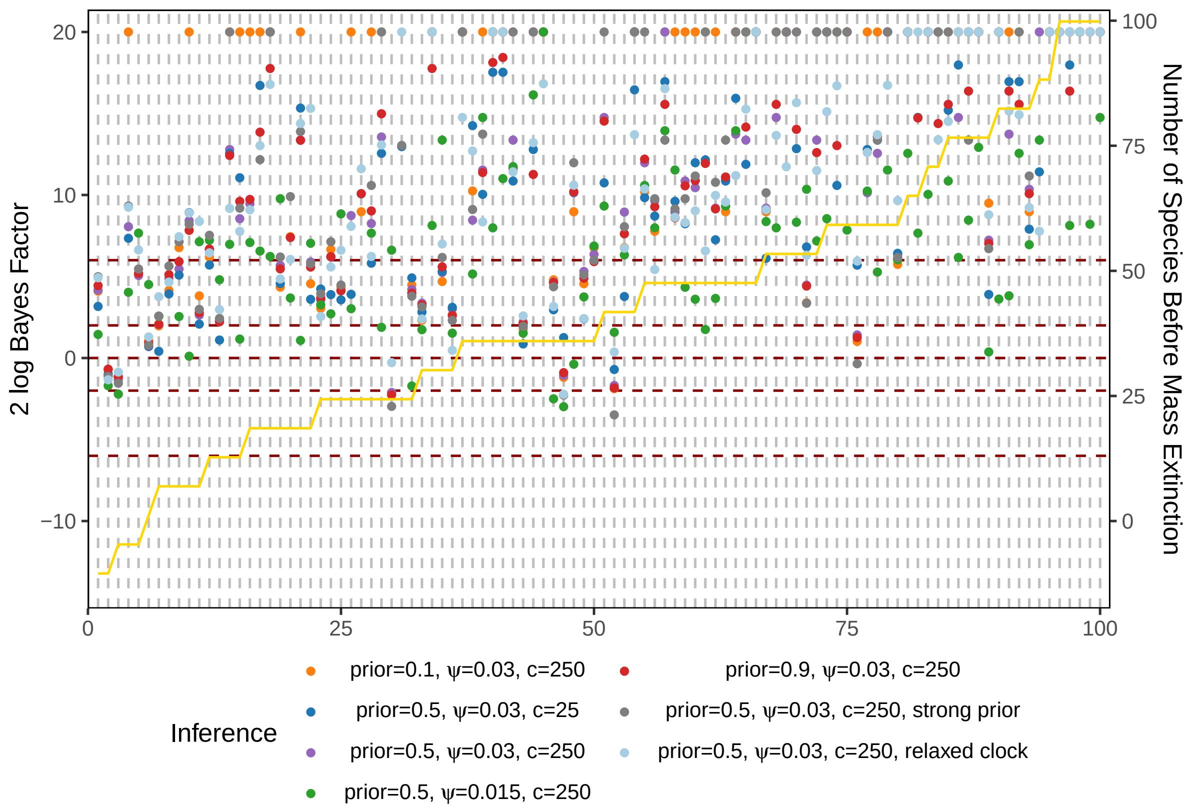


Figure S4: Bayes factors for each simulated tree under different scenarios in total-evidence dating when a mass extinction is present. The x-axis represents 100 simulated trees, sorted from left to right by the number of species before the mass extinction (from smallest to largest). The gold line indicates the number of species before the mass extinction. For the definitions of "$c$", "$\psi$", "prior", and "strong prior", please refer to Figure 1.

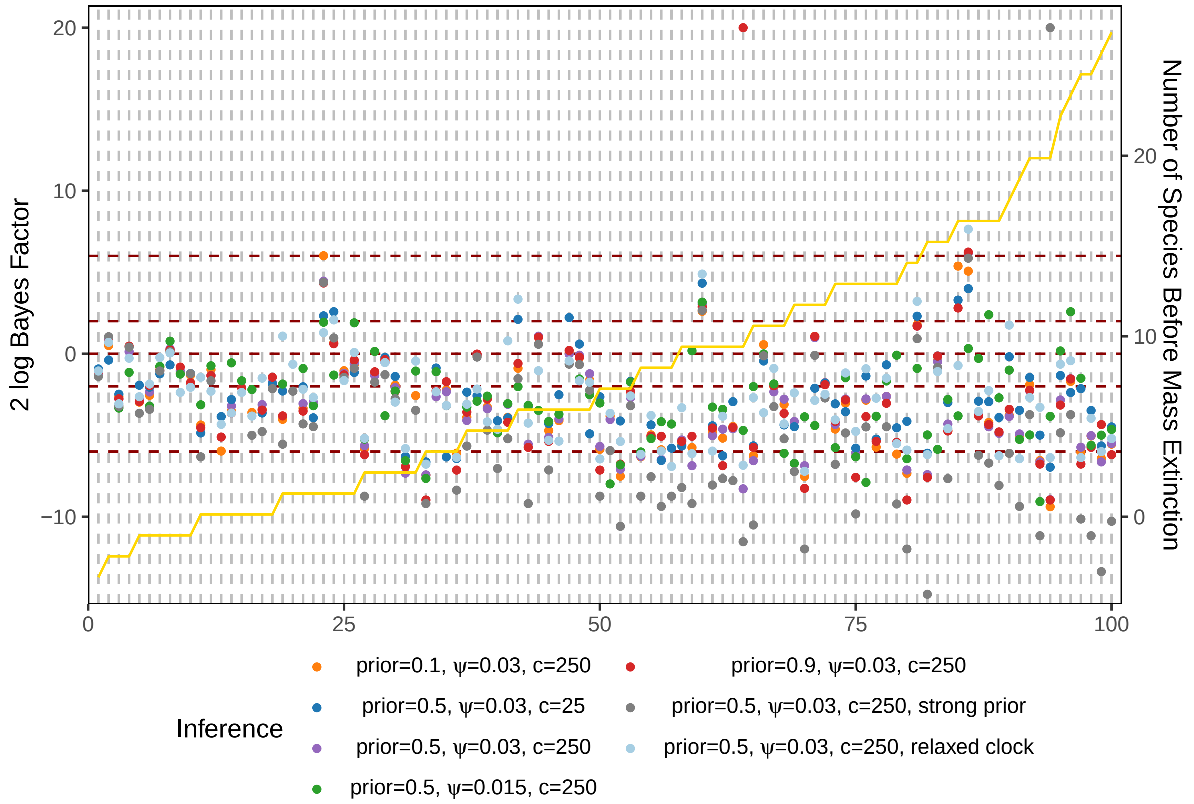


Figure S5: Bayes factors for each simulated tree under different scenarios in total-evidence dating when a mass extinction is absent. The x-axis represents 100 simulated trees, sorted from left to right by the number of species before the mass extinction (set at 30Ma, the mass extinction time during inference, despite no actual mass extinction occurring) (from smallest to largest). The gold line indicates the number of species before the mass extinction. For the definitions of "$c$", "$\psi$", "prior", and "strong prior", please refer to Figure 1.

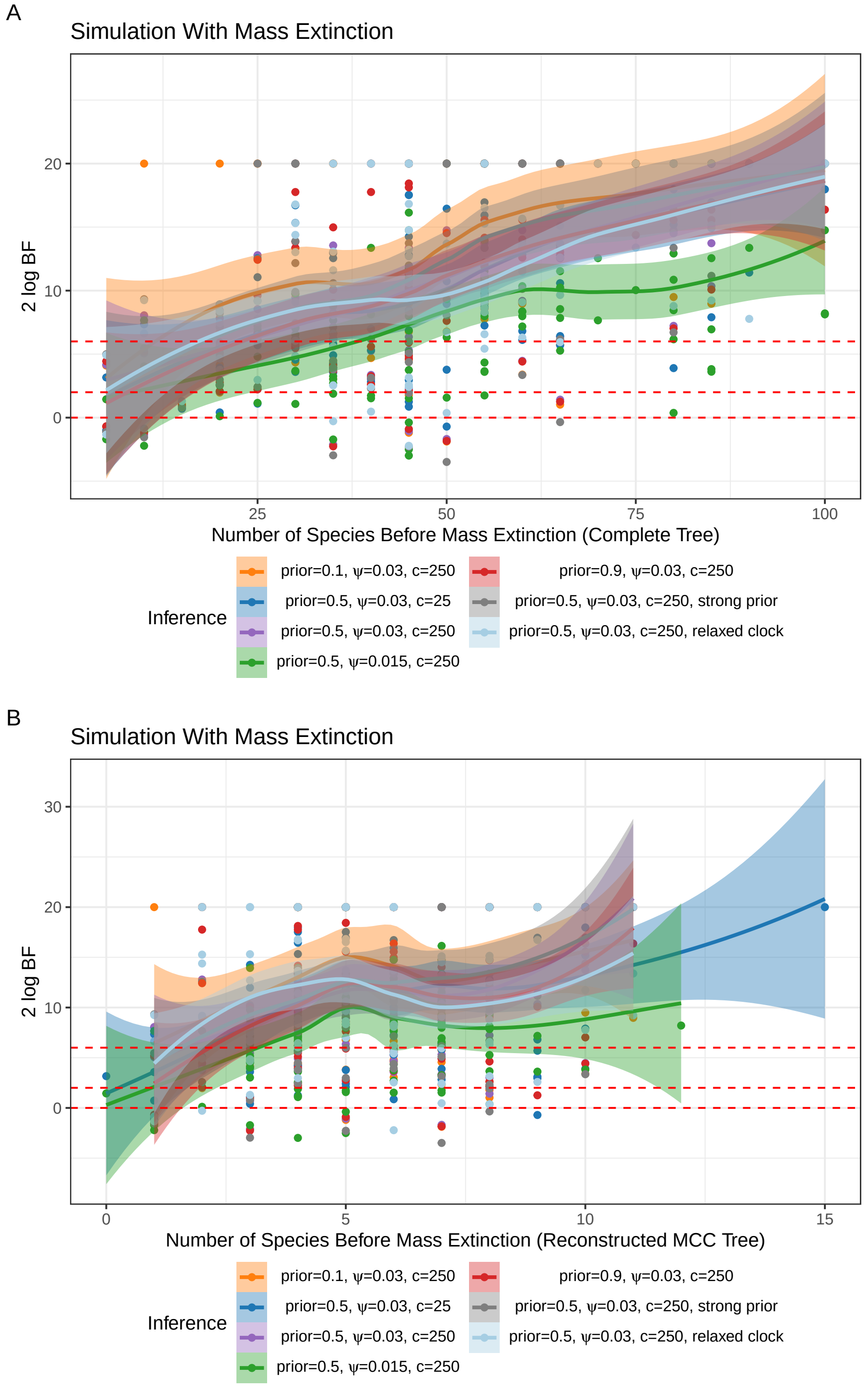


Figure S6: The relationship between Bayes factors and the number of species before the mass extinction under different scenarios when a mass extinction is present for (A) the complete trees and (B) the reconstructed MCC trees. Each point represents a single simulated tree. The curves and shaded regions were fitted using the default parameters of the `geom_smooth` function in R package `ggplot2`. For the definitions of "$c$", "$\psi$", "prior", and "strong prior", please refer to Figure 1.

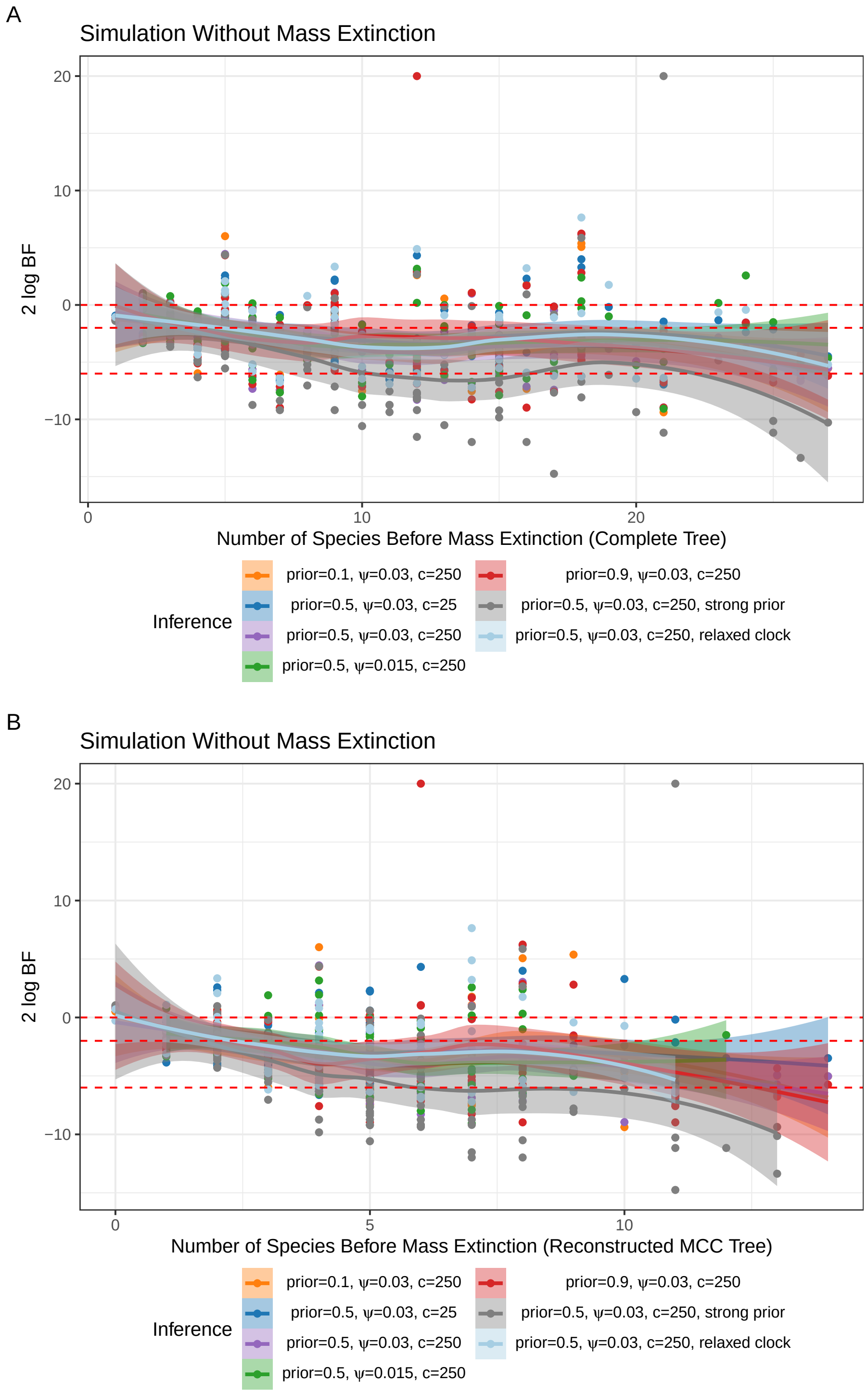


Figure S7: The relationship between Bayes factors and the number of species before the mass extinction (30Ma) under different scenarios when a mass extinction is absent for (A) the complete trees and (B) the reconstructed MCC trees. Each point represents a single simulated tree. The curves and shaded regions were fitted using the default parameters of the `geom_smooth` function in R package `ggplot2`. For the definitions of "$c$", "$\psi$", "prior", and "strong prior", please refer to Figure 1.

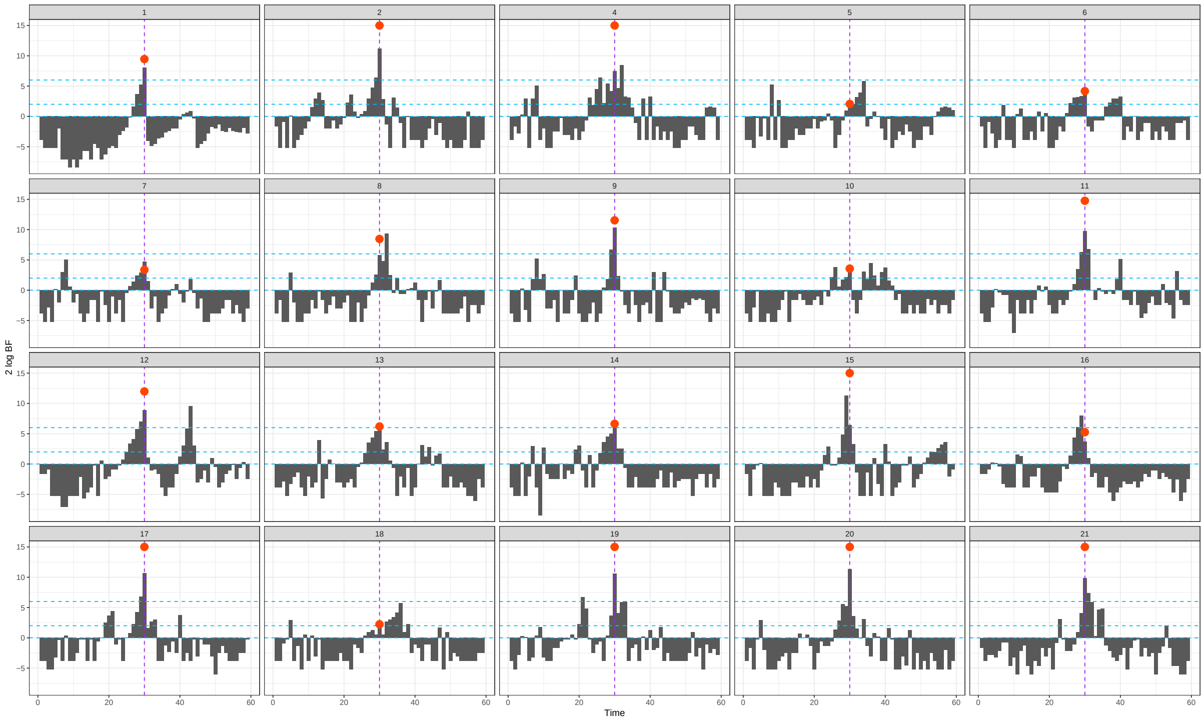


Figure S8: Bayes factors for each possible mass extinction time across all simulated trees, inferred using an FBD model with unknown time and survival probability of mass extinction in total-evidence dating. The blue horizontal lines indicate thresholds of $2 \log \mathrm{BF}_{\mathrm{ME,WME}}$ at 0, 2, and 6, while the purple vertical line marks the true extinction time (30Ma). Red dots represent Bayes factors obtained using an FBD model with known mass extinction time but unknown mass extinction survival probability. The different panels correspond to different simulation replicates.

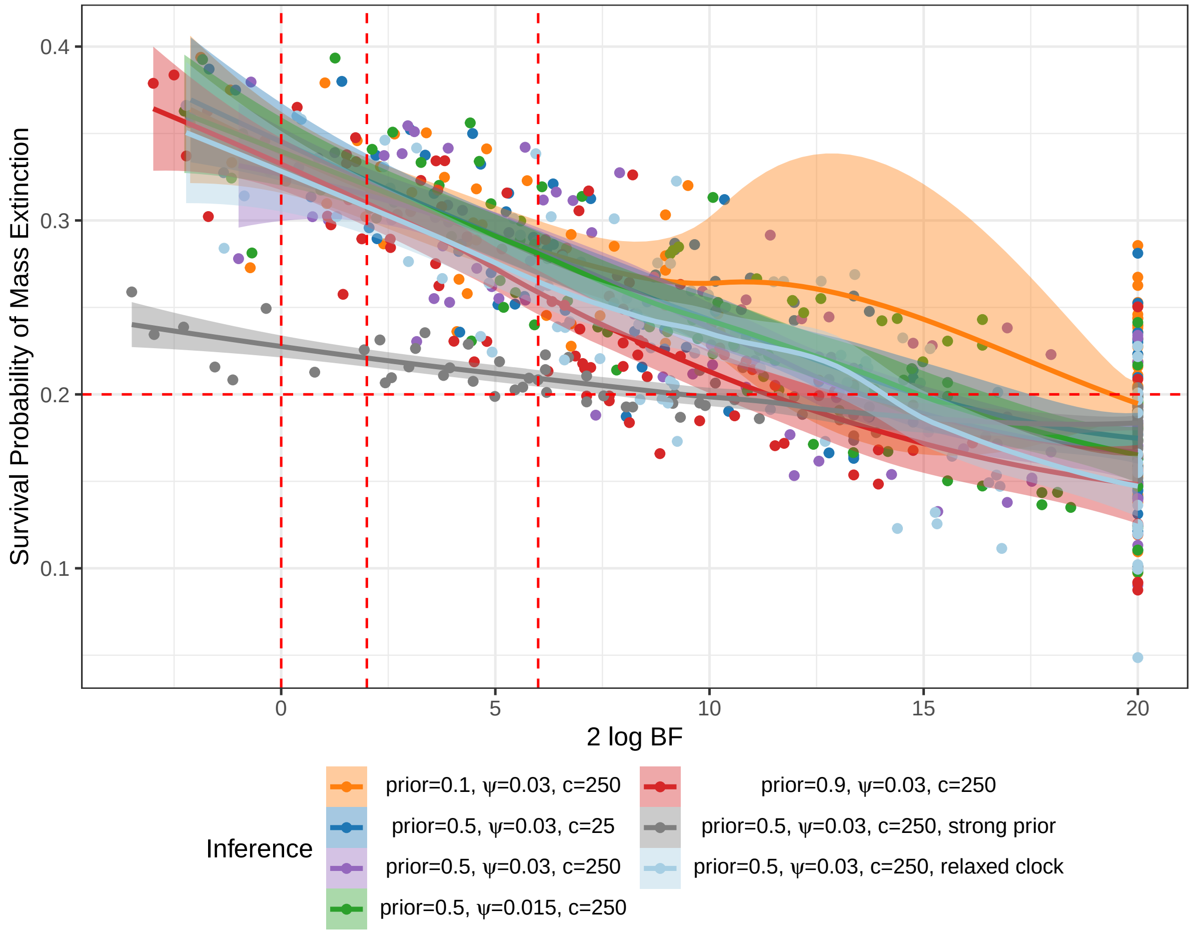


Figure S9: The relationship between inferred survival probability of mass extinction and Bayes factors under different scenarios when a mass extinction is present. Each point represents a single simulated tree. The curves and shaded regions were fitted using the default parameters of the `geom_smooth` function in R package `ggplot2`. For the definitions of "$c$", "$\psi$", "prior", and "strong prior", please refer to Figure 1.

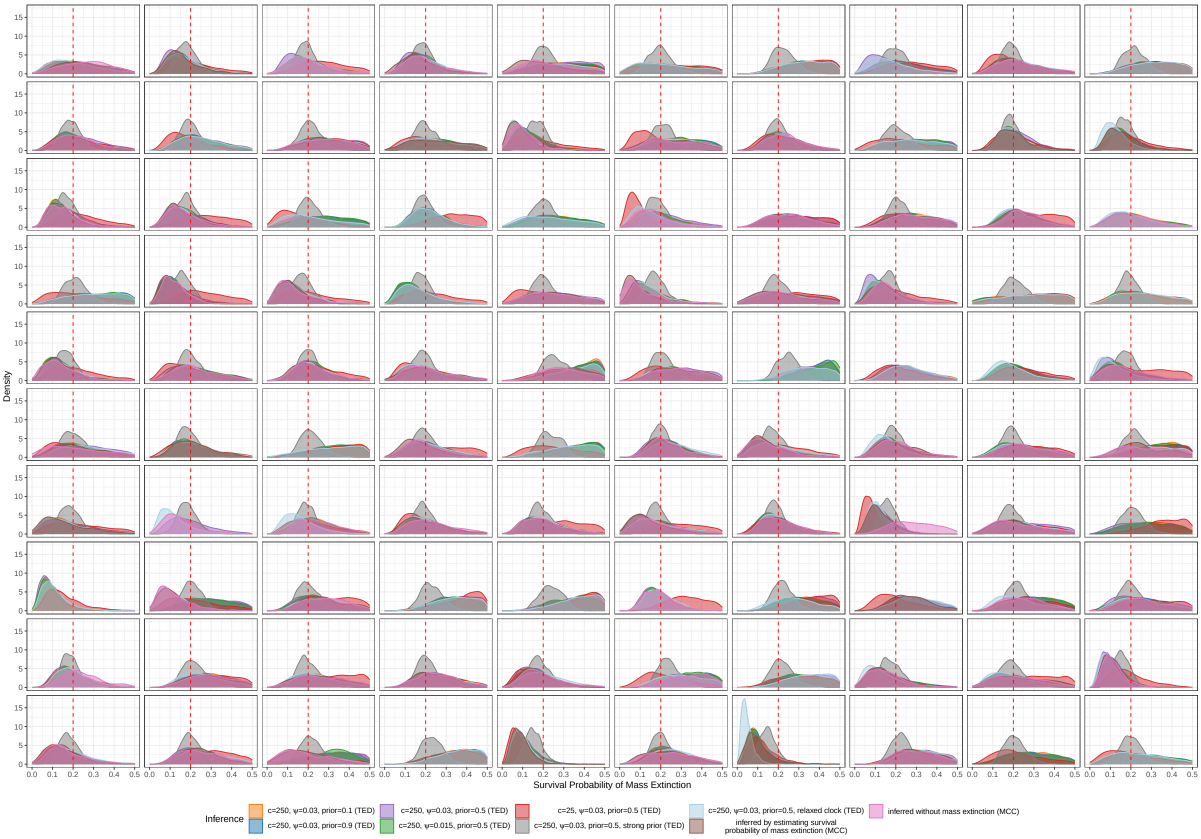


Figure S10: Posterior probability density plots of mass extinction survival probabilities for each simulated trees, obtained using different methods. For the definitions of "$c$", "$\psi$", "prior", and "strong prior", please refer to Figure 1.

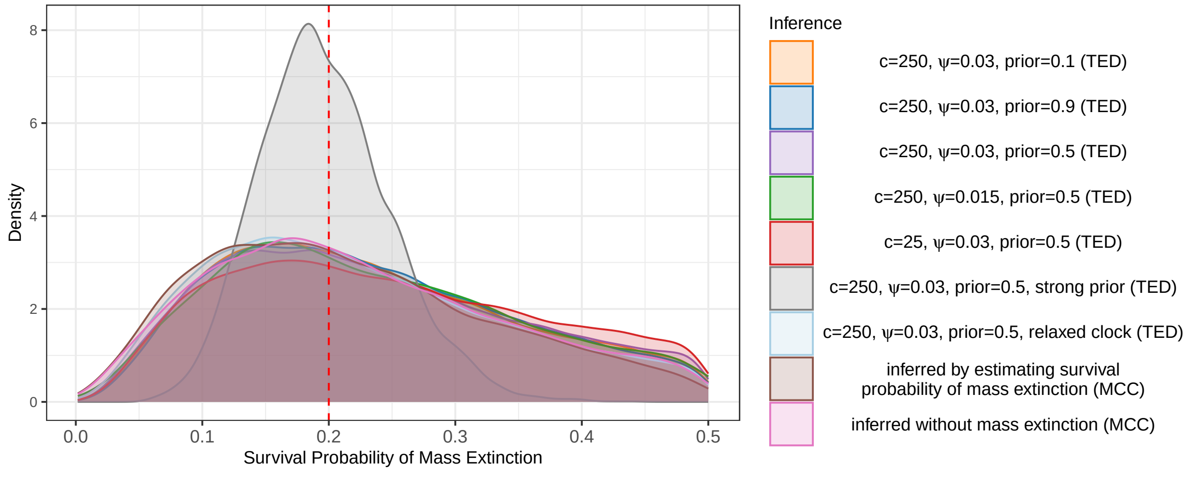


Figure S11: Posterior probability density plots of mass extinction survival probabilities for all simulated trees, obtained using different methods. Similar to Figure 3, but this figure includes only trees that converged across all methods and detected a single mass extinction event in the MCC tree.

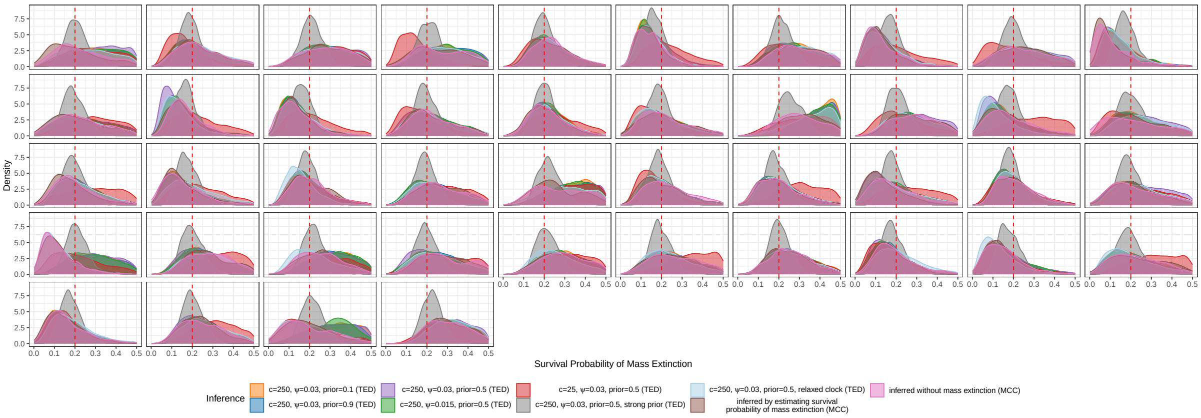


Figure S12: Posterior probability density plots of mass extinction survival probabilities for each simulated trees, obtained using different methods. Similar to Figure S11, but this figure includes only trees that converged across all methods and detected a single mass extinction event in the MCC tree. For the definitions of "$c$", "$\psi$", "prior", and "strong prior", please refer to Figure 1.

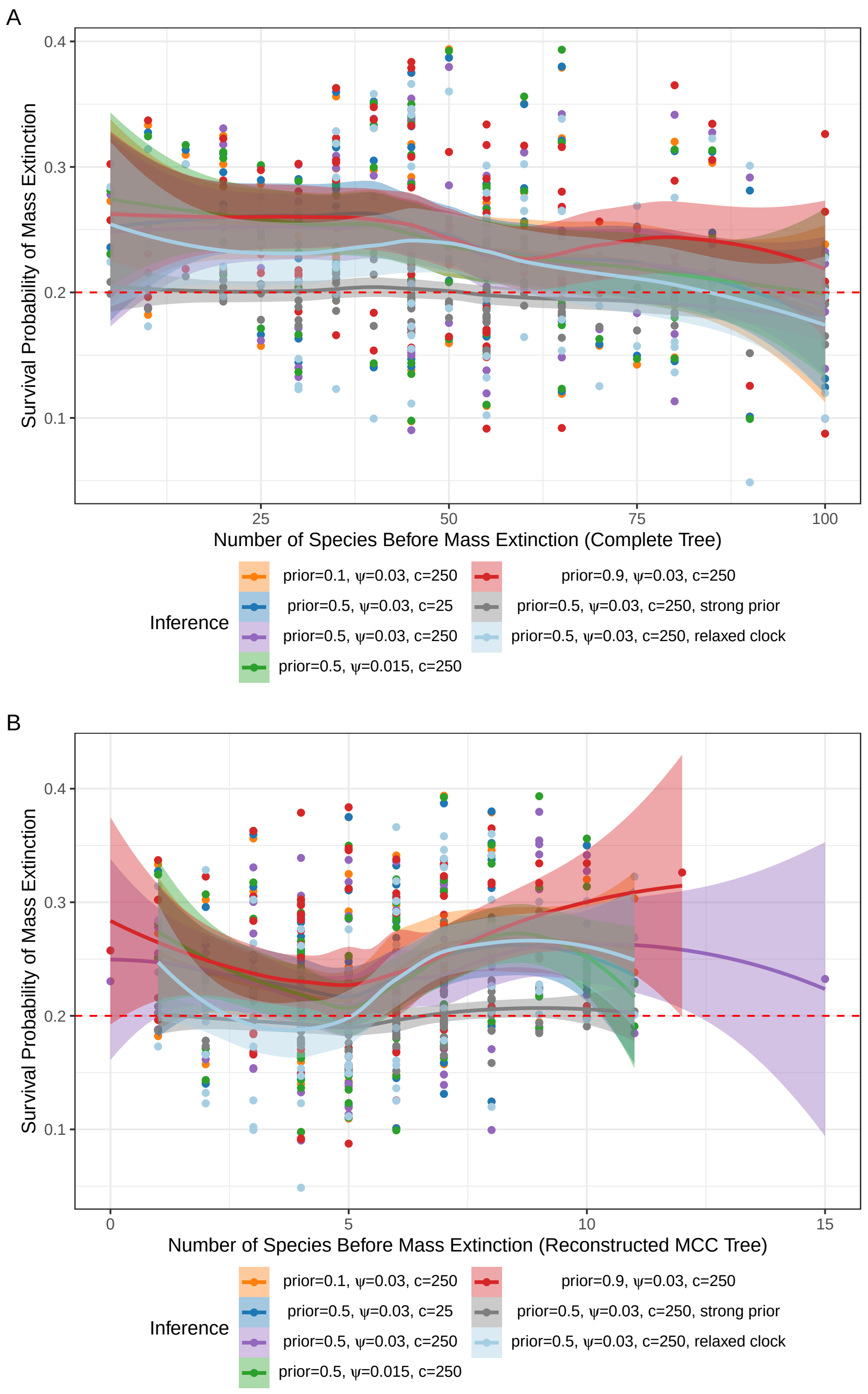


Figure S13: The relationship between inferred survival probability of mass extinction and the number of species before the mass extinction (30Ma) under different scenarios when a mass extinction is present for (A) the complete trees and (B) the reconstructed MCC trees. Each point represents a single simulated tree. The curves and shaded regions were fitted using the default parameters of the `geom_smooth` function in R package `ggplot2`. For the definitions of "$c$", "$\psi$", "prior", and "strong prior", please refer to Figure 1.

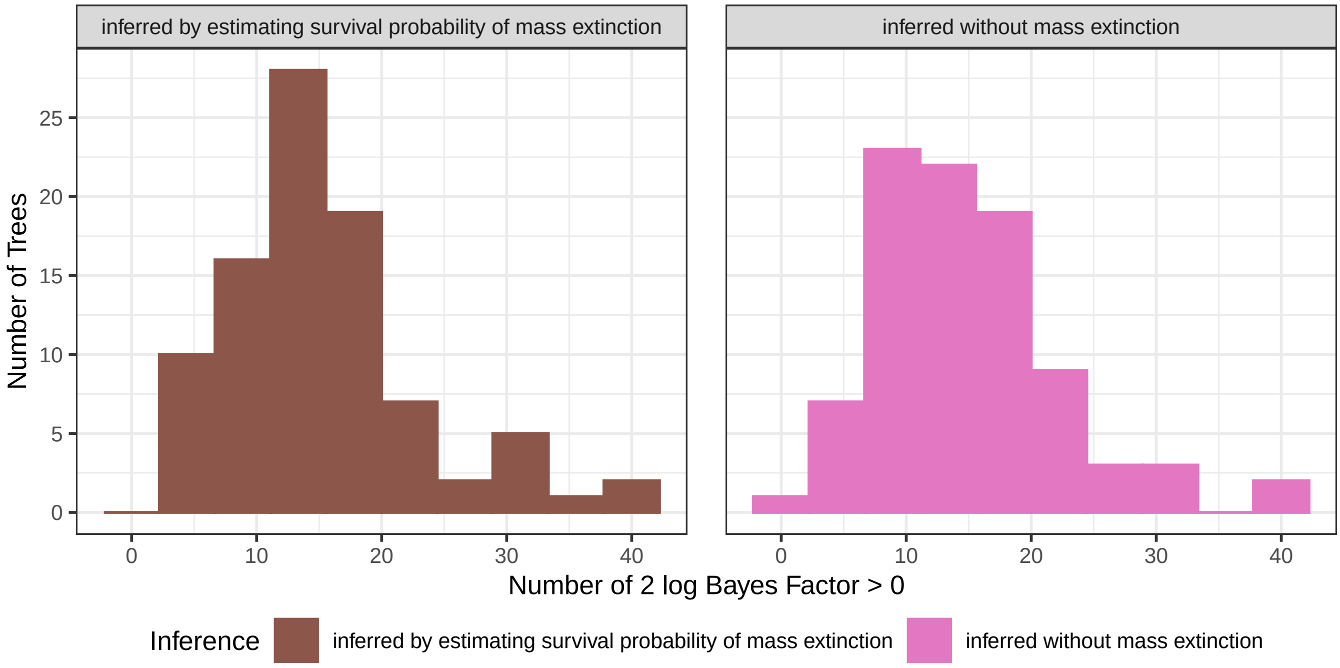


Figure S14: Histogram showing the number of time points with Bayes factors greater than 0 for each MCC tree. For the definitions of "inferred by estimating survival probability of mass extinction" and "inferred without mass extinction", please refer to Figure 6.

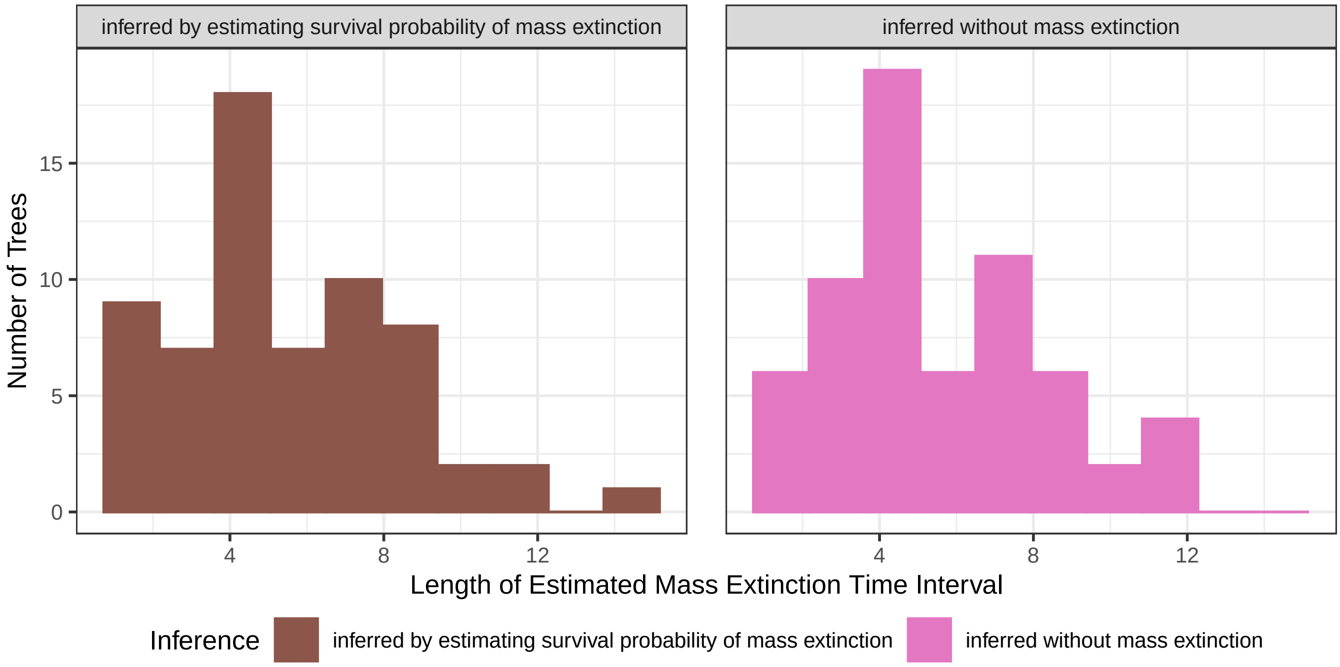


Figure S15: Histogram of the length of the estimated mass extinction time intervals for a single mass extinction event detected in the MCC tree. The length of the estimated intervals corresponds to the line lengths shown in Figure 7. For the definitions of "inferred by estimating survival probability of mass extinction" and "inferred without mass extinction", please refer to Figure 6.

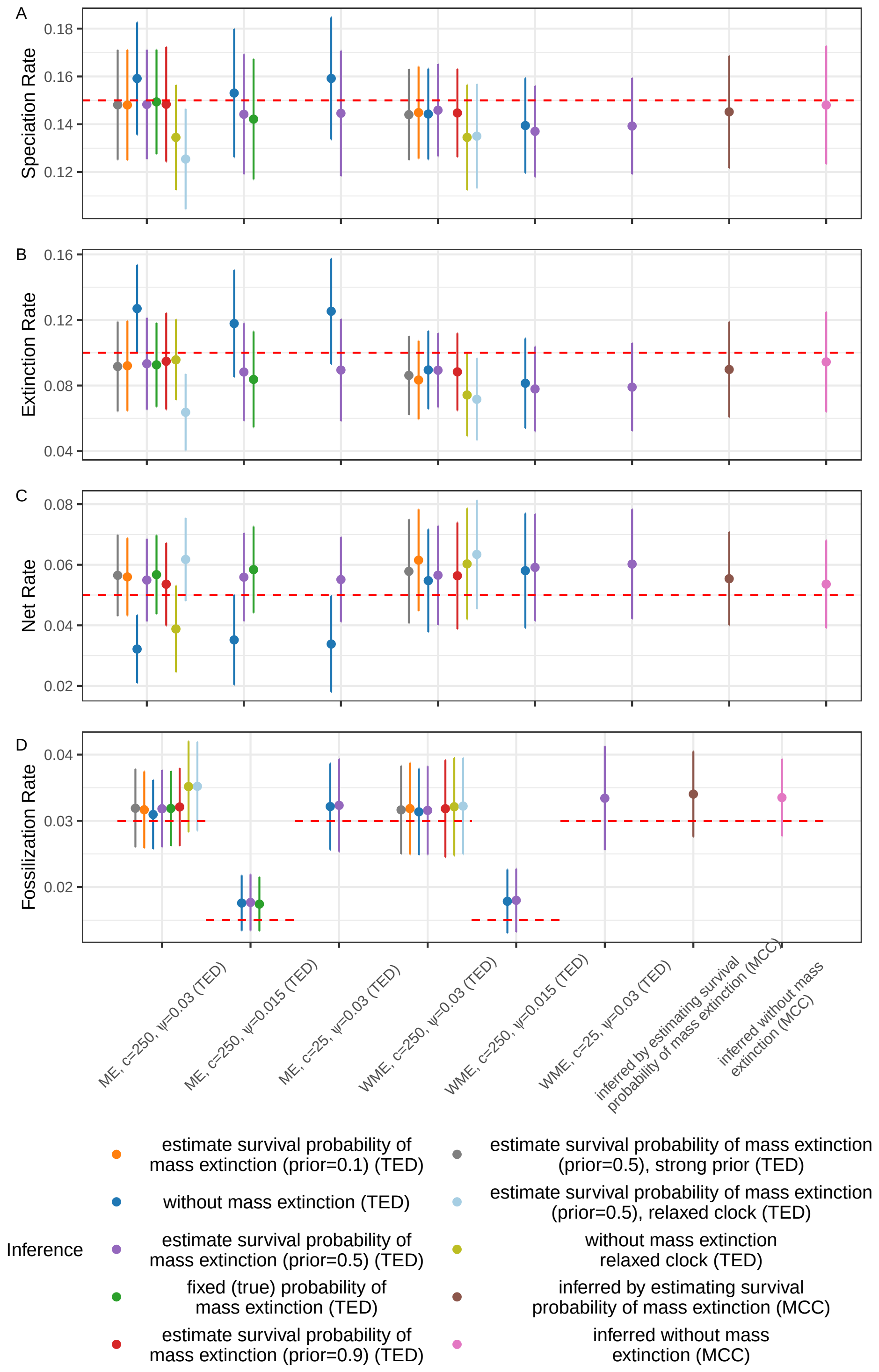


Figure S16: Speciation, extinction, net diversification, and fossilization rates inferred using different methods. The points and bars represent the mean and standard deviation across all simulated trees. The red horizontal lines indicate true value. For the definitions of “TED”, “MCC”, “$c$”, “$\psi$”, “prior”, and “strong prior”, please refer to Figure 3. For the definitions of “inferred by estimating survival probability of mass extinction” and “inferred without mass extinction”, please refer to Figure 6.

.

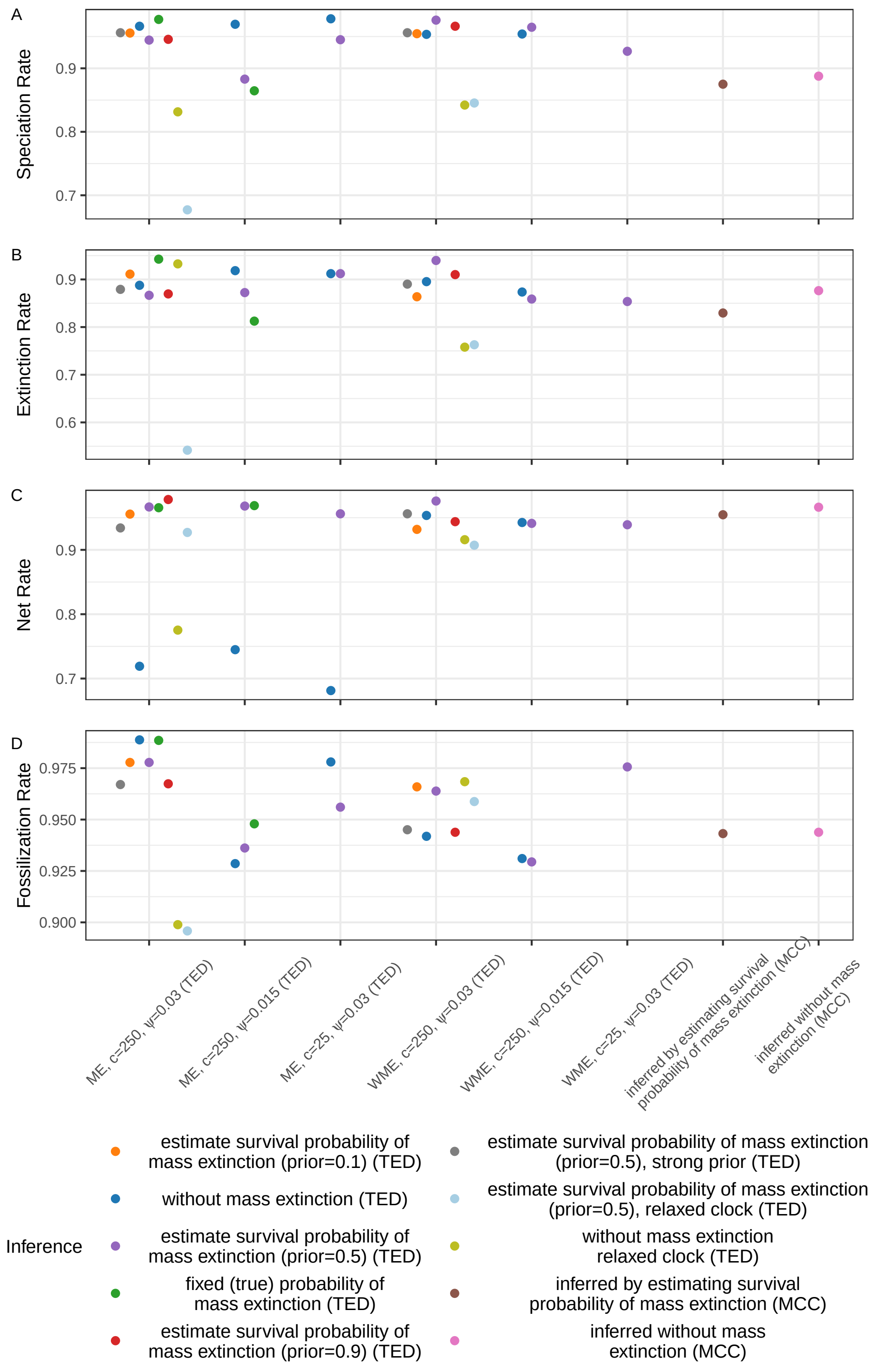


Figure S17: Coverage of speciation, extinction, net diversification, and fossilization rates inferred using different methods. The red horizontal lines indicate true value. For the definitions of “TED”, “MCC”, “$c$”, “$\psi$”, “prior”, and “strong prior”, please refer to Figure 3. For the definitions of “inferred by estimating survival probability of mass extinction” and “inferred without mass extinction”, please refer to Figure 6.

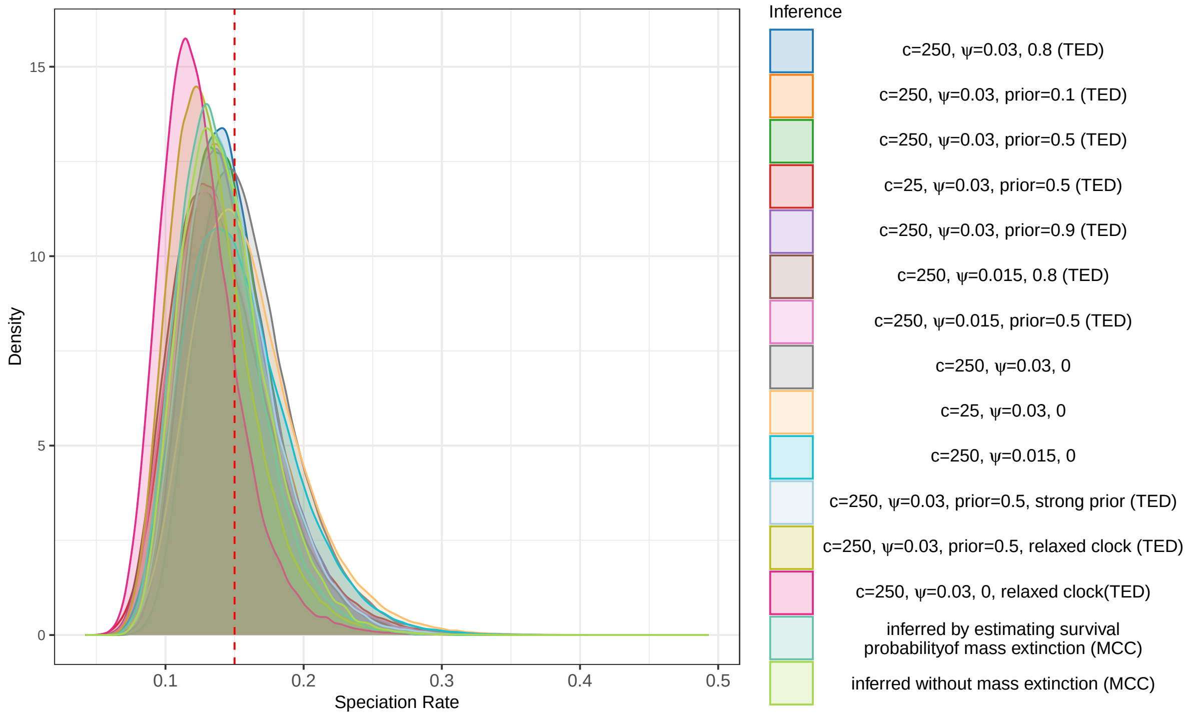


Figure S18: Posterior probability density plots of speciation rate for all simulated trees, obtained using different methods. For details on how the probability density curves were generated, please refer to Figure 2. The red vertical line indicates the true speciation rate (0.15). For the definitions of "TED", "MCC", "$c$", "$\psi$", "prior", and "strong prior", please refer to Figure 3. For the definitions of "inferred by estimating survival probability of mass extinction" and "inferred without mass extinction", please refer to Figure 6.

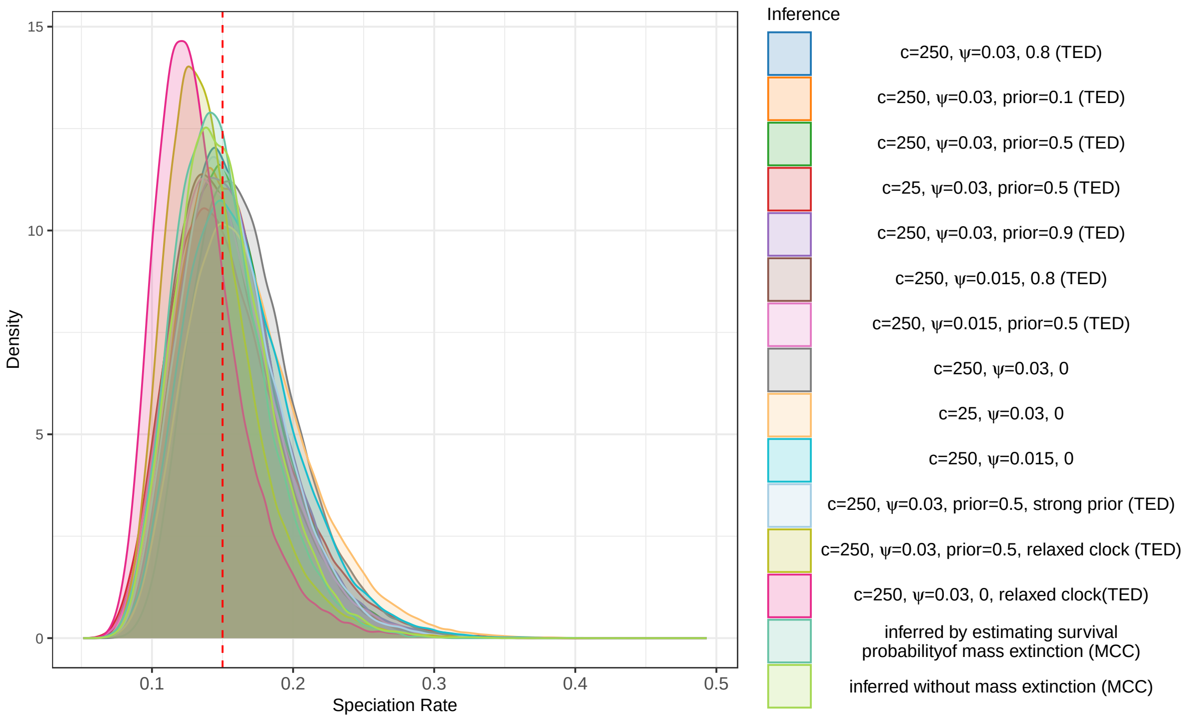


Figure S19: Posterior probability density plots of speciation rate for all simulated trees, obtained using different methods. Similar to Figure S18, but this figure includes only trees that converged across all methods and detected a single mass extinction event in the MCC tree.

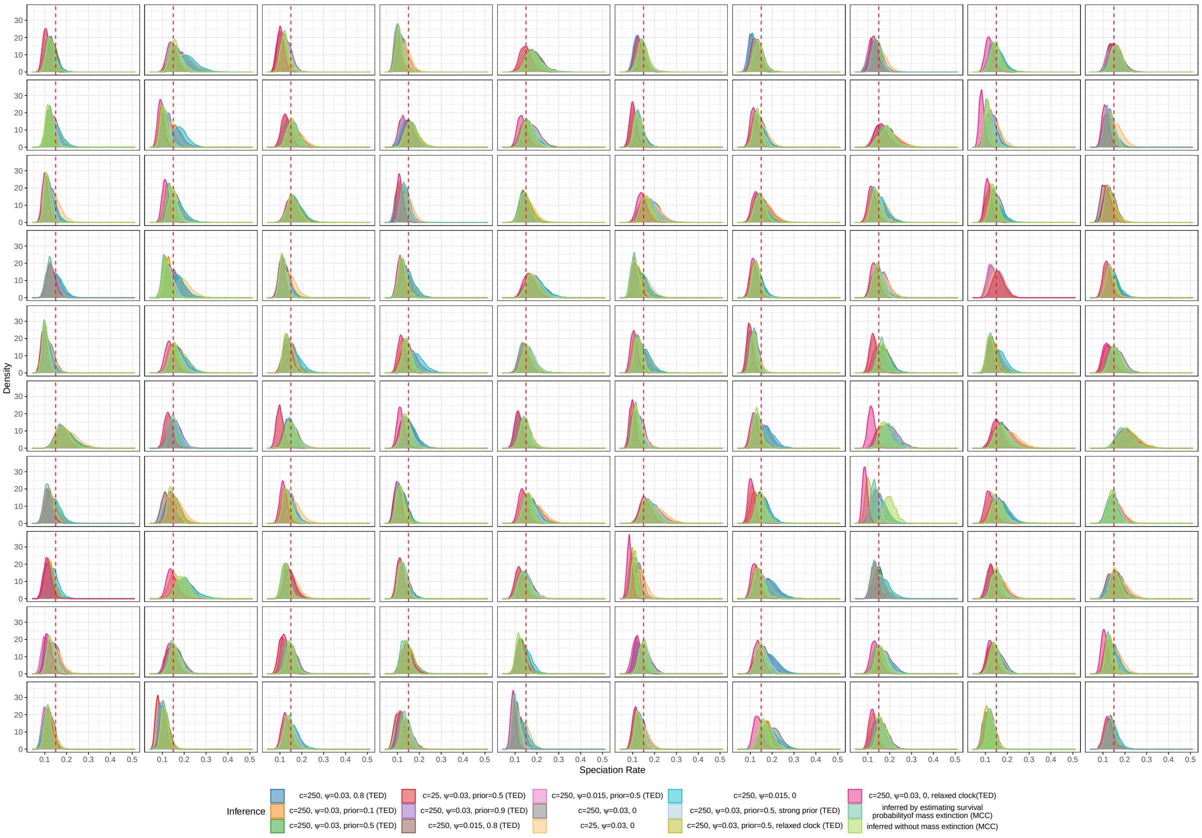


Figure S20: Posterior probability density plots of speciation rate for each simulated trees, obtained using different methods. For details on how the probability density curves were generated, please refer to Figure 2. The red vertical line indicates the true speciation rate (0.15). For the definitions of “TED”, “MCC”, “$c$”, “$\psi$”, “prior”, and “strong prior”, please refer to Figure 3. For the definitions of “inferred by estimating survival probability of mass extinction” and “inferred without mass extinction”, please refer to Figure 6.

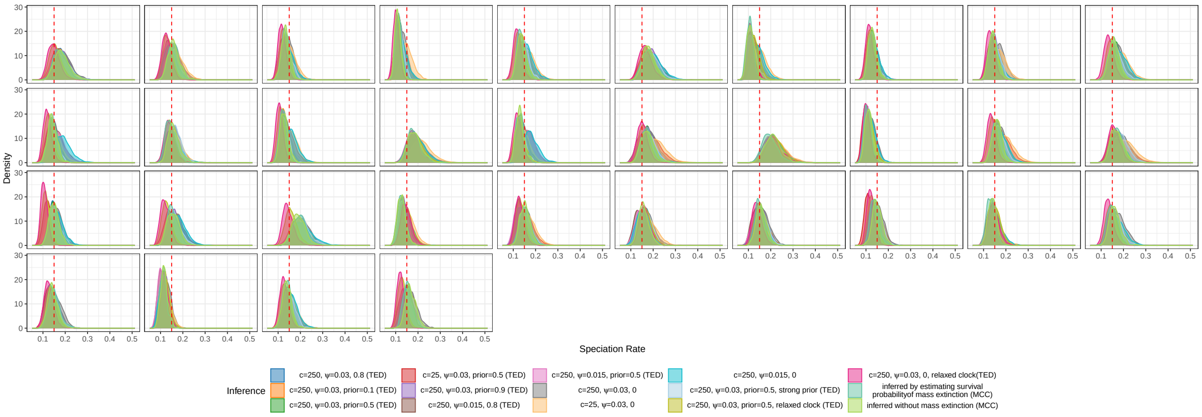


Figure S21: Posterior probability density plots of speciation rate for each simulated trees, obtained using different methods. Similar to Figure S20, but this figure includes only trees that converged across all methods and detected a single mass extinction event in the MCC tree. For the definitions of “$c$”, “$\psi$”, “prior”, and “strong prior”, please refer to Figure 1.

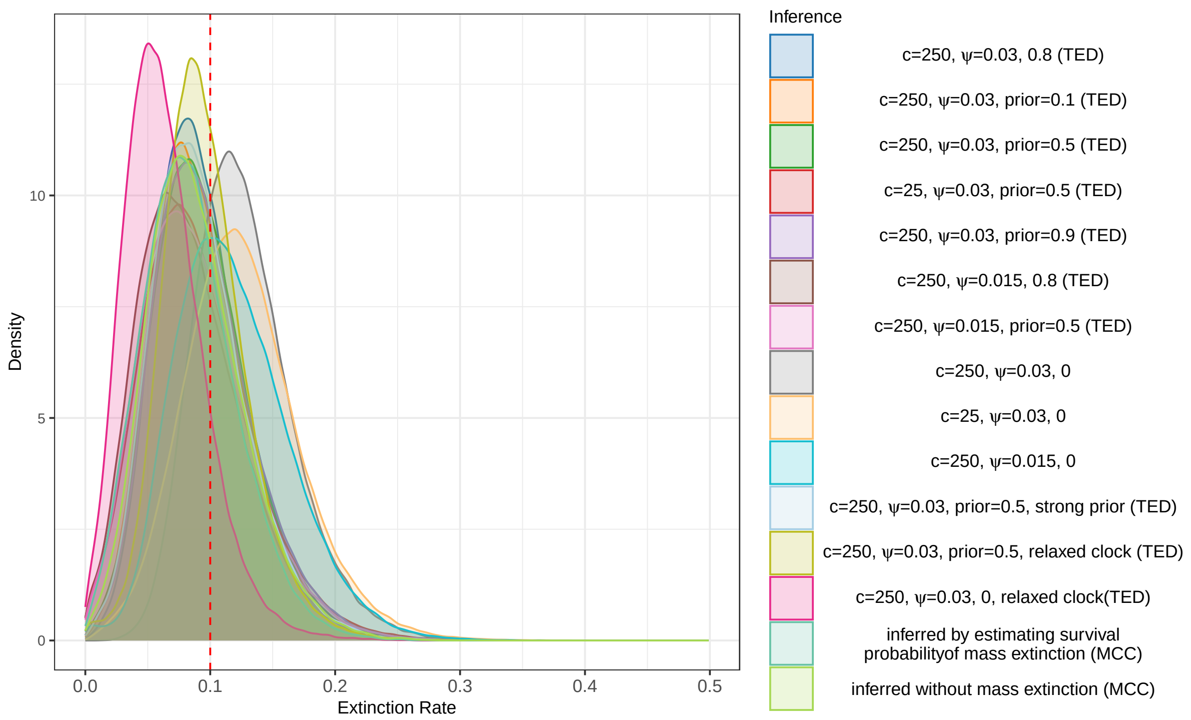


Figure S22: Posterior probability density plots of extinction rate for all simulated trees, obtained using different methods. For details on how the probability density curves were generated, please refer to Figure 2. The red vertical line indicates the true extinction rate (0.1). For the definitions of “TED”, “MCC”, “$c$”, “$\psi$”, “prior”, and “strong prior”, please refer to Figure 3. For the definitions of “inferred by estimating survival probability of mass extinction” and “inferred without mass extinction”, please refer to Figure 6.

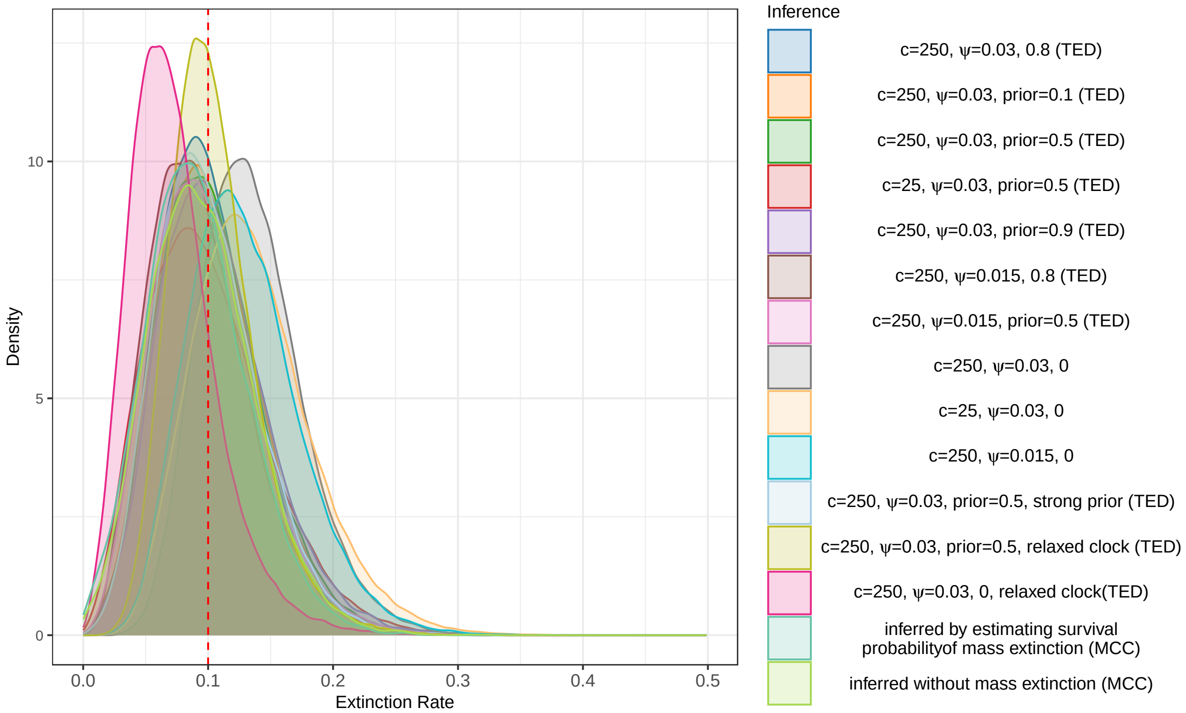


Figure S23: Posterior probability density plots of extinction rate for all simulated trees, obtained using different methods. Similar to Figure S22, but this figure includes only trees that converged across all methods and detected a single mass extinction event in the MCC tree.

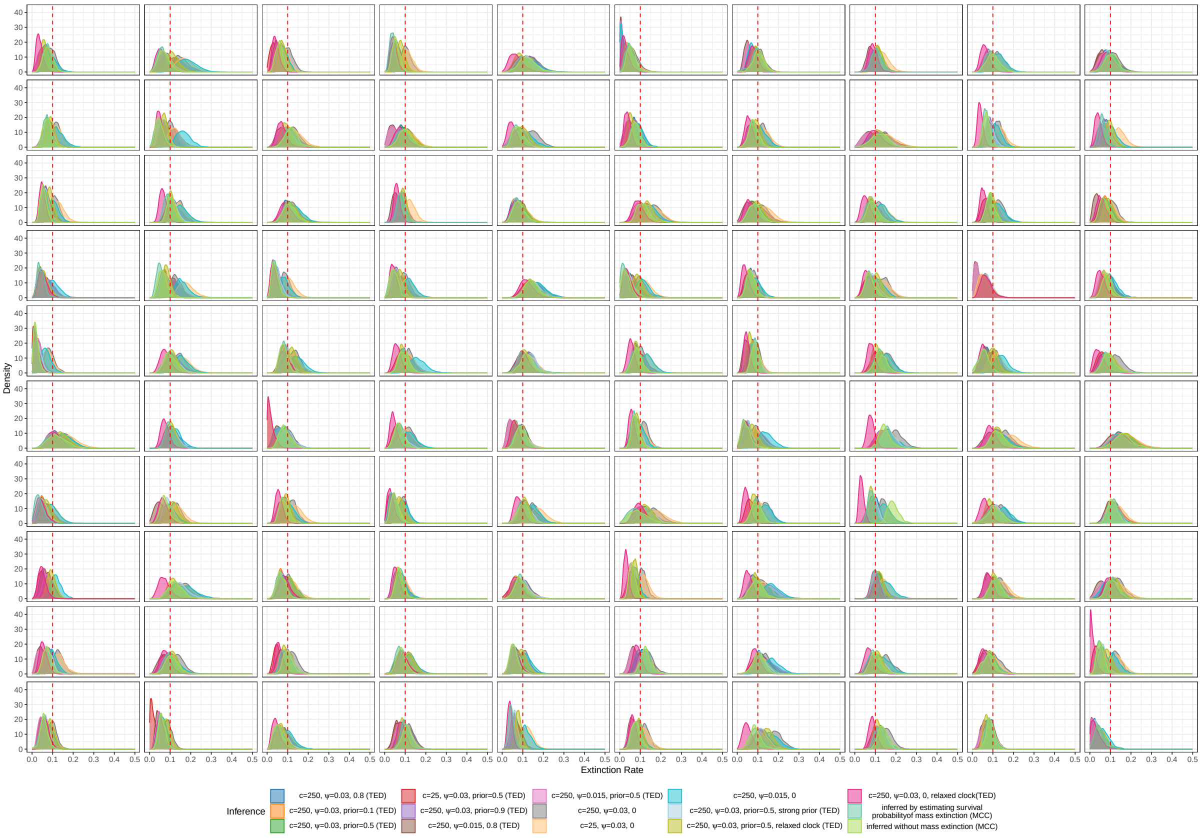


Figure S24: Posterior probability density plots of extinction rate for each simulated trees, obtained using different methods. For details on how the probability density curves were generated, please refer to Figure 2. The red vertical line indicates the true extinction rate (0.1). For the definitions of "TED", "MCC", "$c$", "$\psi$", "prior", and "strong prior", please refer to Figure 3. For the definitions of "inferred by estimating survival probability of mass extinction" and "inferred without mass extinction", please refer to Figure 6.

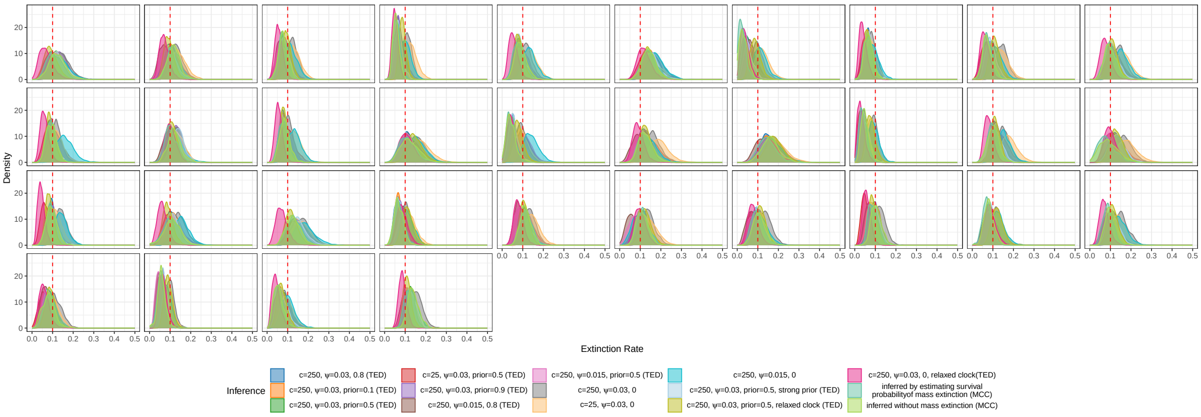


Figure S25: Posterior probability density plots of extinction rate for each simulated trees, obtained using different methods. Similar to Figure S24, but this figure includes only trees that converged across all methods and detected a single mass extinction event in the MCC tree. For the definitions of "$c$", "$\psi$", "prior", and "strong prior", please refer to Figure 1.

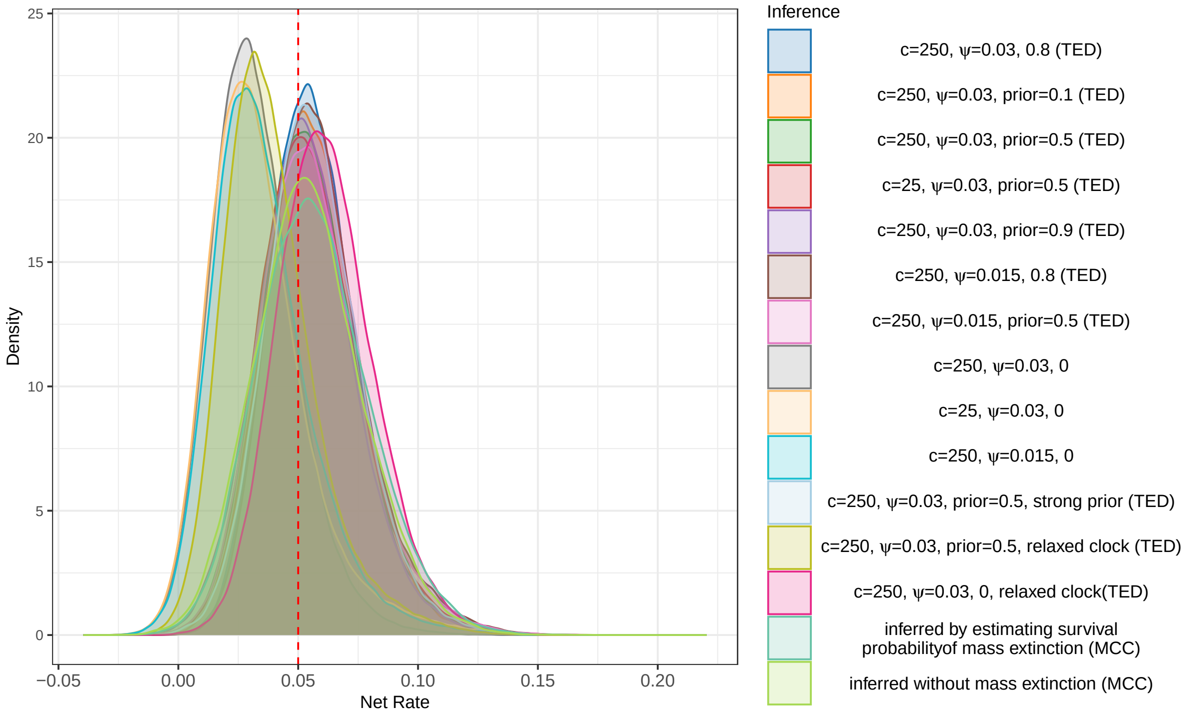


Figure S26: Posterior probability density plots of net rate for all simulated trees, obtained using different methods. For details on how the probability density curves were generated, please refer to Figure 2. The red vertical line indicates the true net rate (0.05). For the definitions of "TED", "MCC", "$c$", "$\psi$", "prior", and "strong prior", please refer to Figure 3. For the definitions of "inferred by estimating survival probability of mass extinction" and "inferred without mass extinction", please refer to Figure 6.

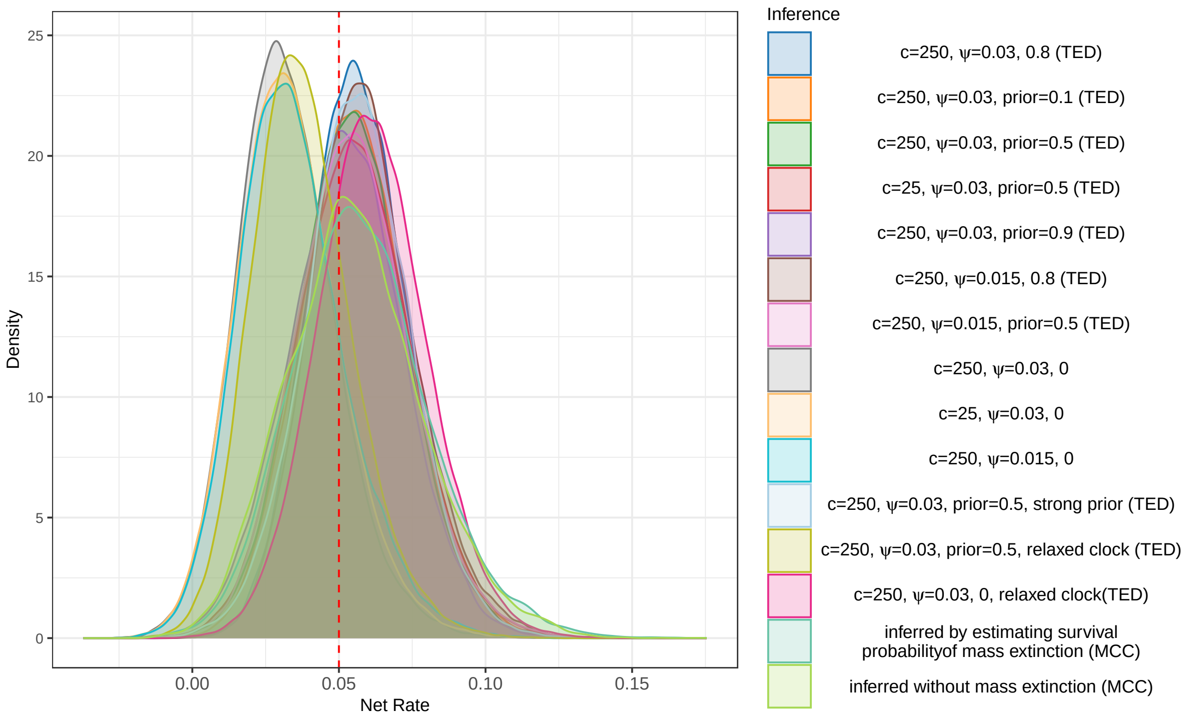


Figure S27: Posterior probability density plots of net rate for all simulated trees, obtained using different methods. Similar to Figure S26, but this figure includes only trees that converged across all methods and detected a single mass extinction event in the MCC tree.

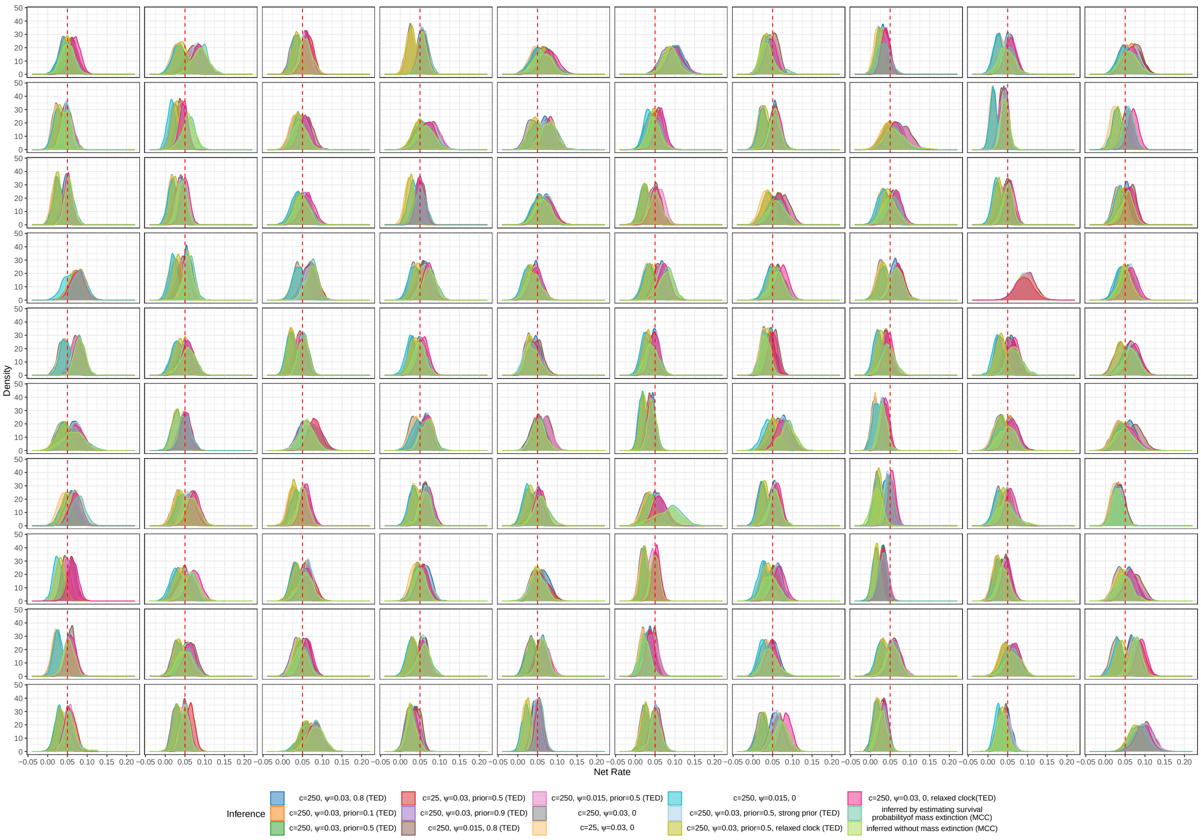


Figure S28: Posterior probability density plots of net rate for each simulated trees, obtained using different methods. For details on how the probability density curves were generated, please refer to Figure 2. The red vertical line indicates the true net rate (0.05). For the definitions of “TED”, “MCC”, “$c$”, “$\psi$”, “prior”, and “strong prior”, please refer to Figure 3. For the definitions of “inferred by estimating survival probability of mass extinction” and “inferred without mass extinction”, please refer to Figure 6.

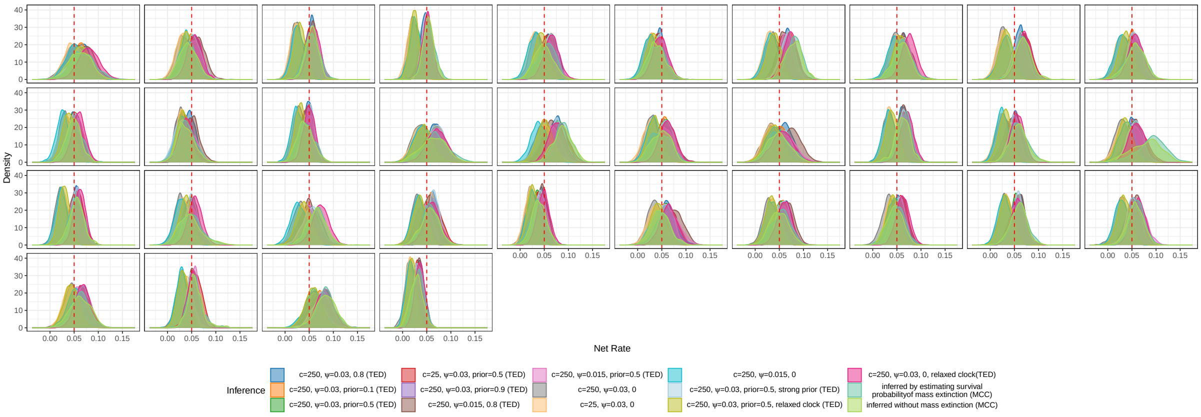


Figure S29: Posterior probability density plots of net rate for each simulated trees, obtained using different methods. Similar to Figure S28, but this figure includes only trees that converged across all methods and detected a single mass extinction event in the MCC tree. For the definitions of “$c$”, “$\psi$”, “prior”, and “strong prior”, please refer to Figure 1.

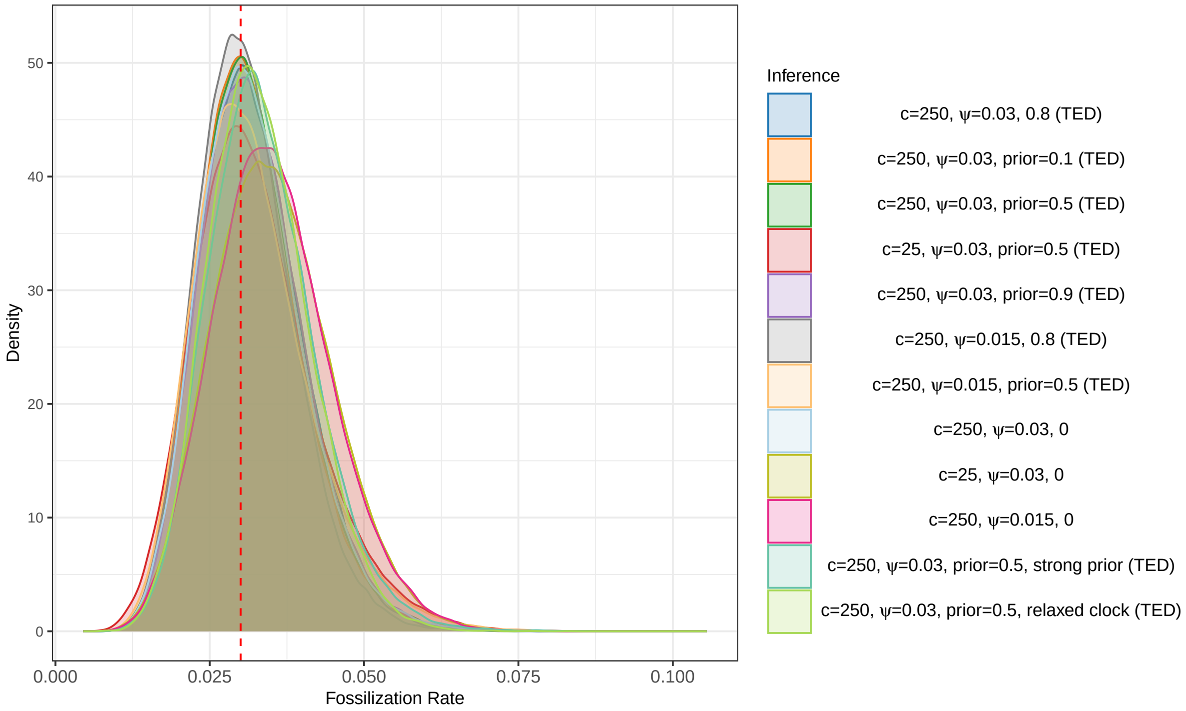


Figure S30: Posterior probability density plots of sampling rate for all simulated trees, simulated with sampling rate of 0.03, obtained using different methods. For details on how the probability density curves were generated, please refer to Figure 2. The red vertical line indicates the true sampling rate (0.03). For the definitions of "TED", "MCC", "$c$", "$\psi$", "prior", and "strong prior", please refer to Figure 3. For the definitions of "inferred by estimating survival probability of mass extinction" and "inferred without mass extinction", please refer to Figure 6.

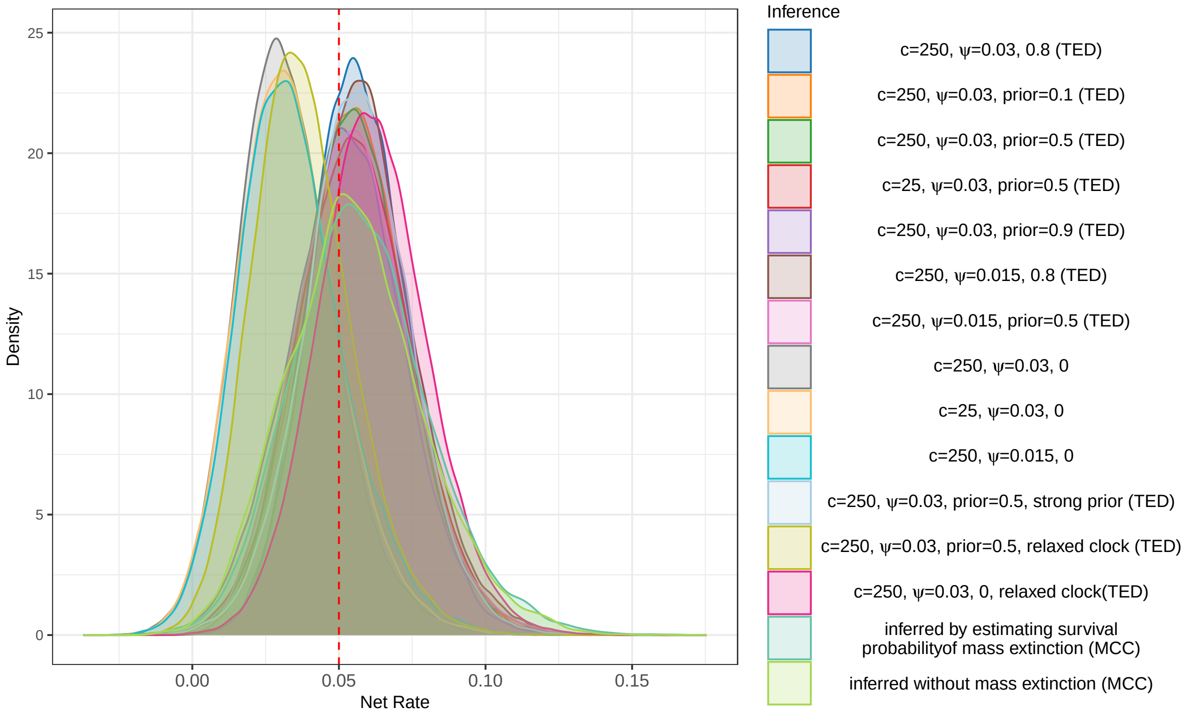


Figure S31: Posterior probability density plots of sampling rate for all simulated trees, simulated with sampling rate of 0.03, obtained using different methods. Similar to Figure S30, but this figure includes only trees that converged across all methods and detected a single mass extinction event in the MCC tree.

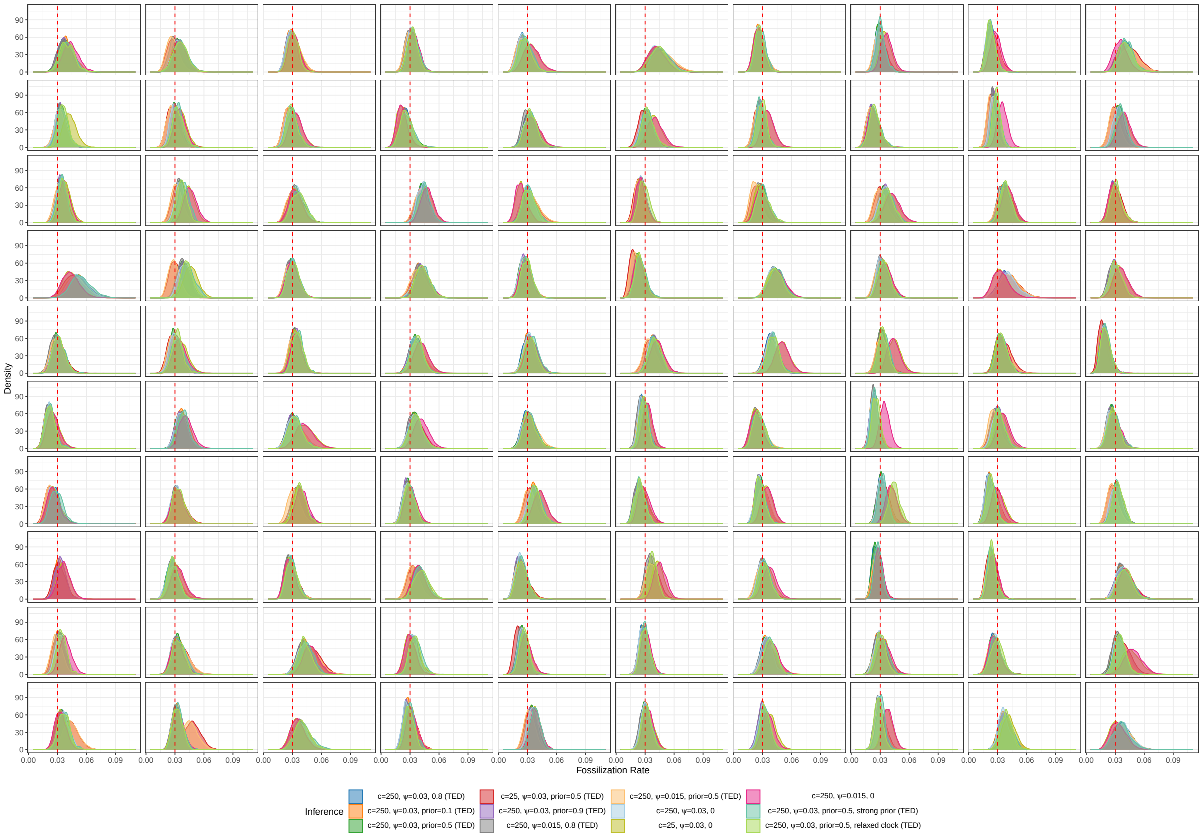


Figure S32: Posterior probability density plots of net rate for each simulated trees, simulated with sampling rate of 0.03, obtained using different methods. For details on how the probability density curves were generated, please refer to Figure 2. The red vertical line indicates the true sampling rate (0.03). For the definitions of "TED", "MCC", "$c$", "$\psi$", "prior", and "strong prior", please refer to Figure 3. For the definitions of "inferred by estimating survival probability of mass extinction" and "inferred without mass extinction", please refer to Figure 6.

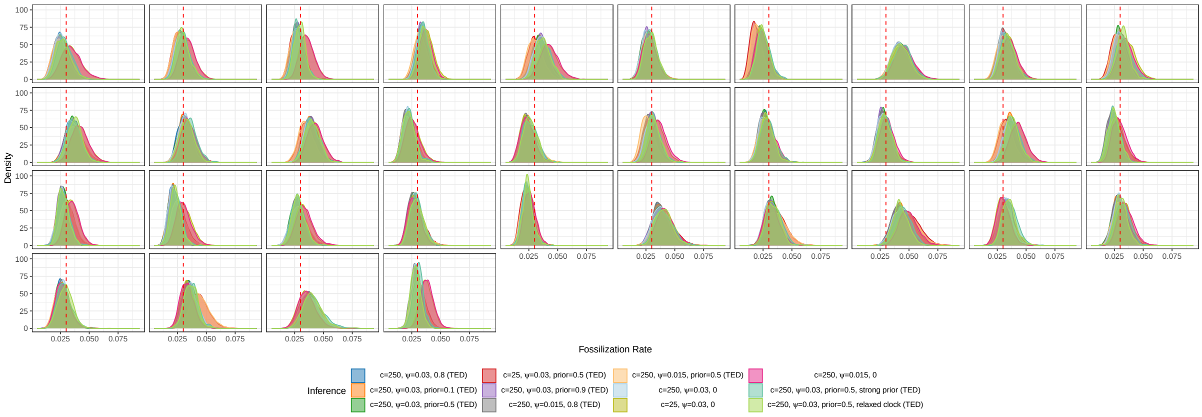


Figure S33: Posterior probability density plots of net rate for each simulated trees, simulated with sampling rate of 0.03, obtained using different methods. Similar to Figure S32, but this figure includes only trees that converged across all methods and detected a single mass extinction event in the MCC tree. For the definitions of "$c$", "$\psi$", "prior", and "strong prior", please refer to Figure 1.

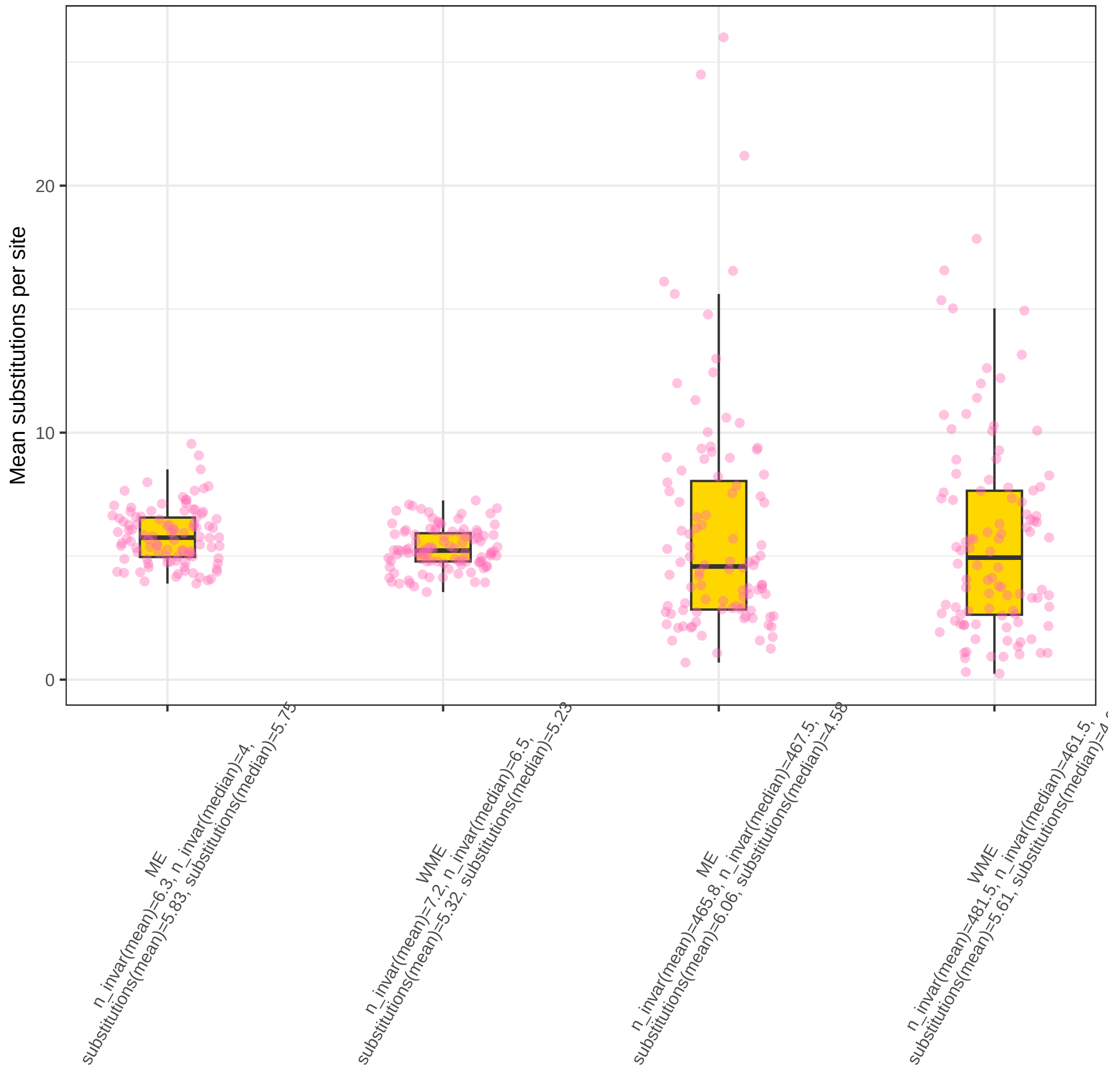


Figure S34: Distribution of the expected number of substitutions per site from 100 simulated trees. The box plots compare results from simulation with and without mass extinction, using both strict and UCNL relaxed clocks. The yellow box plot indicates the median (center line) and the interquartile range (the box), while each individual pink dot represents a single simulation run. The mean and median values for both the number of invariant characters (n_invar) and the substitutions per site (substitutions) are annotated below each corresponding condition.

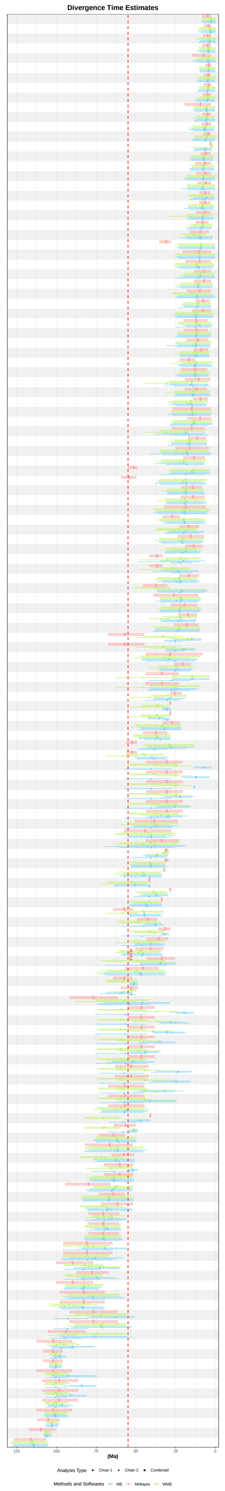


Figure S35: Tetraodontiform fishes divergence time estimates derived from MCC trees using various software and settings. For the RevBayes analyses, “WME” denotes the model without mass extinction (Model 1), and “ME” denotes the model with a mass extinction event (Model 3). Divergence times from the other analyses are plotted relative to the corresponding divergence time estimated from the RevBayes analysis without a mass extinction event (WME).

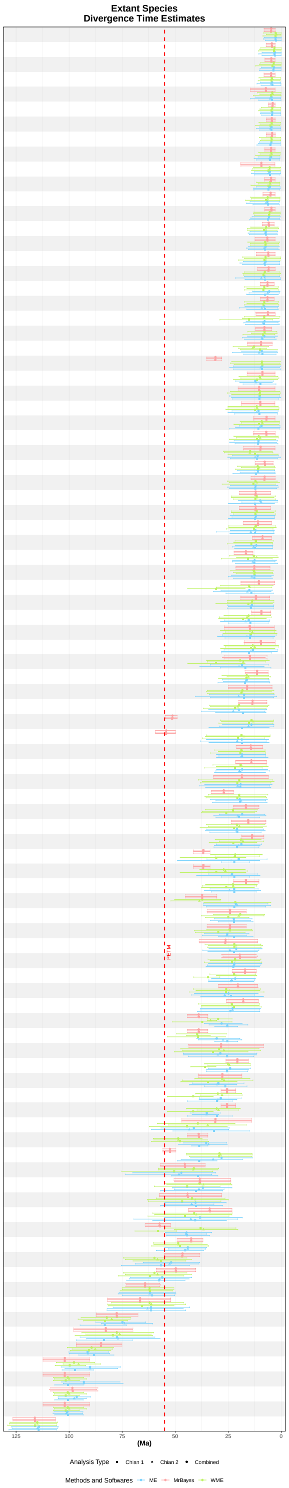


Figure S36: Tetraodontiform fishes (sampled ancestors excluded) divergence time estimates derived from MCC trees using various software and settings. For the RevBayes analyses, "WME" denotes the model without mass extinction (Model 1), and "ME" denotes the model with a mass extinction event (Model 3). Divergence times from the other analyses are plotted relative to the corresponding divergence time estimated from the RevBayes analysis without a mass extinction event (WME).

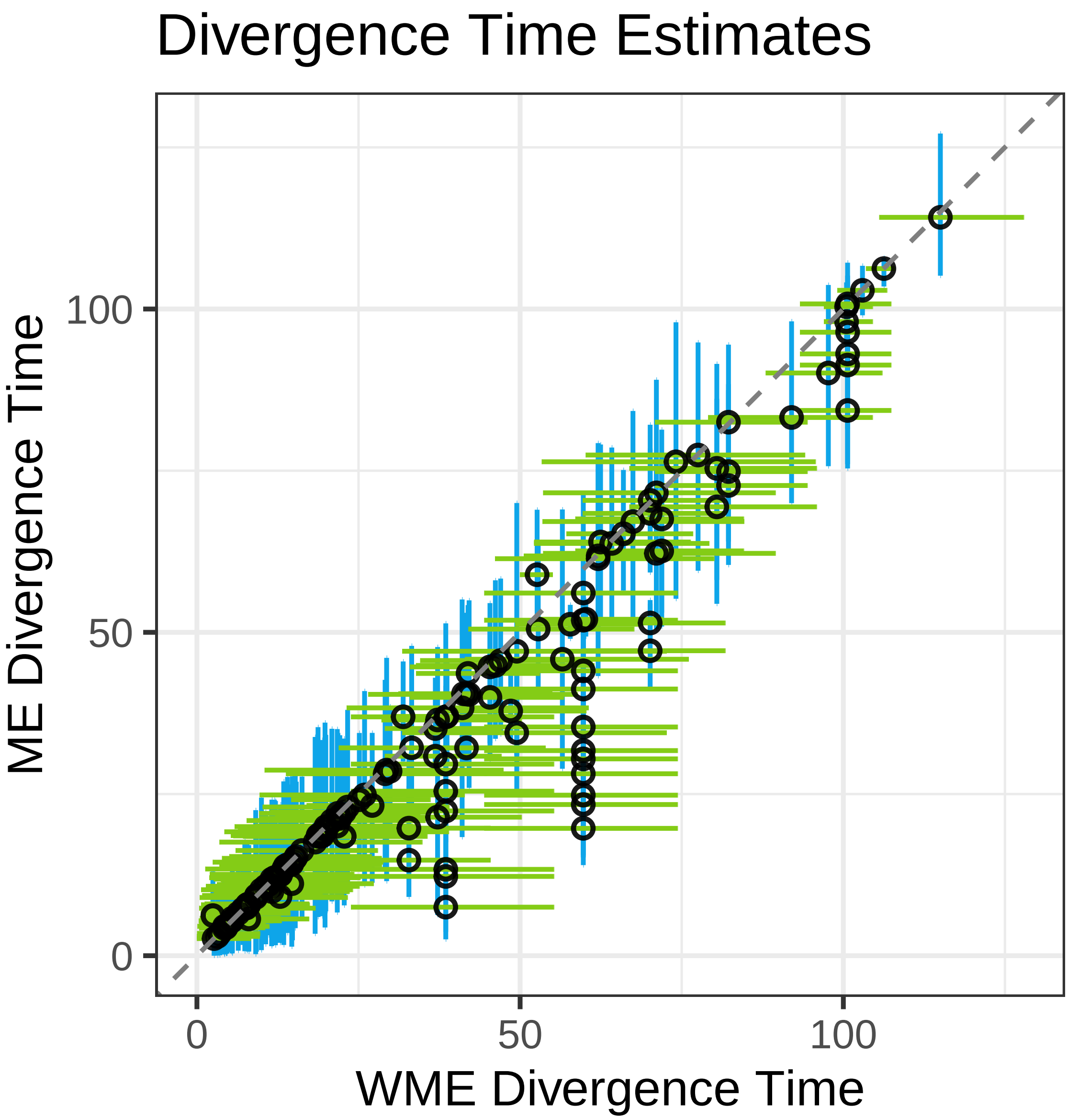


Figure S37: Scatter plot of tetraodontiform fish divergence time estimates from RevBayes, comparing results from a FBD model without a mass extinction (WME) against a model with a mass extinction (ME). The estimates were derived from the MCC trees of the combined MCMC chains for each analysis. The green and blue lines represent the 95% HPD intervals for the divergence times. Divergence times from the analysis with mass extinction (ME) are plotted relative to the corresponding divergence time estimated from the analysis without a mass extinction event (WME).

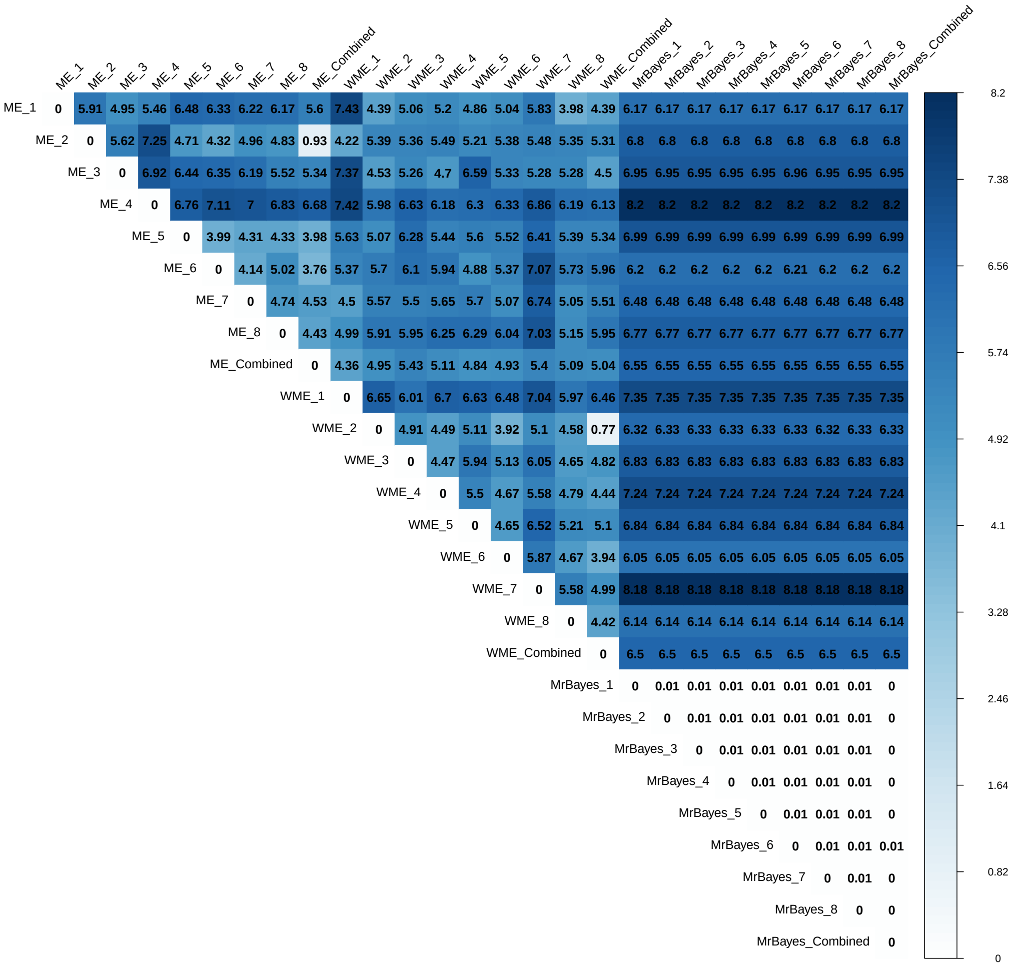


Figure S38: Comparison of divergence time differences among tetraodontiform fishes MCC trees generated using various software and settings. The figure presents pairwise differences in divergence time estimates between trees inferred from different software packages (RevBayes, MrBayes, BEAST2), separate MCMC chains (1 and 2), and combined MCMC chains. For the RevBayes analyses, "WME" denotes the model without mass extinction (Model 1), and "ME" denotes the model with a mass extinction event (Model 3). The divergence times are the average values, calculated using the divergence times from the two respective MCC trees as the reference.

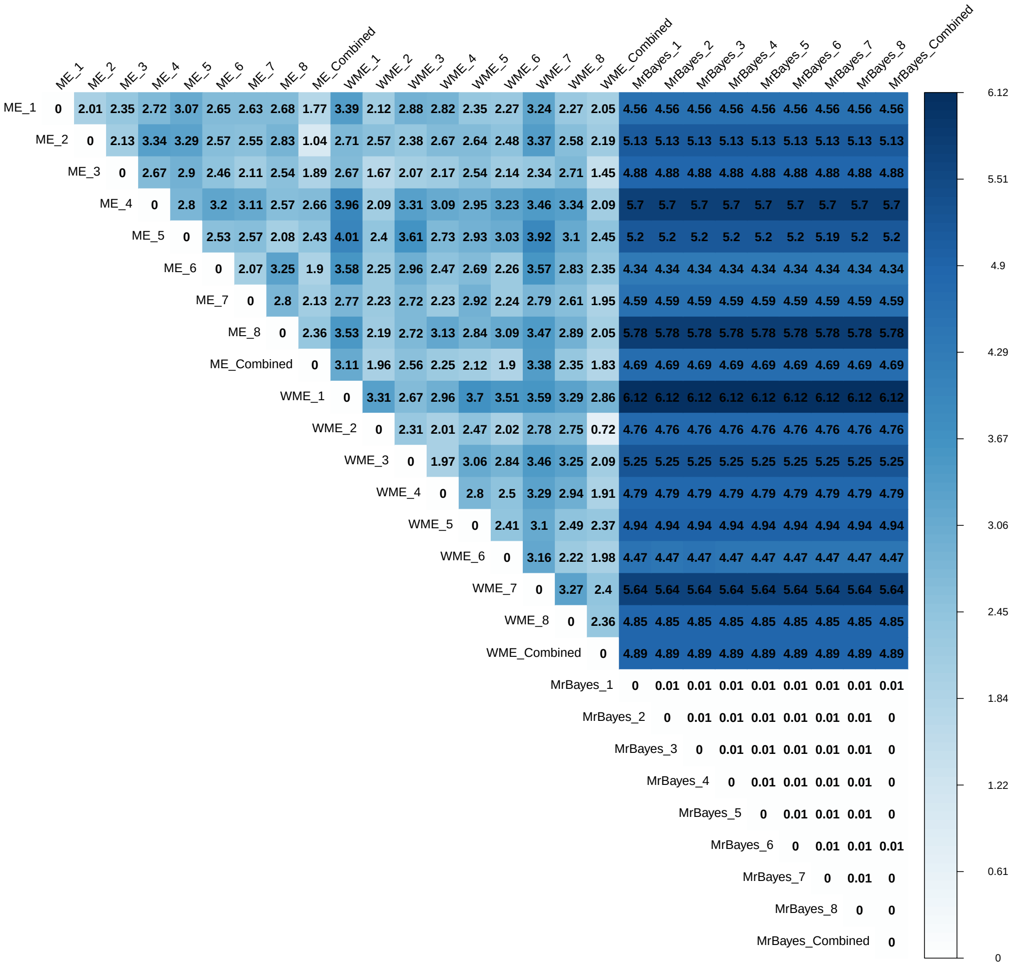


Figure S39: Comparison of fossil species (sampled ancestors excluded) divergence time differences among tetraodontiform fishes MCC trees generated using various software and settings. The figure presents pairwise differences in divergence time estimates between trees inferred from different software packages (RevBayes, MrBayes, BEAST2), separate MCMC chains (1 and 2), and combined MCMC chains. For the RevBayes analyses, "WME" denotes the model without mass extinction (Model 1), and "ME" denotes the model with a mass extinction event (Model 3). The divergence times are the average values, calculated using the divergence times from the two respective MCC trees as the reference.

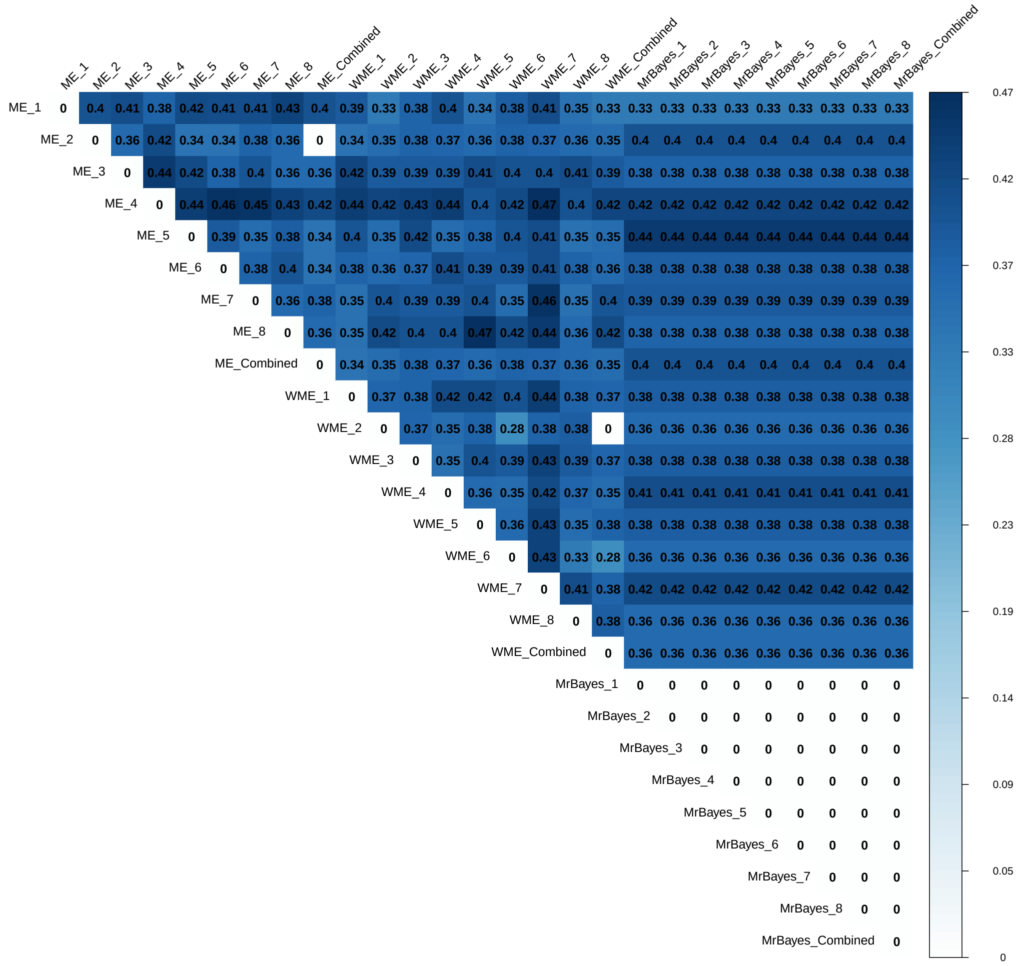


Figure S40: Comparison of RF distances among tetraodontiform fishes MCC trees generated using various software and settings. The figure presents pairwise differences in divergence time estimates between trees inferred from different software packages (RevBayes, MrBayes, BEAST2), separate MCMC chains (1 and 2), and combined MCMC chains. For the RevBayes analyses, "WME" denotes the model without mass extinction (Model 1), and "ME" denotes the model with a mass extinction event (Model 3).

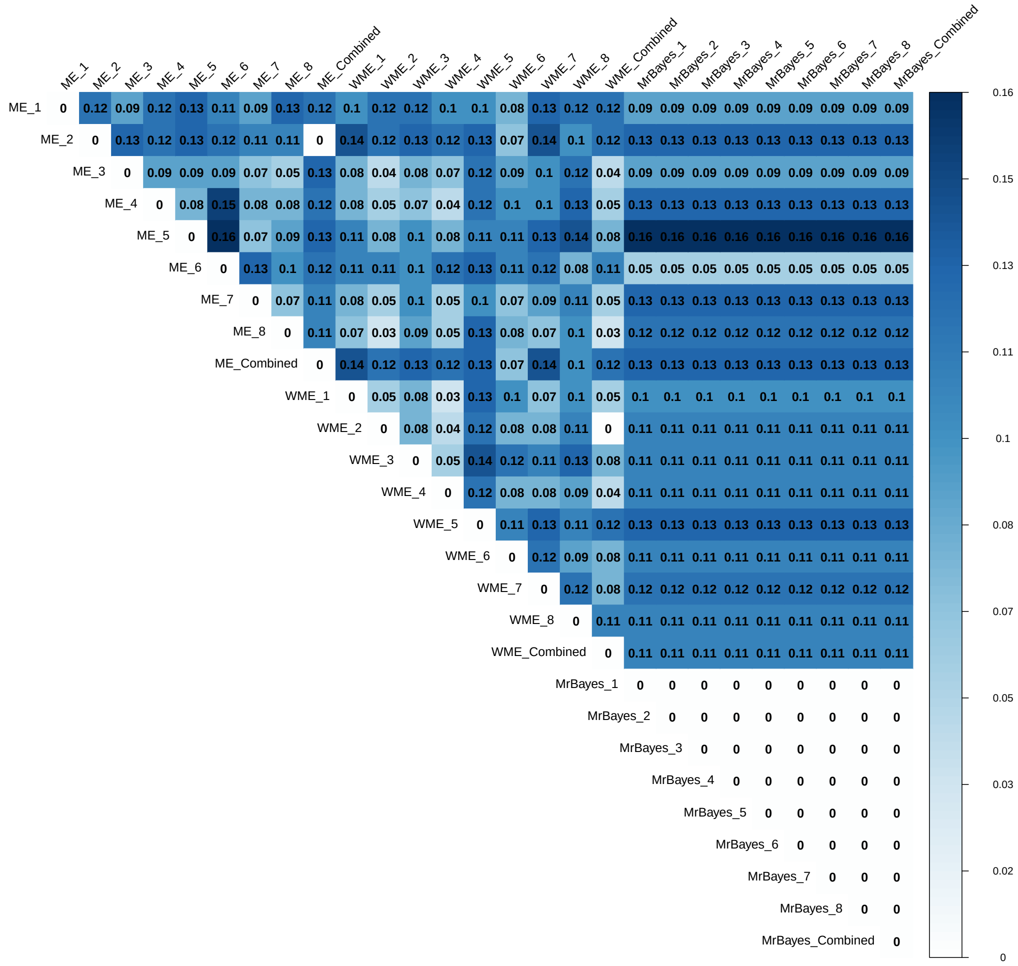


Figure S41: Comparison of RF distances differences among tetraodontiform fishes MCC trees generated using various software and settings. The figure presents pairwise differences in divergence time estimates between trees inferred from different software packages (RevBayes, MrBayes, BEAST2), separate MCMC chains (1 and 2), and combined MCMC chains. For the RevBayes analyses, "WME" denotes the model without mass extinction (Model 1), and "ME" denotes the model with a mass extinction event (Model 3).

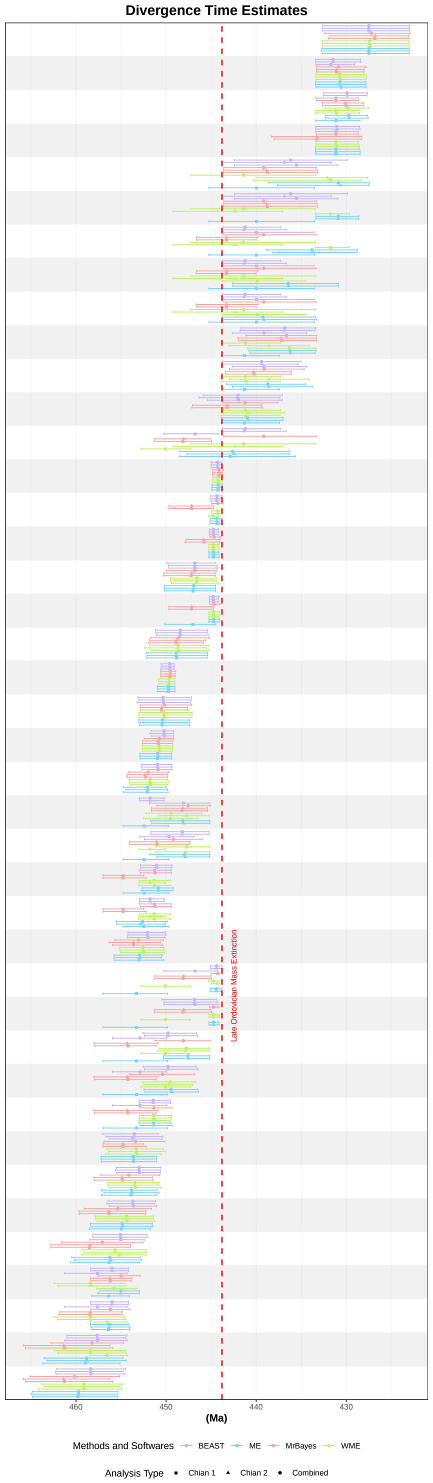


Figure S42: Crinoid divergence time estimates derived from MCC trees using various software and settings. For the RevBayes analyses, "WME" denotes the model without mass extinction (Model 1), and "ME" denotes the model with a mass extinction event (Model 3). Divergence times from the other analyses are plotted relative to the corresponding divergence time estimated from the RevBayes analysis without a mass extinction event (WME).

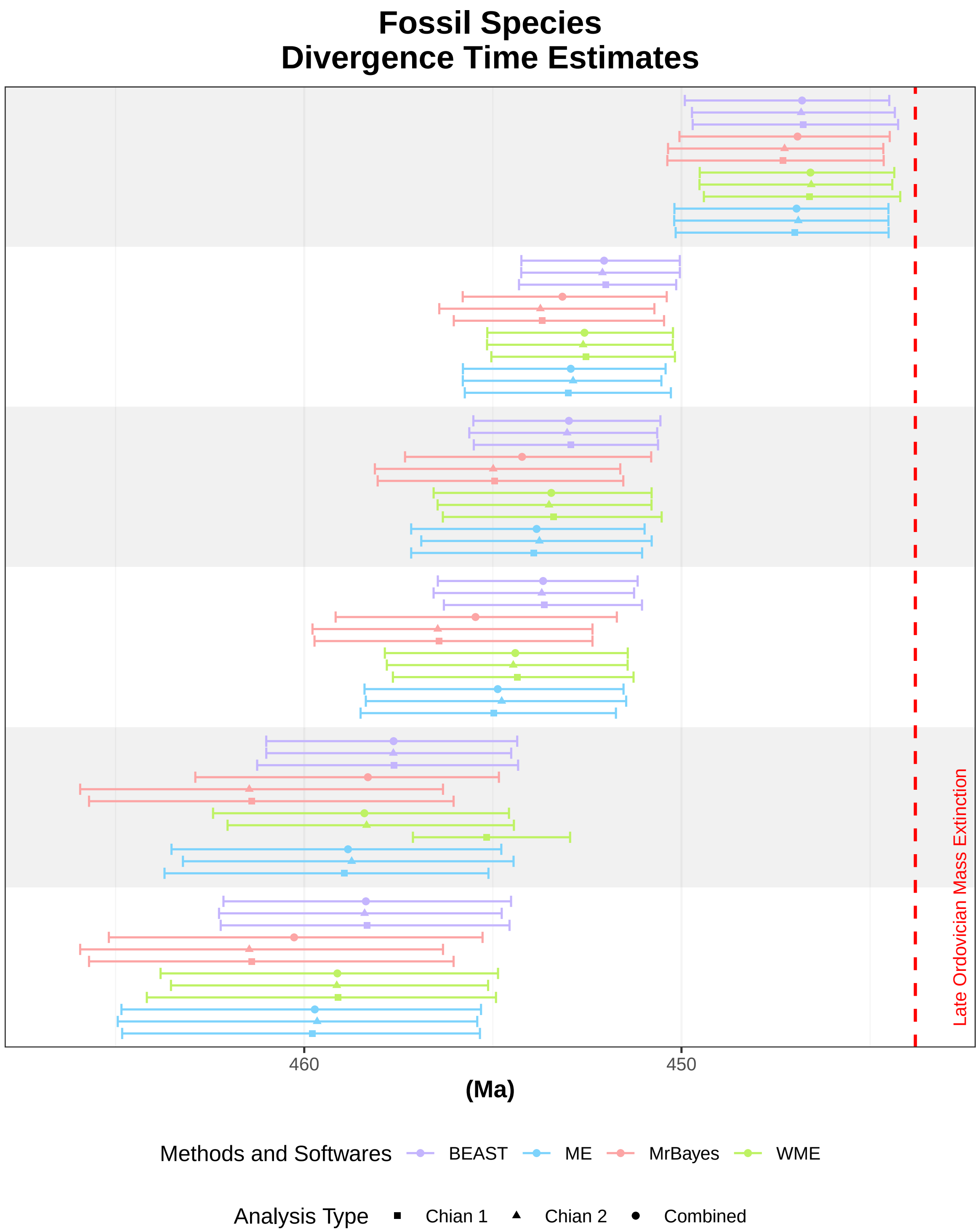


Figure S43: Crinoid fossil species (sampled ancestors excluded) divergence time estimates derived from MCC trees using various software and settings. For the RevBayes analyses, "WME" denotes the model without mass extinction (Model 1), and "ME" denotes the model with a mass extinction event (Model 3). Divergence times from the other analyses are plotted relative to the corresponding divergence time estimated from the RevBayes analysis without a mass extinction event (WME).

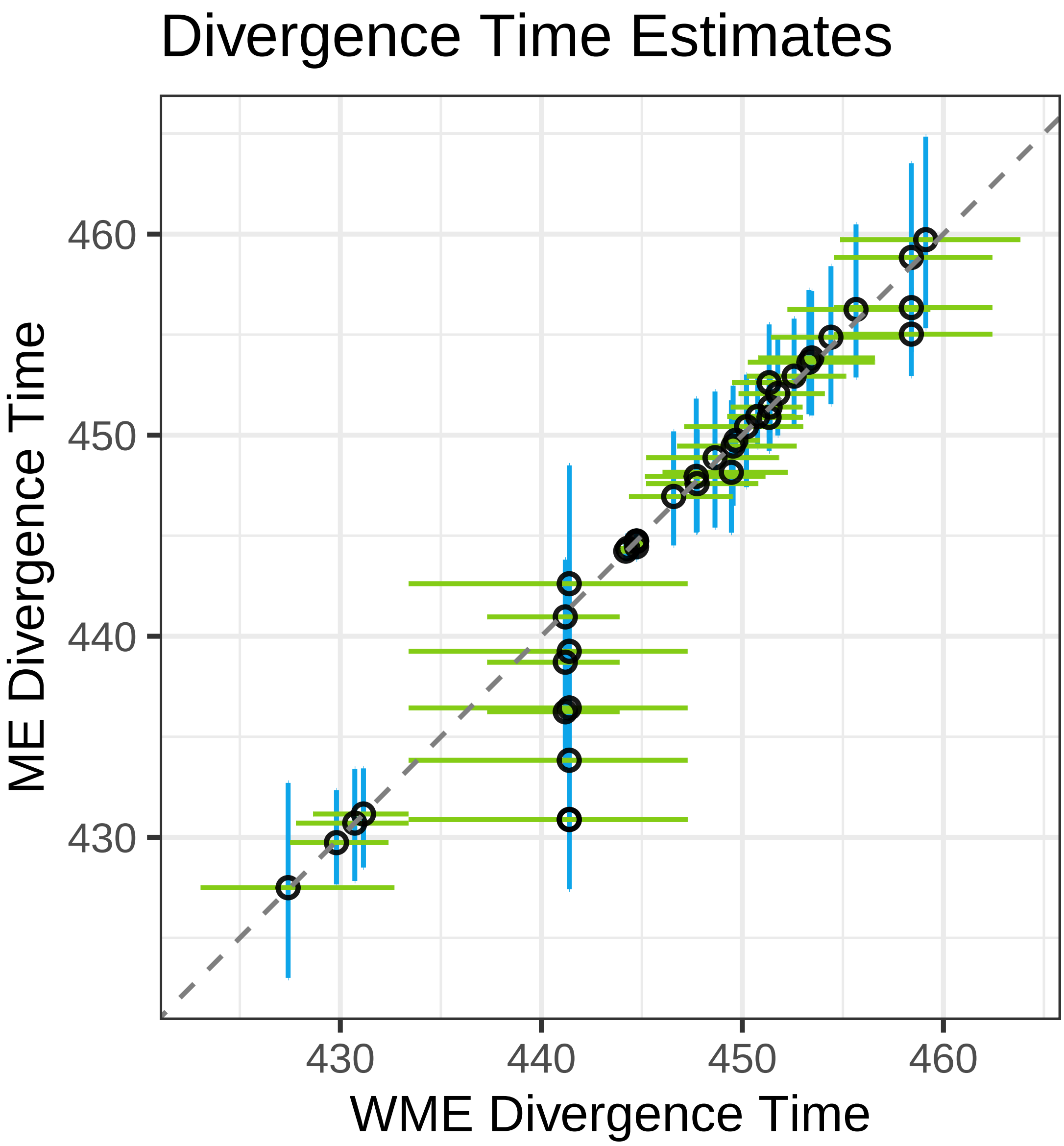


Figure S44: Scatter plot of crinoid divergence time estimates from RevBayes, comparing results from a FBD model without a mass extinction (WME) against a model with a mass extinction (ME). The estimates were derived from the MCC trees of the combined MCMC chains for each analysis. The green and blue lines represent the 95% HPD intervals for the divergence times. Divergence times from the analysis with mass extinction (ME) are plotted relative to the corresponding divergence time estimated from the analysis without a mass extinction event (WME).

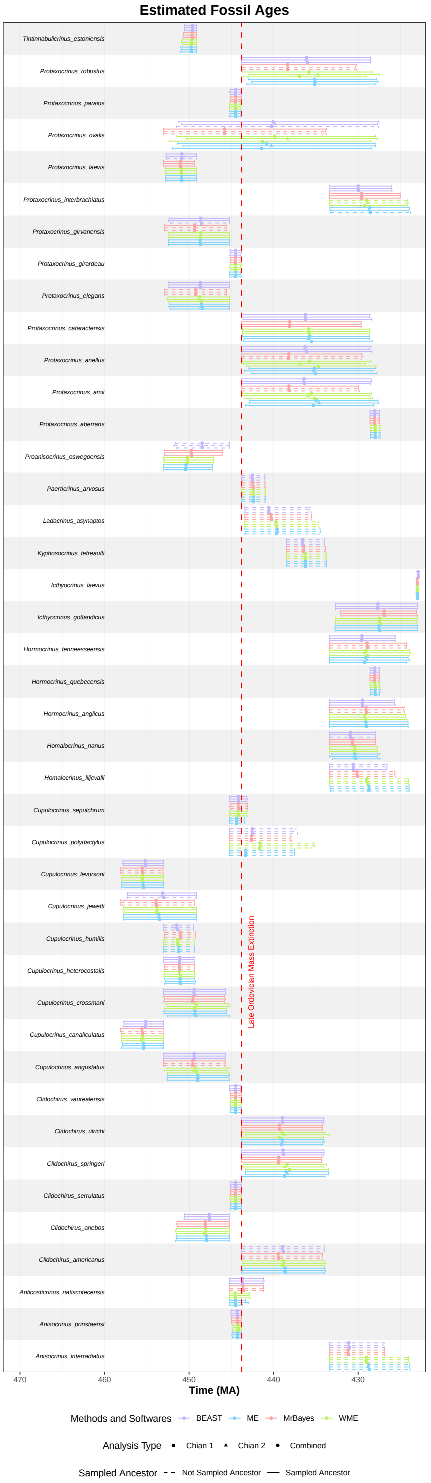


Figure S45: Estimated fossil ages for crinoids using various software and settings. The points represent the posterior mean of the fossil ages, while the bars show the 95% HPD intervals. For the RevBayes analyses, “WME” denotes the model without mass extinction (Model 1), and “ME” denotes the model with a mass extinction event (Model 3).

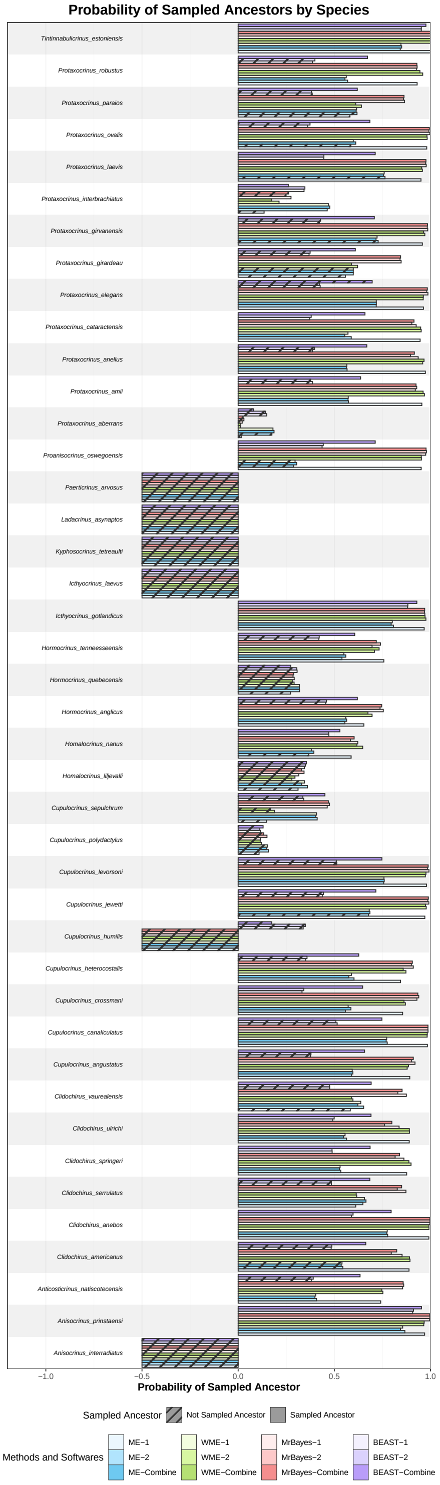


Figure S46: Posterior probability of each crinoid fossil being a sampled ancestor in the MCC using various software and settings. Fossils that are not sampled ancestors in their respective MCC tree are indicated with diagonal hash marks. For visual distinction, posterior probabilities of zero are plotted at a value of -0.5. For the RevBayes analyses, "WME" denotes the model without mass extinction (Model 1), and "ME" denotes the model with a mass extinction event (Model 3). Here, "1" and "2" denote different MCMC chains, and "combined" refers to the combined chains.

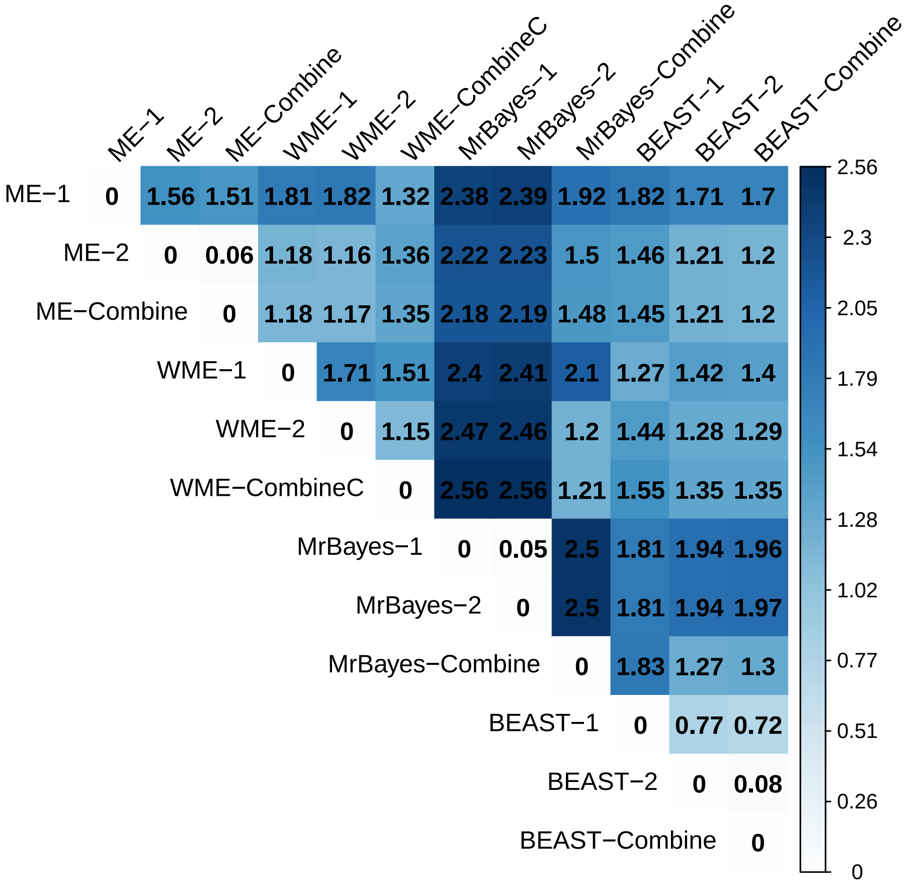


Figure S47: Comparison of divergence time differences among crinoid MCC trees generated using various software and settings. The figure presents pairwise differences in divergence time estimates between trees inferred from different software packages (RevBayes, MrBayes, BEAST2), separate MCMC chains (1 and 2), and combined MCMC chains. For the RevBayes analyses, "WME" denotes the model without mass extinction (Model 1), and "ME" denotes the model with a mass extinction event (Model 3). The divergence times are the average values, calculated using the divergence times from the two respective MCC trees as the reference.

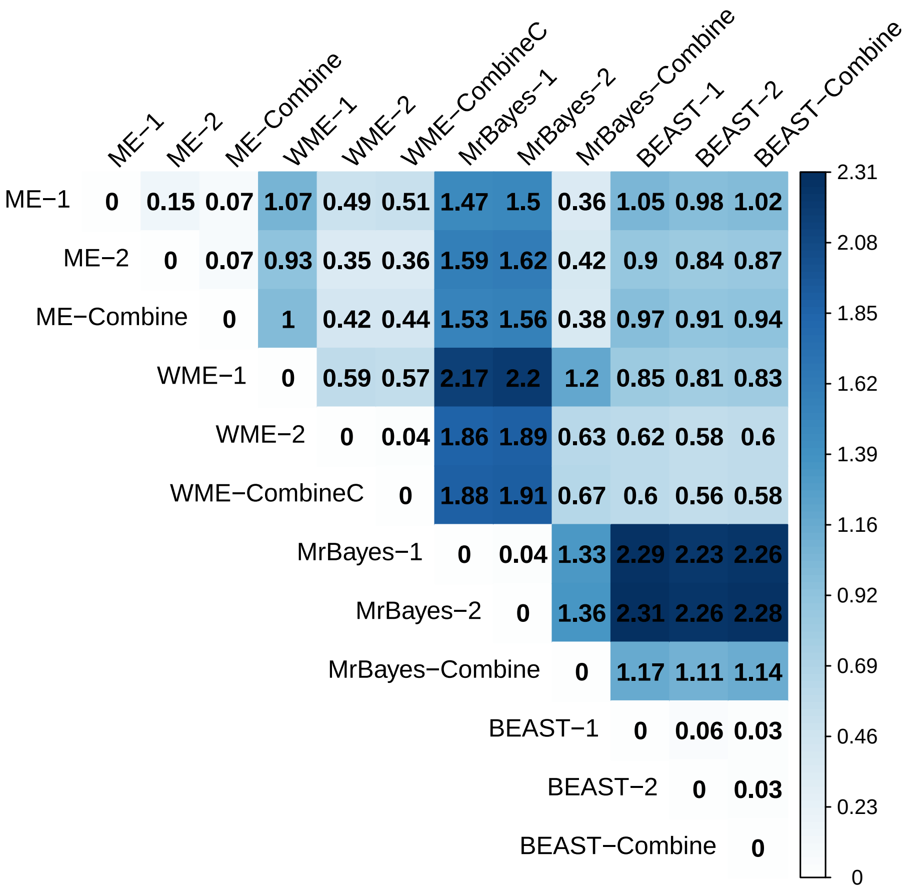


Figure S48: Comparison of fossil species (sampled ancestors excluded) divergence time differences among crinoid MCC trees generated using various software and settings. The figure presents pairwise differences in divergence time estimates between trees inferred from different software packages (RevBayes, MrBayes, BEAST2), separate MCMC chains (1 and 2), and combined MCMC chains. For the RevBayes analyses, "WME" denotes the model without mass extinction (Model 1), and "ME" denotes the model with a mass extinction event (Model 3). The divergence times are the average values, calculated using the divergence times from the two respective MCC trees as the reference.

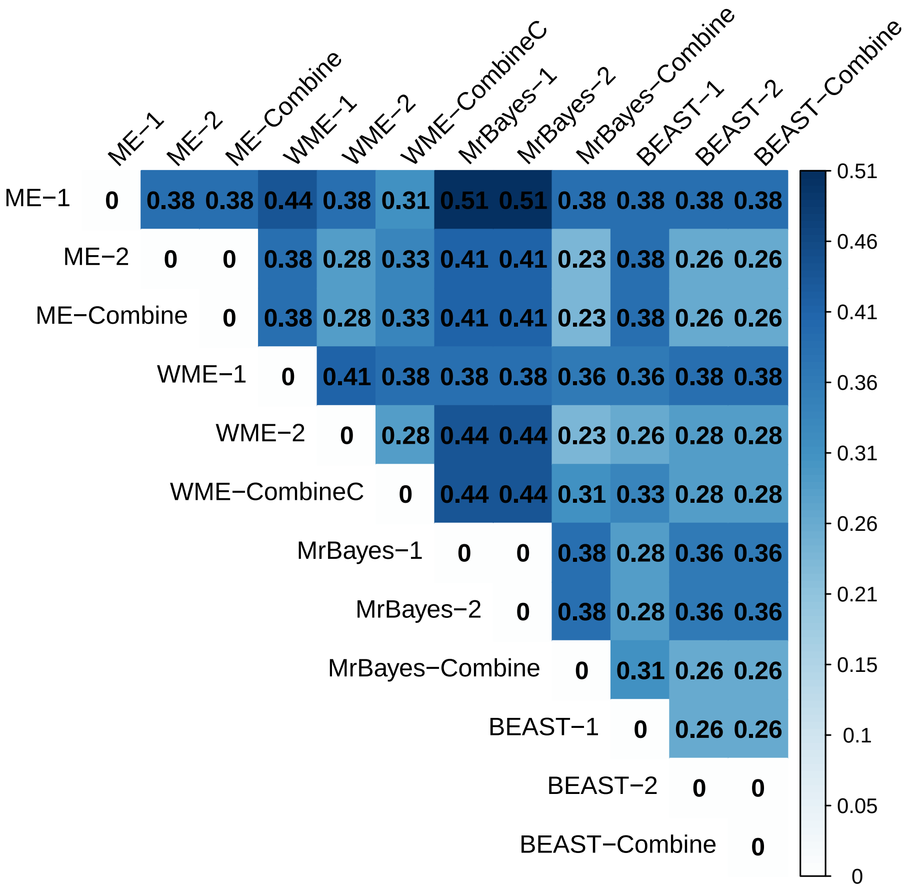


Figure S49: Comparison of RF distances among crinoid MCC trees generated using various software and settings. The figure presents pairwise differences in divergence time estimates between trees inferred from different software packages (RevBayes, MrBayes, BEAST2), separate MCMC chains (1 and 2), and combined MCMC chains. For the RevBayes analyses, “WME” denotes the model without mass extinction (Model 1), and “ME” denotes the model with a mass extinction event (Model 3).

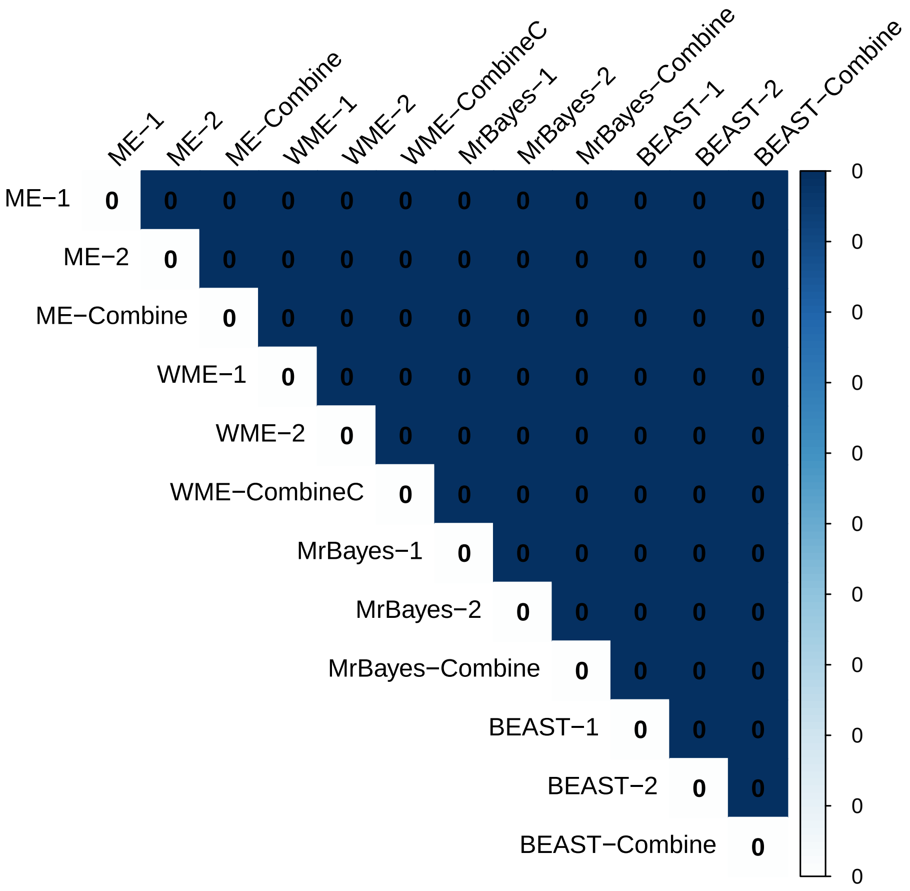


Figure S50: Comparison of RF distances differences among crinoid MCC fossil species (sampled ancestors excluded) trees generated using various software and settings. The figure presents pairwise differences in divergence time estimates between trees inferred from different software packages (RevBayes, MrBayes, BEAST2), separate MCMC chains (1 and 2), and combined MCMC chains. For the RevBayes analyses, "WME" denotes the model without mass extinction (Model 1), and "ME" denotes the model with a mass extinction event (Model 3).

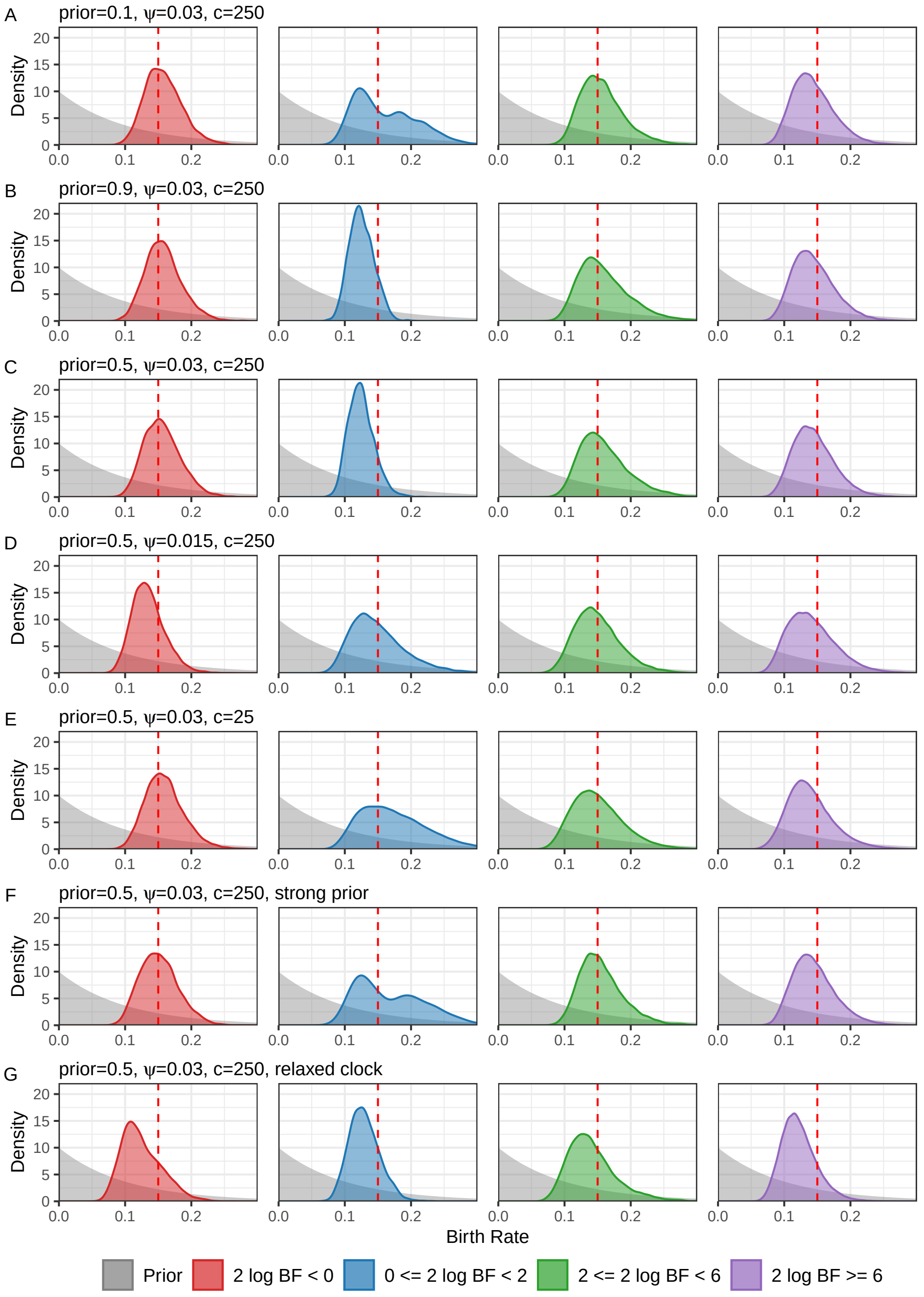


Figure S51: Posterior probability density plots of birth rates across all simulated trees, estimated using the FBD model with known mass extinction times and unknown mass extinction survival probabilities as tree priors in total-evidence dating. To ensure equal contributions from each tree when plotting the probability density curves for specific groups (e.g., $2 \log \mathrm{BF} < 0$ in panel A), MCMC sample sizes were normalized between replicates. Specifically, all post-burn-in MCMC chains within a group were randomly subsampled to the length of the shortest chain. The red vertical line indicates the true mass extinction survival probability (0.2). For the definitions of “ME”, “$c$”, “$\psi$”, “prior”, and “strong prior”, please refer to Figure 1.

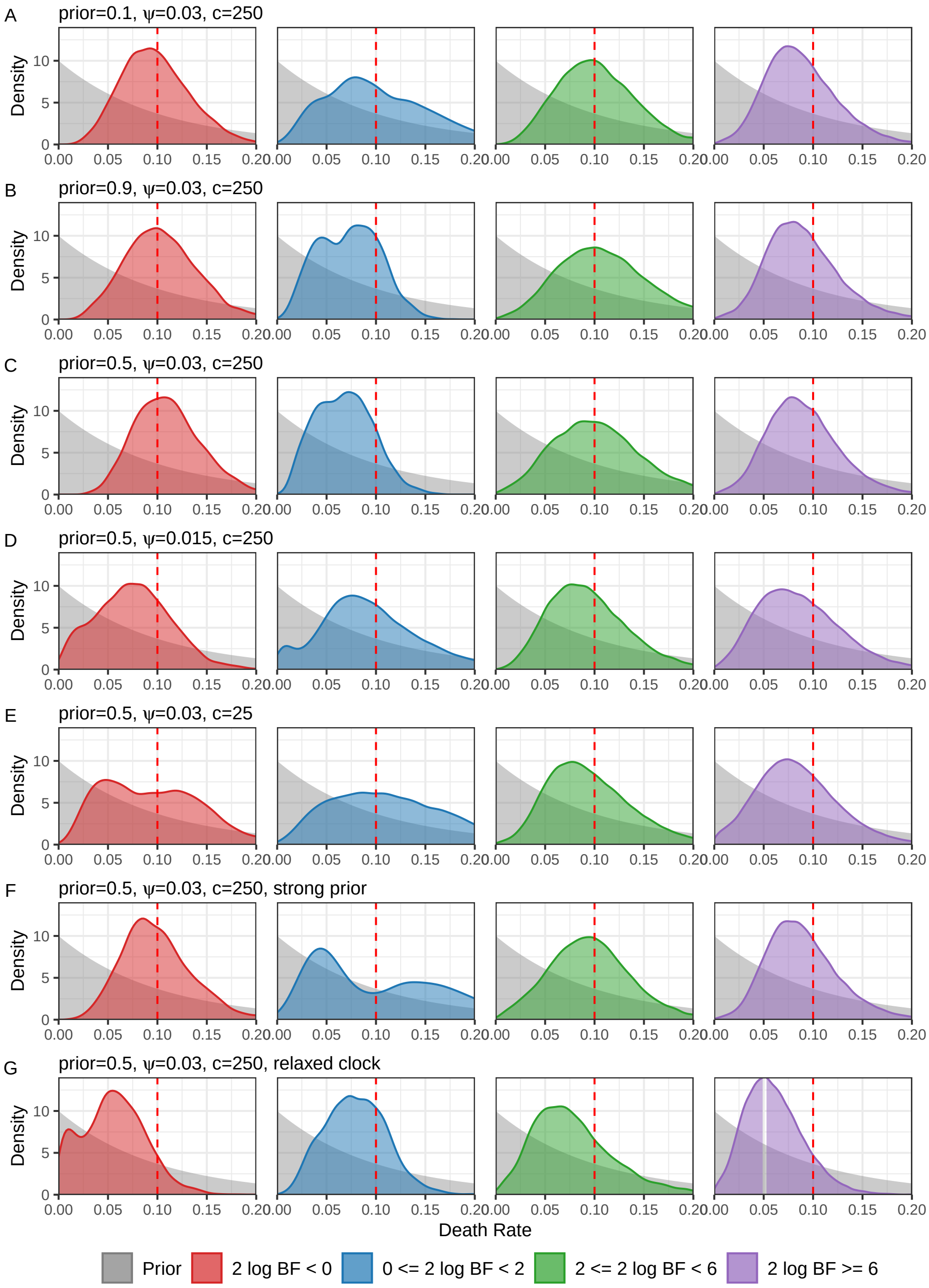


Figure S52: Posterior probability density plots of death rates across all simulated trees, estimated using the FBD model with known mass extinction times and unknown mass extinction survival probabilities as tree priors in total-evidence dating. To ensure equal contributions from each tree when plotting the probability density curves for specific groups (e.g., $2 \log \mathrm{BF} < 0$ in panel A), MCMC sample sizes were normalized between replicates. Specifically, all post-burn-in MCMC chains within a group were randomly subsampled to the length of the shortest chain. The red vertical line indicates the true mass extinction survival probability (0.2). For the definitions of "ME", "$c$", "$\psi$", "prior", and "strong prior", please refer to Figure 1.

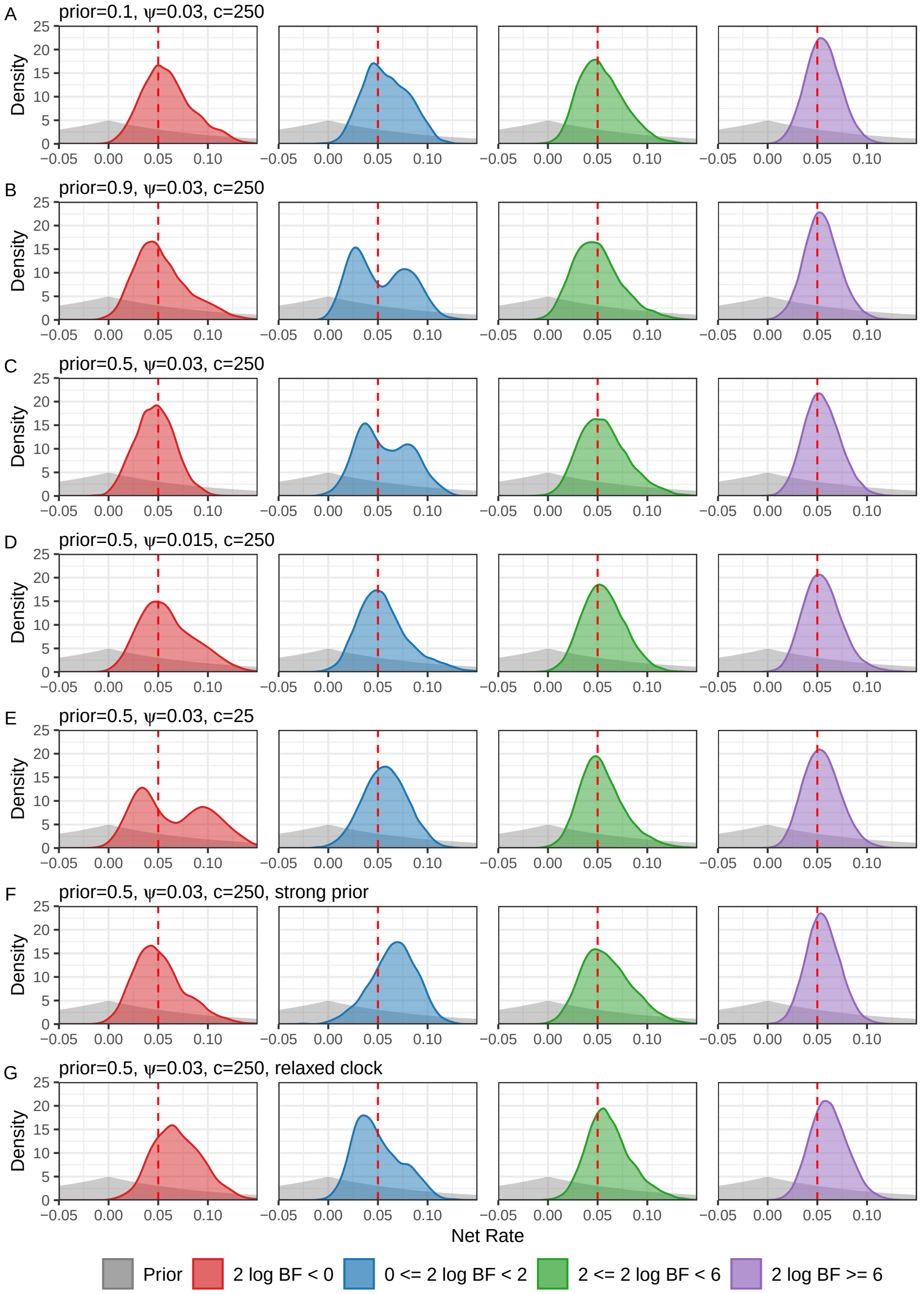


Figure S53: Posterior probability density plots of net rates across all simulated trees, estimated using the FBD model with known mass extinction times and unknown mass extinction survival probabilities as tree priors in total-evidence dating. To ensure equal contributions from each tree when plotting the probability density curves for specific groups (e.g., $2 \log \mathrm{BF} < 0$ in panel A), MCMC sample sizes were normalized between replicates. Specifically, all post-burn-in MCMC chains within a group were randomly subsampled to the length of the shortest chain. The red vertical line indicates the true mass extinction survival probability (0.2). For the definitions of "ME", "$c$", "$\psi$", "prior", and "strong prior", please refer to Figure 1.

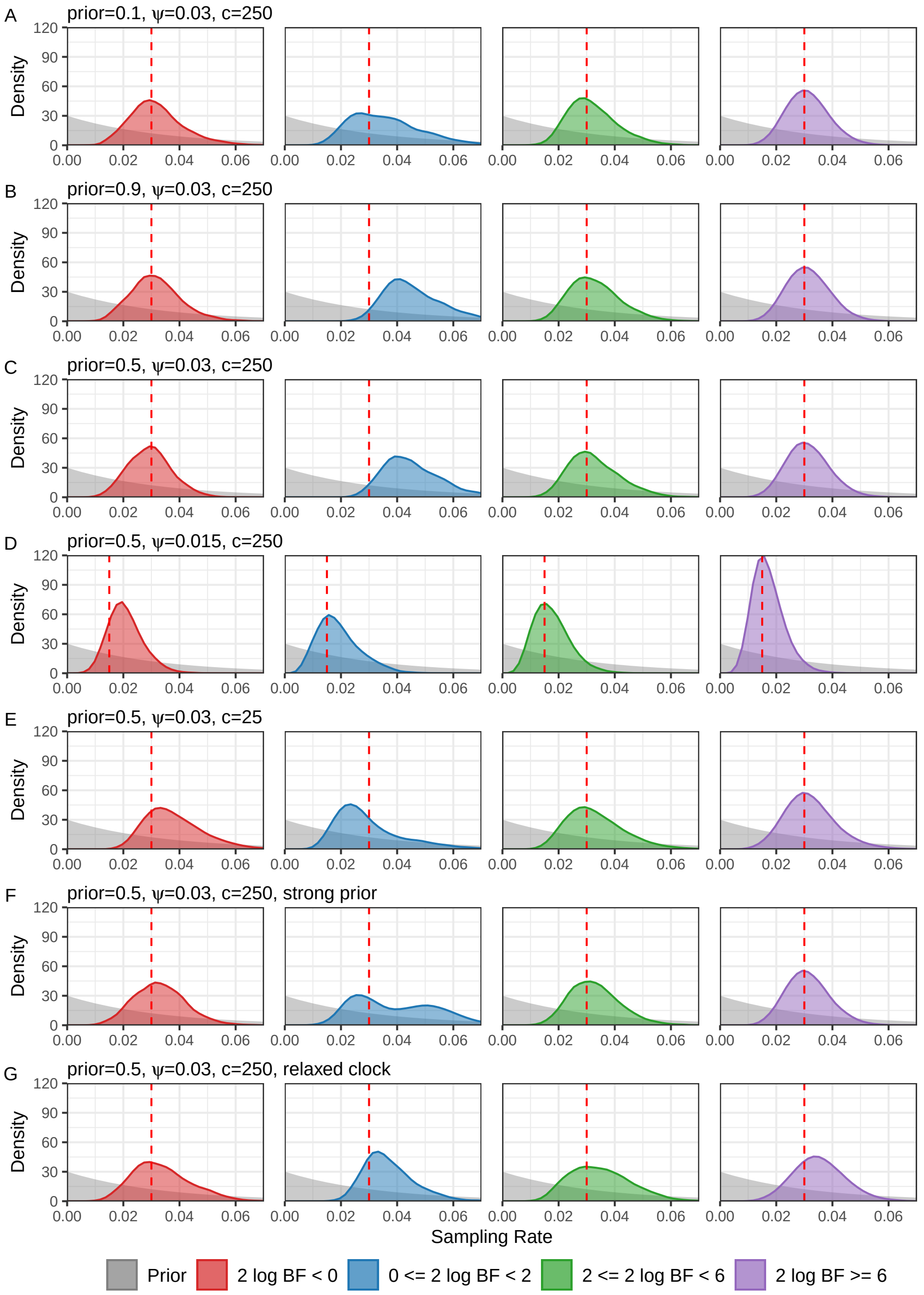


Figure S54: Posterior probability density plots of sampling rates across all simulated trees, estimated using the FBD model with known mass extinction times and unknown mass extinction survival probabilities as tree priors in total-evidence dating. To ensure equal contributions from each tree when plotting the probability density curves for specific groups (e.g., $2 \log \mathrm{BF} < 0$ in panel A), MCMC sample sizes were normalized between replicates. Specifically, all post-burn-in MCMC chains within a group were randomly subsampled to the length of the shortest chain. The red vertical line indicates the true mass extinction survival probability (0.2). For the definitions of "ME", "$c$", "$\psi$", "prior", and "strong prior", please refer to Figure 1.
